# Separation of cohorts on the basis of bacterial type IV conjugation systems identified from metagenomic assemblies

**DOI:** 10.1101/2021.04.15.440092

**Authors:** Benjamin R. Joris, Tyler S. Browne, Thomas A. Hamilton, David R. Edgell, Gregory B. Gloor

**Author notes:** Corresponding author: Gregory B. Gloor.

## Abstract

Conjugation enables the exchange of genetic elements throughout environments, including the human gut microbiome. Conjugative elements can carry and transfer clinically relevant metabolic pathways which makes precise identification of these systems in metagenomic samples clinically important. Here, we outline two distinct methods to identify conjugative systems in the human gut microbiome. We first show that conjugative systems exhibit strong population and age-level stratification. Additionally, we find that the total relative abundance of all conjugative systems present in a sample is not an informative metric to use, regardless of the method of identifying the systems. Finally, we demonstrate that the majority of assembled conjugative systems are not included within metagenomic bins, and that only a small proportion of the binned conjugative systems are included in “high-quality” metagenomic bins. Our findings highlight that conjugative systems differ between general North Americans and a cohort of North American pre-term infants, revealing a potential use as an age-related biomarker. Furthermore, conjugative systems can distinguish between other geographical-based cohorts. Our findings emphasize the need to identify and analyze conjugative systems outside of standard metagenomic binning pipelines.

**Importance:** The human gut microbiome is increasingly being associated with human health outcomes through shotgun metagenomic sequencing. The usual approach of metagenomic-level analyses is to bin assembled sequences into approximations of bacterial genomes and perform further investigations on the resultant bins. Here, we show that type IV conjugative systems differ between age and geographically-based cohorts and that these systems are systematically excluded by binning algorithms. We suggest that analysis of type IV conjugative systems should be added to the current metagenomic analysis approaches as they contain much information that could explain differences between cohorts beyond those we investigated.

## Background

Bacteria can acquire exogenous DNA through horizontal gene transfer. Conjugation is a common mechanism of horizontal gene transfer that relies on direct cell-cell contact to unidirectionally transfer DNA from a bacterial donor to a recipient cell. In bacteria, integrative conjugative elements (ICEs) and conjugative plasmids are mobilizable through the actions of type IV secretion systems (T4SS). Approximately half of the known plasmids are mobilizable in *trans* where the conjugative machinery is on a different genetic element than the transferred element, and the remainder are mobilizable in *cis* because the conjugative machinery is present on the same genetic element [1]. ICEs encode their own T4SS, and can mobilize other elements [2]. Conjugative elements often contain antibiotic resistance genes, but also can harbour useful biosynthetic and biodegradation genes [3]. Furthermore, conjugative systems can serve as vectors to introduce CRISPR systems, metabolic pathways or novel functions into the gut microbiota [4–9]. Therefore, characterizing the full complement of conjugative systems in the human gut could expand the number of useable vectors for these applications. Precise identification of conjugative systems from metagenomic samples could also provide insights to their distribution in populations and their correlation with antibiotic exposure, age, and health status.

For a DNA sequence to be considered mobilizable, it must encode an origin of transfer (*oriT*) sequence that is recognized and nicked by a relaxase protein [1, 10]. Relaxase proteins contain a conserved histidine triad that coordinates a divalent metal ion, as well as tyrosine residues that bind the oriT DNA sequence and catalyze the nicking reaction [11, 12]. In addition to a relaxase gene and an *oriT* sequence, a full complement of type IV secretion system and coupling proteins are required for a sequence to be mobilizable. In the well-studied *Agrobacterium tumefaciens* conjugative system, there are 12 proteins involved in the transfer of the DNA-relaxase complex from one bacterial cell to another [13, 14]. Homologs of the VirB4 ATPase that are essential for assembly of the conjugative system and DNA transfer are generally similar to the phylogeny of the bacteria harbouring them [15] and thus are useful for classifying conjugative systems [16]. The synteny of conjugative transfer genes is also highly conserved among conjugative systems [14]. Both the synteny and presence of highly-conserved genes involved in conjugation facilitates the classification of genetic elements as potentially conjugative if the sequences are annotated as belonging to the components of the T4SS [17] (Figure 1).

**Figure 1:**
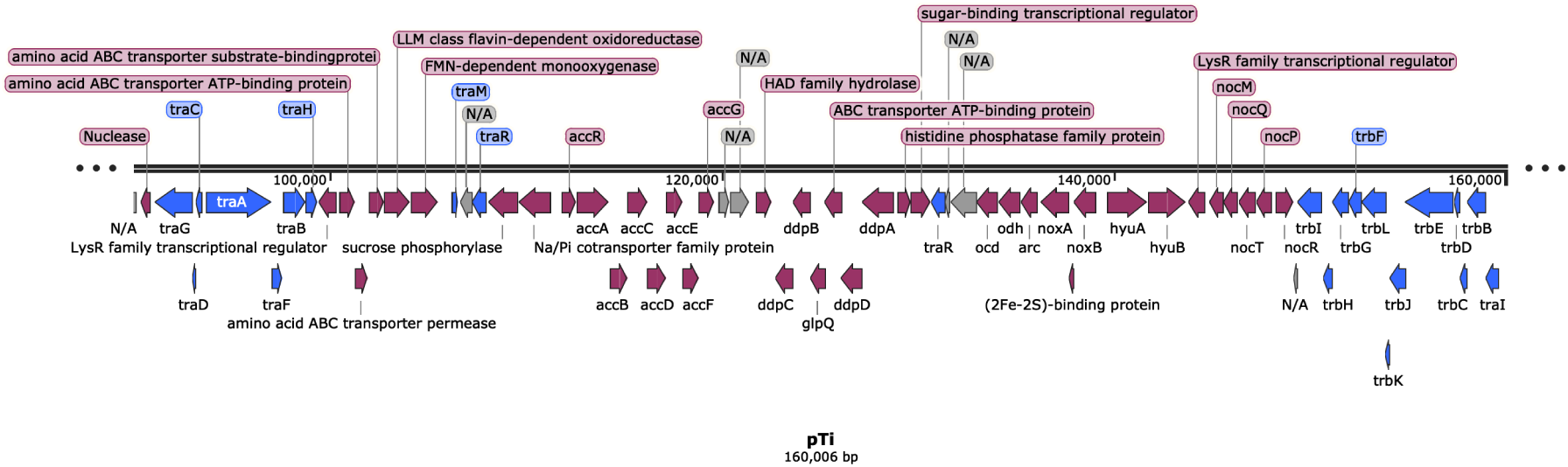
Example schematic of the gene organization of a bacterial conjugation system.

Previous work has identified novel conjugative systems in the human and animal gut microbiomes, but the focus was mainly on ICEs and not on conjugative plasmids [3, 18, 19]. Identifying conjugative plasmids from a short-read metagenomic assembly is difficult for several reasons. The initial barrier is the difficulty in assembling circularized plasmids from short-read sequencing data [20]. A second barrier is that the contiguous DNA sequences (contigs) that compose metagenome-assembled genomes (MAGs) are binned together based on sequence composition and coverage. Binning of a plasmid with its cognate genome will not happen unless the contigs that compose the plasmid are maintained in the same copy-number and have the same sequence composition as the chromosome. These criteria are generally not met because conjugative systems are usually more AT rich than the cognate chromosome [1] and often do not have a unit copy number. Since nearly 80% of the non-redundant set of genomes from the human-gut microbiome are from difficult-to-culture species that are known only from MAGs [21], alternate methods must be employed to assemble and identify conjugative plasmids from the metagenomic sequencing data. Computational tools have recently been developed to identify plasmids from metagenomic assemblies [22], but would be rendered pointless if applied to already binned data that systematically excludes plasmids [23]. Methods that identify conjugative systems prior to binning should be able to capture the full spectrum of ICEs and conjugative plasmids.

Here, we show that T4SS conjugative systems can be identified using two distinct methods (Figure 2). First, we used profile HMMs (pHMMs) to identify conjugative systems directly from metagenomic assemblies of general North American and North American pre-term infant samples [17]. Second, we searched predicted protein sequences vs. UniRef90 [24] for proteins involved in conjugation to identify conjugative systems from a human gut microbiome genome set. With this approach differences between additional cohorts in the relative abundances of extracted systems could be recognized. While the differences between cohorts found using the two methods were not identical, both methods did illustrate that different age and population cohorts were distinct. Non-equivalent data between the methods suggests some level of incompleteness or bias in the conjugative systems found by one or both methods. Finally, we demonstrate that the majority of conjugative systems produced by a metagenomic assembly are not included in high-quality bins that were used to compose human gut microbiome genome sets. Our findings provide a roadmap to integrate the analysis of conjugative systems alongside the genomic content of bacteria.

**Figure 2:**
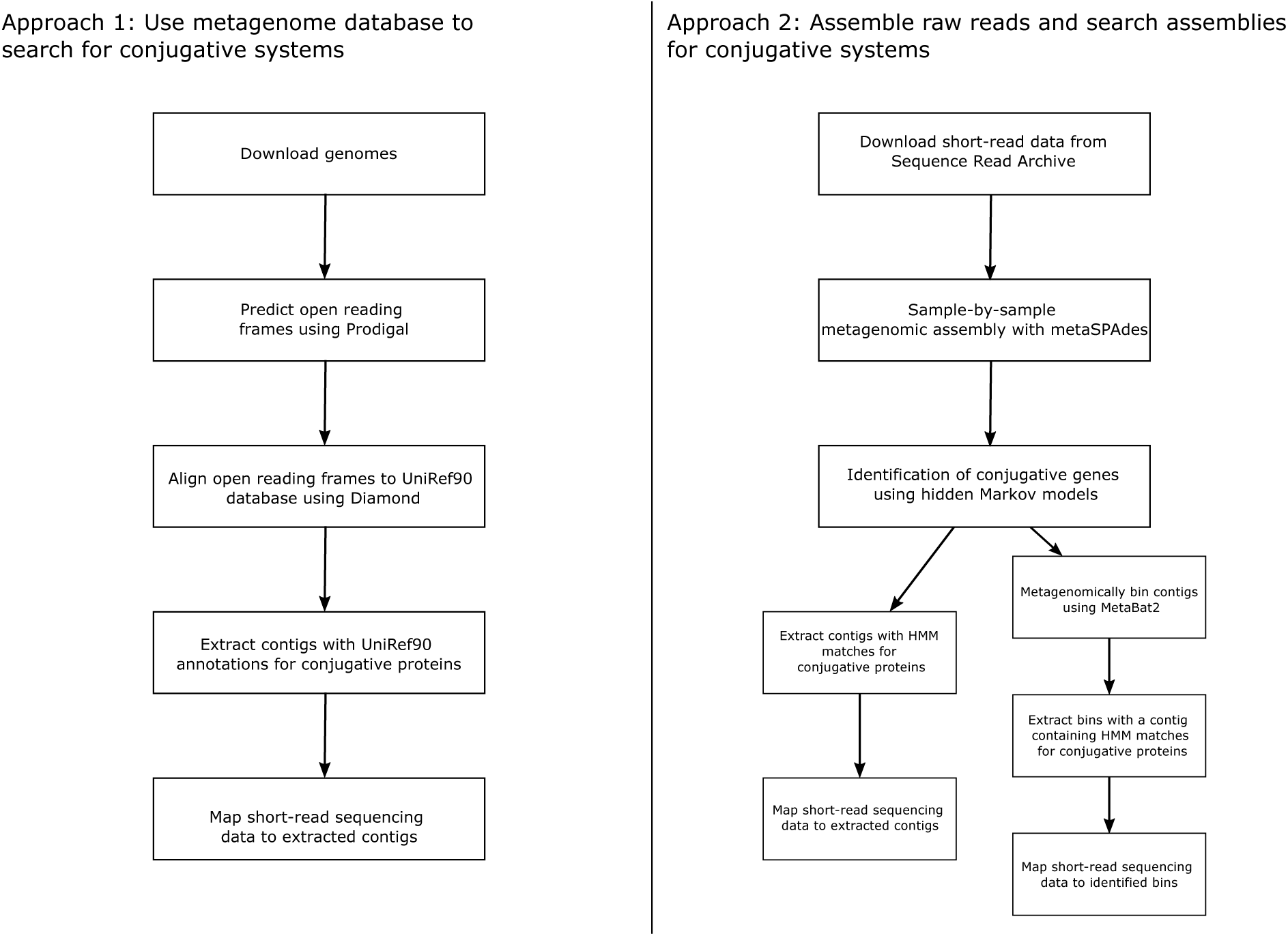
Overview of methods employed in this study. In the left panel is the workflow used to identify conjugative systems from previously assembled human gut bacterial genomes. The right panel outlines the workflow for the assembly of select North American samples and the use of pHMMs to identify the conjugative systems.

## Methods

### Assembly and identification of conjugative systems in North American short-read data

Samples belonging to a general North American (n=50) and a North American pre-term infant cohort (n=51) were assembled *de novo* (Supplemental Table 1). Reads from these samples were downloaded from the Sequence Read Archive using the SRA toolkit version 2.9.2, deduplicated with dedupe.sh [25], and trimmed with Trimmomatic version 0.36 [26] with options LEADING:10 TRAILING:10. Processed reads were assembled sample-by-sample using SPAdes version 3.14.0, option --meta [27]. The resultant assemblies were imported into Anvi’o version 6.0 [28] where the presence of T4SS, T4CP, and relaxase proteins were predicted using the anvi-run-hmms module, which integrates HMMER3 functionality [29]. Contigs that contained pHMM matches for all three classes of conjugative proteins were extracted and annotated by aligning open reading frames (ORFs) predicted with Prodigal version 2.6.3 [30] to the UniRef90 database [24]. Subregions of the contigs where annotations for conjugative proteins were present, with no more than 20 ORFs between successive UniRef90 annotations for conjugative proteins, were extracted. The processed read data were mapped to the extracted conjugative systems using Bowtie2 version 2.3.5 [31] with the settings --no-unal --no-mixed --no-discordant. Extraction of the subregions was to avoid an artificially high proportion of reads mapping in samples where the bacterium is present, but the ICE has not integrated in its chromosome (Supplemental Figure 1). Taxonomic prediction of the contigs was conducted with Kaiju version 1.7.2 utilizing the RefSeq non-redundant protein database [32]. MOB-suite verion 1.4.9.1 was utilized to characterize the incompatibility grouping of the conjugative system, if possible [33]. PlasFlow version 1.1.0 was used to classify whether the system was chromosomally integrated or a plasmid [22]. The proportion of reads mapping to the conjugative systems was extracted from the Bowtie2 output, and the mapping data was visualized using Anvi’o [28]. Raw counts of reads mapping to the extracted conjugative systems were transformed using a centered log-ratio. The principal component coordinates of the first 2 components were used for clustering by hdbscan [34].

### Reference human gut metagenome set

A near-complete and non-redundant set of human gut microbiome genomes were downloaded from the European Bioinformatics Institute FTP site (ftp://ftp.ebi.ac.uk/pub/databases/metagenomics/umgs_analyses/) [21]. These genomes were assembled from 13,133 metagenomic samples using SPAdes [27] and binned using MetaBAT2 [35]. The quality of binned genomes were assessed using CheckM [36]. High-quality genomes were defined as >90% completeness and <5% contamination and medium-quality genomes were defined as >50% completeness and <10% contamination, and these genomes were used to create the non-redundant set of genomes. The program dRep was used to cluster the genomes at 99% sequence identity [37] thereby dereplicating the genome bins, creating a set of 2505 genomes.

### Identifying and quantifying conjugative systems in reference human gut metagenome set

ORFs were predicted in the genome by Prodigal version 2.6.3 [30]. The predicted protein sequences were then aligned to the UniRef90 database [24] using the Diamond protein aligner version 0.9.14 [38]. Contigs were extracted from the genomes if they contained annotations for a relaxase/mobilization protein and a type IV secretion/type IV coupling protein using a word-search strategy. MOB-suite verion 1.4.9.1 was utilized to characterize the incompatibility grouping of the conjugative system, if possible [33]. PlasFlow version 1.1.0 was used to classify whether the system was predicted to be chromosomally integrated or located on a plasmid [22]. Short-read data from 785 samples (Supplemental Table 2) were downloaded from the Sequence Read Archive using the SRA toolkit version 2.9.2. The reads were processed and aligned to the regions of the contigs where type IV conjugative systems were located using the previously described methods. The principal component coordinates of the first 3 components were used for clustering by hdbscan [34].

### Binning of Assemblies

For each assembly, all 101 samples were mapped to the contigs using Bowtie2 [31]. The mapping files were sorted and indexed with SAMtools [39] and then the assemblies were binned using MetaBAT2 version 2.12.1 [35]. CheckM version 1.1.2 was used to assess the quality of the resultant bins [36]. High-quality bins were defined using the same cutoffs (>90% completion and <5% redundancy) as Almeida *et al* (2019) defined. Bins not passing that threshold were classified as “low-quality”. The previously identified contigs with conjugative systems were classified based on their presence in bins, and the types of bins they were present in. Results of this classification were visualized using SankeyMATIC (http://sankeymatic.com/).

## Results

### Conjugative systems identified from assembly of short-read data distinguish North American cohorts

51 samples from a pre-term infant cohort and 50 from a general North American cohort were assembled sample-by-sample using metaSPAdes [27] to identify T4SS conjugative systems from a full pool of assembled contigs (i.e. not binned) and to compare the relative abundances of these systems between cohorts. For these analyses, contigs with conjugative systems were defined by pHMM matches for a relaxase, a type IV coupling protein, and a type IV secretion system, which offers a fast and precise method to annotate a limited number of protein families. From the assembly of the pre-term infant cohort 96 of 470500 contigs met the criteria, whereas 268 of 15100646 contigs from the general cohort did (Supplemental Table 3). Predicted ORFs from contigs with conjugative systems were aligned to the UniRef90 database and the subregions with conjugative systems were extracted. The short-read data from all 101 samples were mapped to all 391 subregions of conjugative proteins.

The patterns of conjugative system occurrence and relative abundance in the two cohorts are distinct (Figure 3). Conjugative systems belonging to the *Proteobacteria* phylum were only assembled from pre-term infant samples and did not have any apparent occurrence in the general population. Furthermore, most identified conjugative contigs were private to the cohort they were assembled from. When the log-transformed principal component analysis data were clustered with hdbscan [34], the two cohorts formed separate clusters (Figure 4, Supplemental Figure 2). These findings show that North American pre-term infants have a strikingly different array of conjugative systems than the general North American cohort.

**Figure 3:**
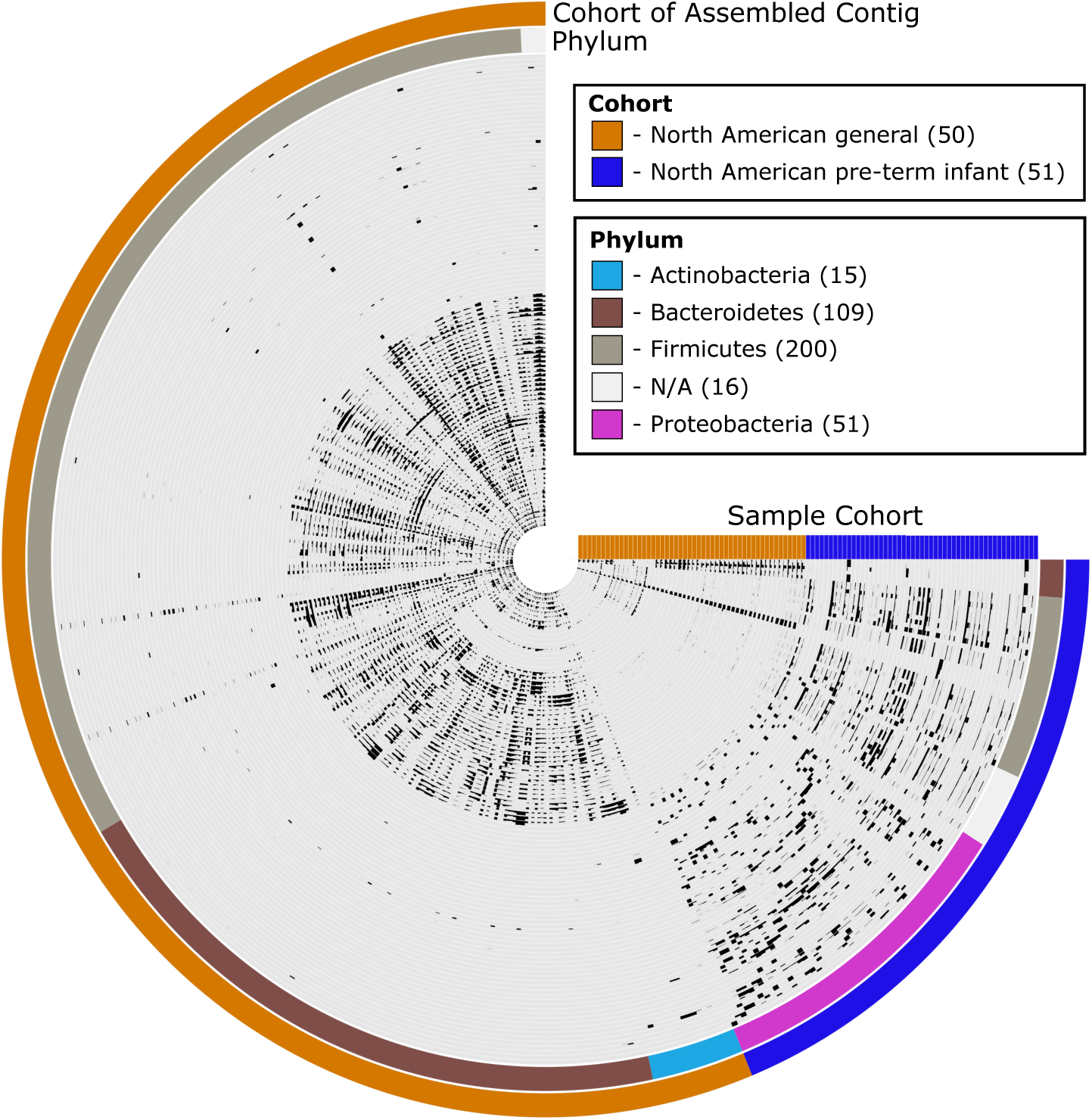
Anvi’o cladogram of potentially conjugative contigs from 51 North American pre-term infants samples and 50 general North American samples. Inner rings of the phylogram represent individual samples, second-most outer ring being the phylum of conjugative system as predicted by Kaiju, and the outermost ring represents the cohort that the conjugative contig was assembled from. Each slice of the circle phylogram are individual conjugative contig identified by pHMMs of conjugative proteins. For the inner plot, intensity of the position represents the mean coverage of the contig for a given sample proportional to the other conjugative systems.

**Figure 4:**
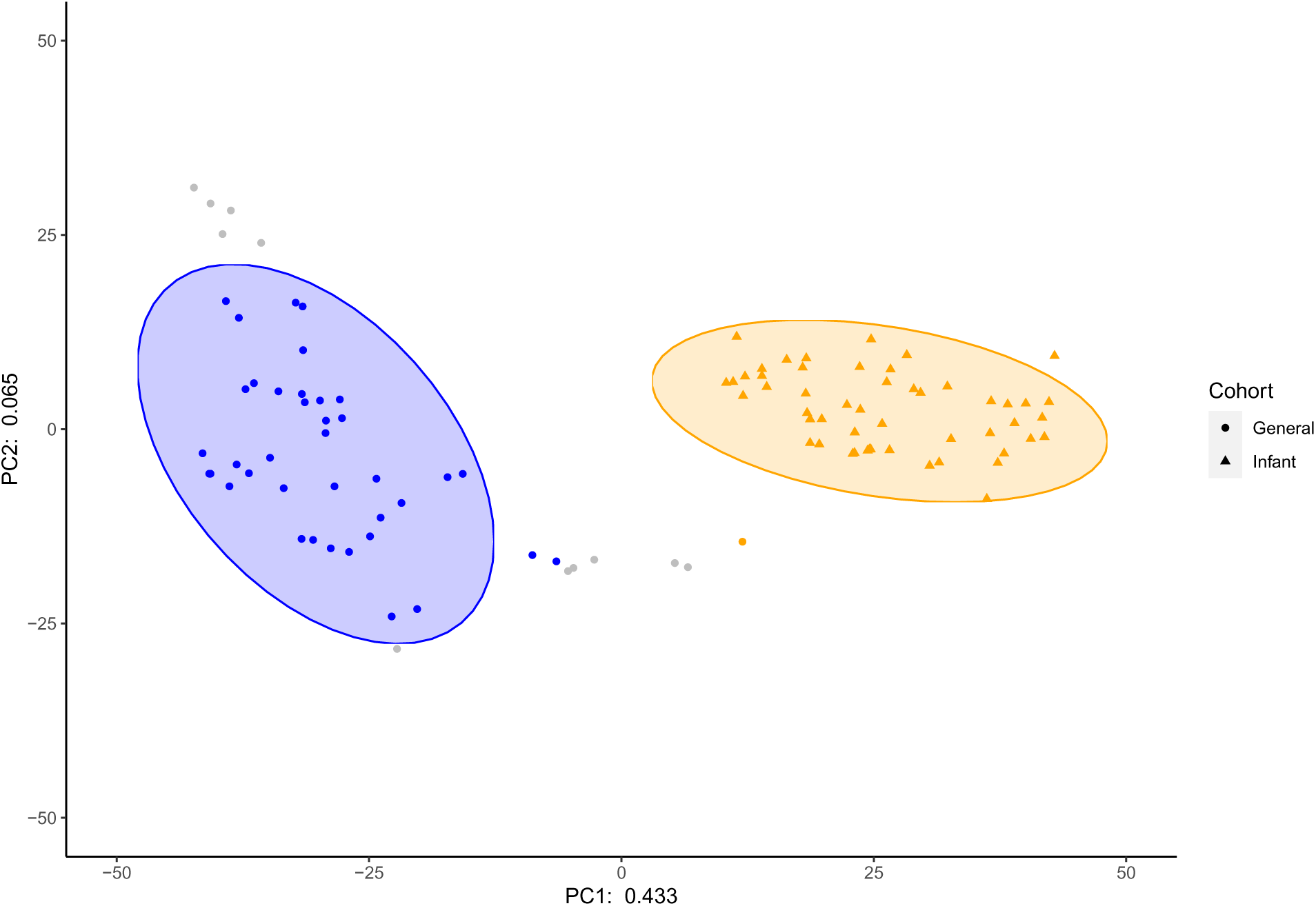
Clustering of the principal component coordinates of the CLR transformed relative abundances of the extracted conjugative regions from the assemblies of North American datasets. Ellipses represent a 95 percent confidence interval using a multivariate t-distribution.

### Mapping human gut microbiome data from cohorts to conjugative systems reveals distinct patterns

Next we explored relative abundances of conjugative systems in a greater number of cohorts, without having to conduct computationally-expensive metagenomic assemblies. For this analysis, conjugative systems were identified from a set of 2505 bacterial genomes, which represent a non-redundant and near-complete picture of the human gut microbiome [21]. A total of 1598 contigs from 787 genomes that contain UniRef90 annotations for relaxase/mobilization and T4SS/T4CP proteins were identified (Supplemental Table 3). From these contigs, 3216 subregions where conjugative protein annotations were concentrated on the contig were extracted (Supplemental Figure 1), with 2413 being >1kb in size and used for visualization. Short-read human gut microbiome sequencing data from 785 samples, spread across 8 cohorts were aligned to the extracted subregions (Supplemental Table 1). With the conjugative systems identified from the human gut metagenome set, the two North American cohorts that were previously analyzed are still distinct, albeit in a different way (Figure 5). Only a very small number of the reads from the North American and European Infant cohorts mapped to conjugative systems. The only notable signal is in the *Proteobacteria* phylum for the North American pre-term infants, a finding consistent with what was found by *de novo* assembly. The West African and South American cohorts also share similar characteristics as both have an overall lower apparent relative abundance of conjugative systems compared to the other non-infant cohorts, particularly in the *Bacteroidetes* phylum. The other four cohorts appear similar with regards to the presence and absence of the conjugative systems. The cohorts separated into three distinct clusters (Figure 6, Supplemental Figure 3), when the principal components of the centered log-ratio transformed data were clustered using hdbscan [34]. In this analysis infant cohorts were excluded because of their extreme sparsity. The majority of the West African and South American samples clustered together consistent with Figure 5. Not readily apparent from the cladogram was the East Asian cohort that clustered primarily on its own. The North American Indigenous, North American general, and Western European general samples largely clustered together. Like the conjugative systems identified from the *de novo* assemblies of short-read data, the relative abundances of conjugative systems identified from a human gut metagenome set separated cohorts into distinct groups.

**Figure 5:**
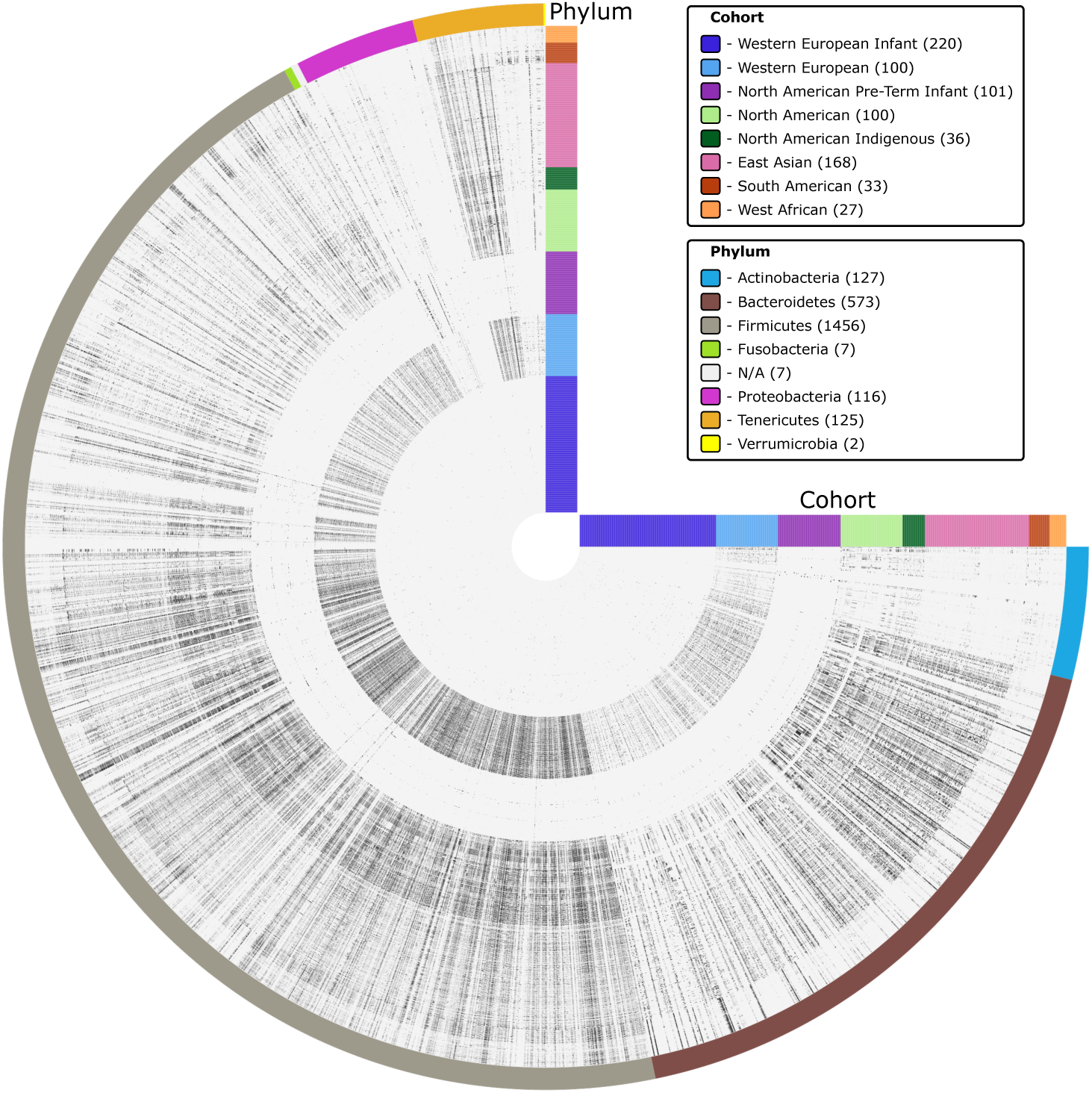
Anvi’o cladogram of potentially conjugative systems originating from 785 samples across 8 cohorts. Inner rings of the phylogram represent individual samples and the outermost ring being the phylum of conjugative system. Each slice of the circle phylogram are individual conjugative regions. For each point on the inner plot, the intensity of the black colouring represents the mean coverage of the system for a given sample proportional to the other conjugative systems.

**Figure 6:**
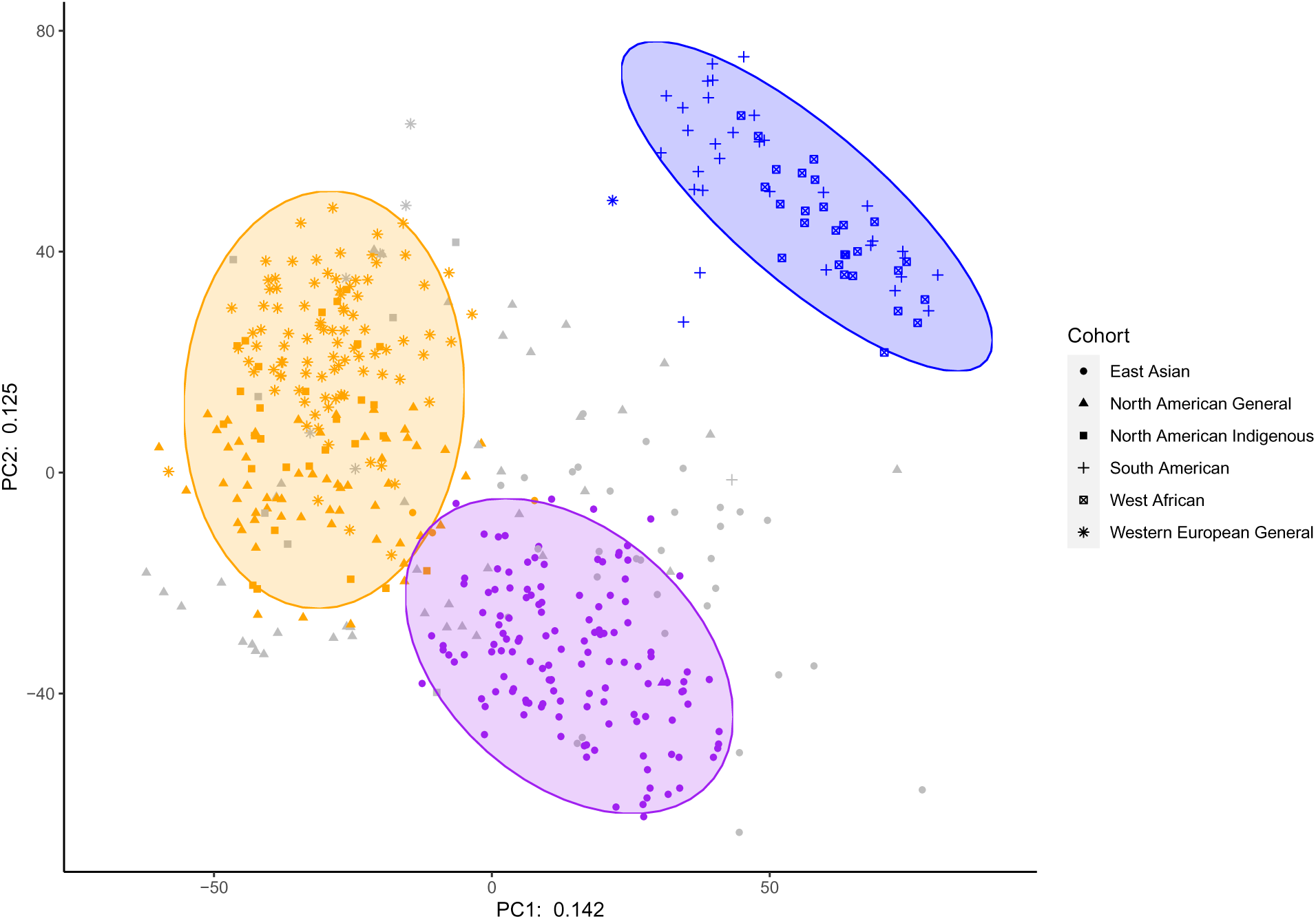
Clustering of the principal component coordinates of the CLR transformed relative abundances of the extracted conjugative regions from the genome database. Ellipses represent a 95 percent confidence interval using a multivariate t-distribution.

### Percentage of reads mapping to conjugative systems is inconsistent between methods

For the cohorts that were examined using both methods of conjugative system identification, the percentages of total reads mapping to the identified conjugative systems were not equal between methods (Figure 7). From the conjugative systems identified from the general North American cohort assembly of the short reads the mean percentage of reads mapping was 0.62% (95% CI [0.22%,1.1%]), whereas the mean percentage of reads mapping to the conjugative systems identified from the genome set was (2.69% 95% CI [1.25%,4.64%]). A lower average percentage in the assembled dataset than the genome set conjugative systems suggests that the assemblies of the short read sequences were not able to successfully capture the full diversity of conjugative systems found in an average North American individual. Conversely, for the pre-term North American infants, the mean percentage of total reads mapping to conjugative systems identified from the assemblies was 3.33% (95% CI [0.00182%,2.39%]), whereas from the reference genome set the mean percentage was (0.271% 95% CI [0.025%,1.21%]). In terms of the composition of the reference gut metagenome set, it is probable that the bulk of the bacterial genomes are sourced from deeply sequenced cohorts, like a general North American population, rather than a niche cohort like pre-term infants. This results in a more precise relative abundance estimate of conjugative systems in the general North American cohort than in the pre-term infant cohort. These findings together suggest that the total percentage of reads mapping to conjugative systems is not particularly informative and these data should be treated as compositions with relative abundances of features being compared between groups.

**Figure 7:**
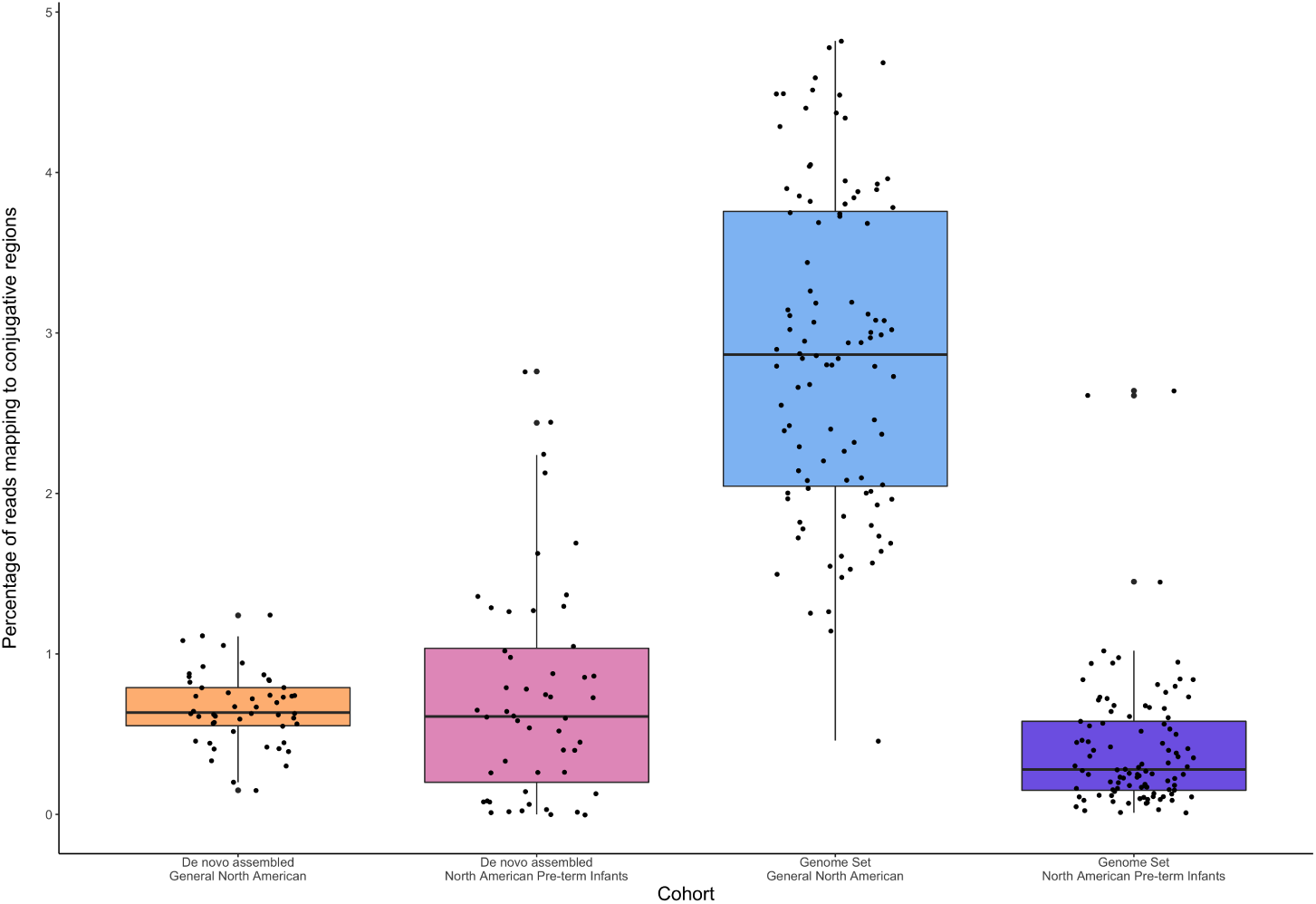
Plot of the percentage of total reads from North American datasets mapping to conjugative regions extracted from de novo assemblies and the genome set, separated by cohort.

### The majority of conjugative systems identified by assembly are omitted from metagenomic bins

The assemblies were binned using MetaBAT2 [35], which was also used to bin the MAGs in the human gut genome set used in the prior analyses[21] to further explore how conjugative systems are distributed within common metagenomic analyses. Of the 364 assembled contigs containing pHMM matches to all three protein categories, 270 were not included in any metagenomic bins (Figure 8). For the 94 contigs included in metagenomic bins, 65 of those were found in high-quality bins (>90% completion and <5% redundancy). Among the 29 contigs included in bins that do not meet the aforementioned threshold, 8 are within bins that are greater than or equal to 1 megabase in size, potentially suggesting that fragments of a conjugative plasmid may have binned together.

**Figure 8:**
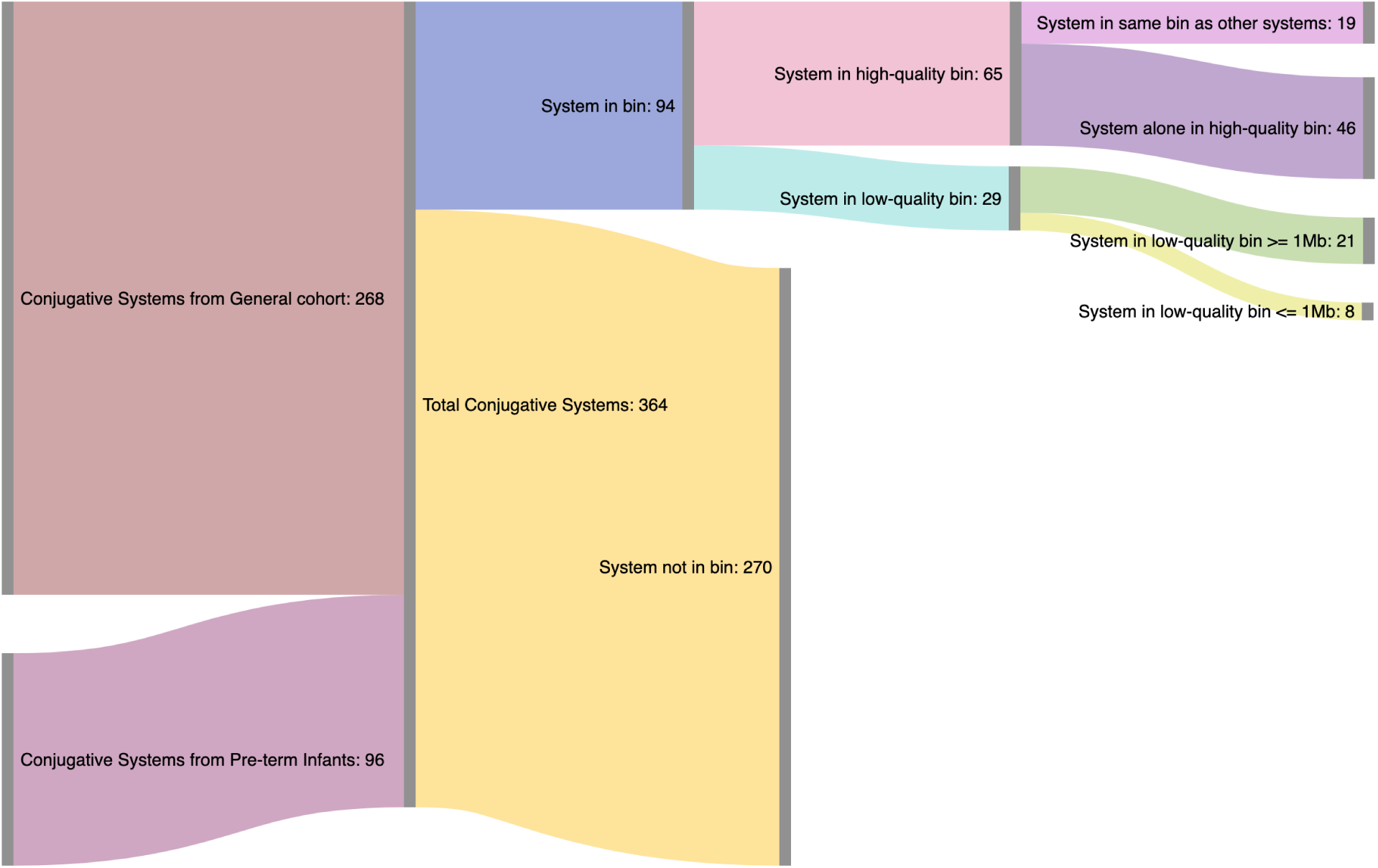
Sankey diagram representing the flow of 364 contigs containing conjugative systems into bins generated by MetaBAT2 from assembled data.

## Discussion

The relative abundances of conjugative systems identified from MAGs and isolate genomes from the human gut differ between cohorts similarly to how relative abundances of human gut MAGs of different species are differential between cohorts [40]. The infant cohorts stood out the most from the other cohorts; the infant gut microbiome is composed largely of members of the genus *Bifidobacteria* and is recognized as being distinct to the microbiomes of adults over the first few years of life [41]. Furthermore, we observed that the gut microbiome of pre-term infants were distinct from other infants and adults. This could be because of exposure to antibiotics from birth and colonization by opportunistic *Proteobacteria* pathogens such as members of the genera *Escherichia*, *Klebsiella*, and *Enterobacter* [42]. As shown in Figure 2a, the only conjugative systems that showed a signal for the samples belonging to the North American pre-term infants were those belonging to the *Proteobacteria* phylum. The degree of difference in relative abundances of conjugative systems between infants and adults suggests that conjugative systems could be a potential biomarker for age, or for the relative maturity of the infant microbiome.

The relative abundances and distributions of conjugative systems in the West African and South American cohorts were distinct from the other non-infant cohorts, which is similar to the findings in the relative abundances of bacterial species in these cohorts [40]. As well, the East Asian cohort clustered separately from the other non-infant cohorts. These findings suggest that conjugative systems might be useful biomarkers for other factors beyond just age and further focus on geographical or health-related differences may yet reveal additional separation between cohorts based on the relative abundances of conjugative systems.

Comparing the findings between methods in Figure 7, it is clear that the total percentage of reads mapping to conjugative systems is not an effective metric; the differences between the relative abundances in conjugative systems that were present from the genome set between the general North American and North American pre-term infants did not persist when examining the relative abundances of the assembled systems. Comparing Figure 3 and Figure 5, the conjugative systems assembled in pre-term infants are almost entirely missing in the genome set. It is clear that there is a degree of incompleteness in terms of infant conjugative systems in a database consisting of primarily MAGs assembled from non-infant datasets. However, for the non-infant data, the genome set appeared to capture a larger percentage of the conjugative systems. Neither method of identifying and quantifying conjugative systems is perfect; the reference bacterial genome set may be incomplete for less commonly studied cohorts, but the sequencing depth of a single sample may not be sufficient to assemble the less abundant conjugative systems in an environment.

We found that a reference bacterial genome set can be useful for identifying coarse differences in the conjugative systems between populations; however, this method may not capture the true diversity of conjugative between populations, because many conjugative systems may be omitted. To produce MAGs, contigs generated by metagenomic assembly are typically binned using a program such as MetaBAT2 [35]. Conjugative systems are often more AT rich than the parent genomes [1], which would result in the conjugative system and cognate genome not occurring in the same metagenomic bin. Additionally, plasmids are not necessarily maintained in a unit copy number within the cell, causing differential sequence coverage in comparison to the parent genome, which results in plasmids being excluded from MAGs. Therefore to capture a more complete image of the conjugative systems present in an environment, identification of the systems must take place before binning.

The assembled contigs were binned with MetaBAT2 [35] as a way of quantifying the effect of binning, which revealed that the vast majority of the assembled conjugative systems were not included in metagenomic bins and therefore would not be included in a MAG database, which confirms recent findings [23]. Many of the binned conjugative systems were not within a bin that would pass the quality cutoff to be included in the genome set as well [21]. Interestingly, eight of the conjugative systems were binned into low-quality bins that were smaller than <1MB in size, which may suggest that the fragments of a conjugative plasmid could be binned together, which would increase the completeness of the conjugative system.

## Conclusions

Conjugative systems differ between cohorts and require special consideration to ensure that they are included in analyses. ICEs and plasmids can carry harmful systems, such as antimicrobial resistance, but also can act as vectors for bile salt metabolism and for detoxification modules [3]. These cargo genes are relevant for research relating to the gut microbiome’s role in pathogenicity as well as metabolism and digestion. Comprehensive identification and quantification of conjugative systems could allow for association of conjugative systems with different health outcomes. Because assembled plasmid-based conjugative systems are rarely included in metagenomic bins [23], they need to be identified and analyzed outside of standard binning pipelines. At present, it is not possible to assemble complete plasmids from short-read metagenomic data [20], so it may helpful to identify bins containing conjugative systems in an attempt to cluster the fragments of plasmids present in an assembly together. Identifying type IV conjugative systems using pHMMs or annotations and using tools such as PlasFlow [22] to identify plasmids out of a full assembly in parallel with standard binning analyses will enhance research of the associations between the human gut microbiome and human health.

In the future, improvements in assembly and binning algorithms will continue to improve the recovery of low relative abundance conjugative elements and improve the completeness and accuracy of the assembled fragments. Additionally, long-read assembly permits the circularization of genomes and plasmids [43, 44] and the binning of plasmids to their cognate genomes using methylation data [45], which will reduce the ambiguity of the origins of conjugative systems (i.e. whether they are an ICE or independently circularized plasmid) and provide a more complete picture of the cargo they carry and the differences between cohorts.

---

**Supplemental Table 1:** Accession numbers and cohorts for samples used in *de novo* assembly workflow.

**Supplemental Table 2:** Accession numbers and cohorts for samples used in reference human gut metagenome set workflows.

**Supplemental Table 3:** Summary data for all predicted conjugative systems.

## Declarations

### Ethics approval and consent to participate

Not applicable.

### Consent for publication

Not applicable.

### Availability of data and materials

All code needed to reproduce the results are available on Github, https://github.com/bjoris33/humanGutConj_mSystems

### Authors’ Contributions

BRJ designed the experiments, analyzed and interpreted the data, and wrote the manuscript. TSB analyzed the data. TAH interpreted the data. DRE designed the experiments, interpreted the data, and provided funding. GBG designed the experiments, interpreted the data, edited the manuscript, and provided funding.

## Acknowledgements

We thank Daniel Giguere for his input on the analyses and figures.

---

**Figure 1:**
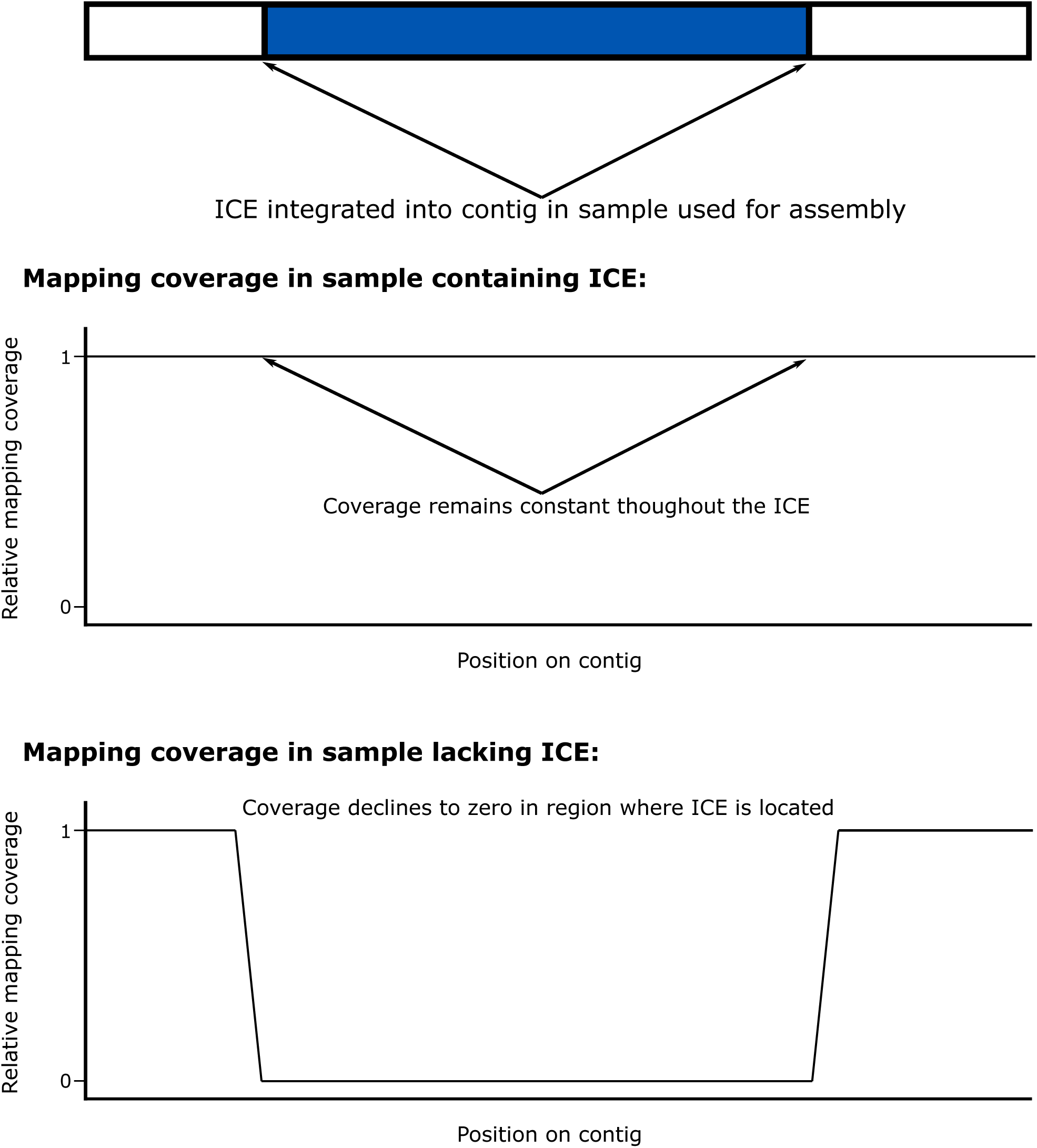
Conceptual diagram of the mapping coverage of an assembled integrative and conjugative element. The mapping coverage in the first plot shows an even mapping coverage across the contig because the ICE is present in the sample and the average mapping coverage of the contig would be an accurate metric. In the second plot, the ICE is missing in the sample and the mapping coverage falls to zero where the ICE is located on the contig. As a way to quantify the presence of the ICE, the average mapping coverage for the entire contig would be artificially high. Limiting the mapping to only the region containing the conjugative proteins solves this issue.

**Figure 2:**
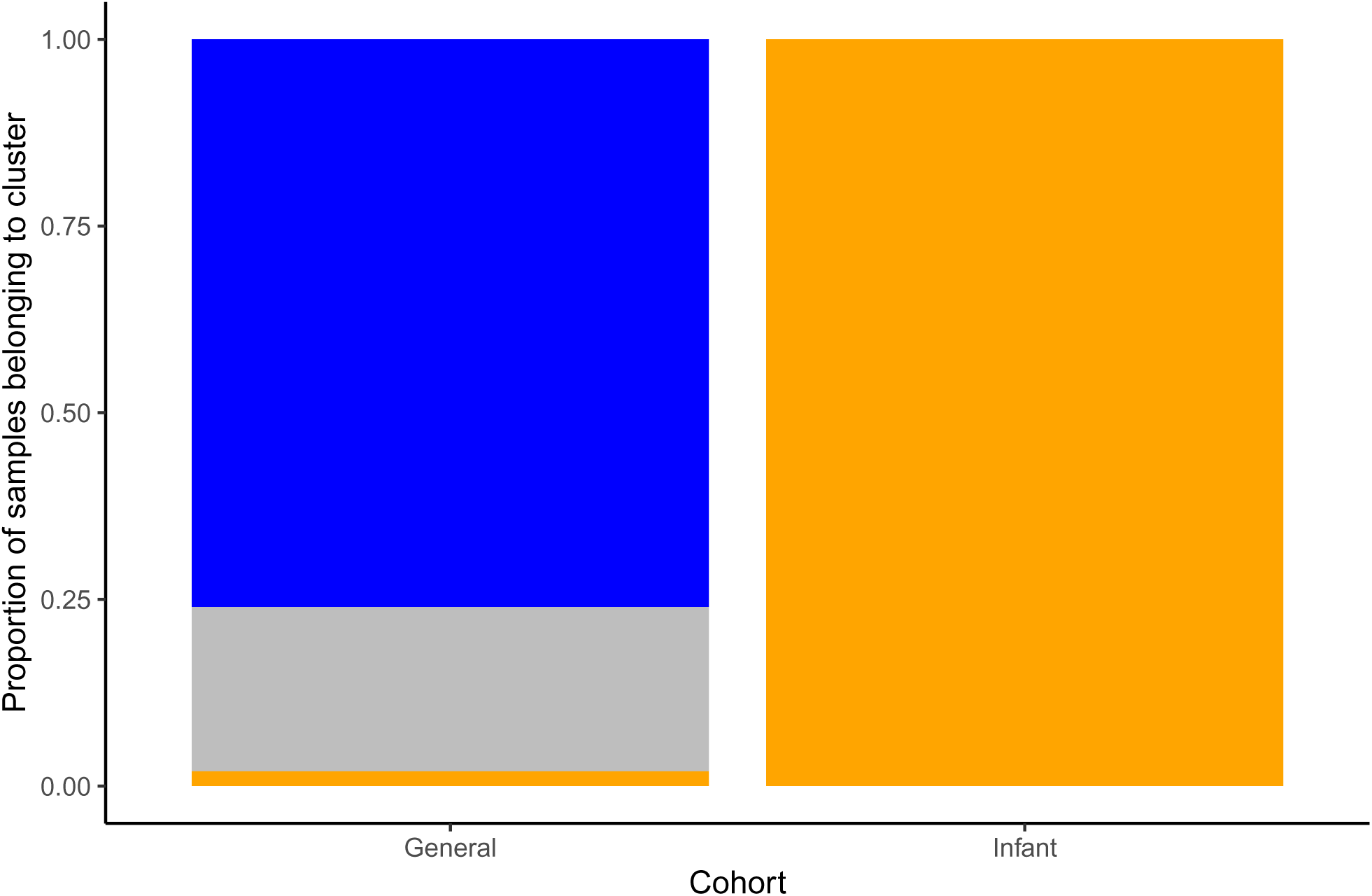
Stacked bar plot of the proportions of samples belonging to the hdbscan clusters from each cohort.

**Figure 3:**
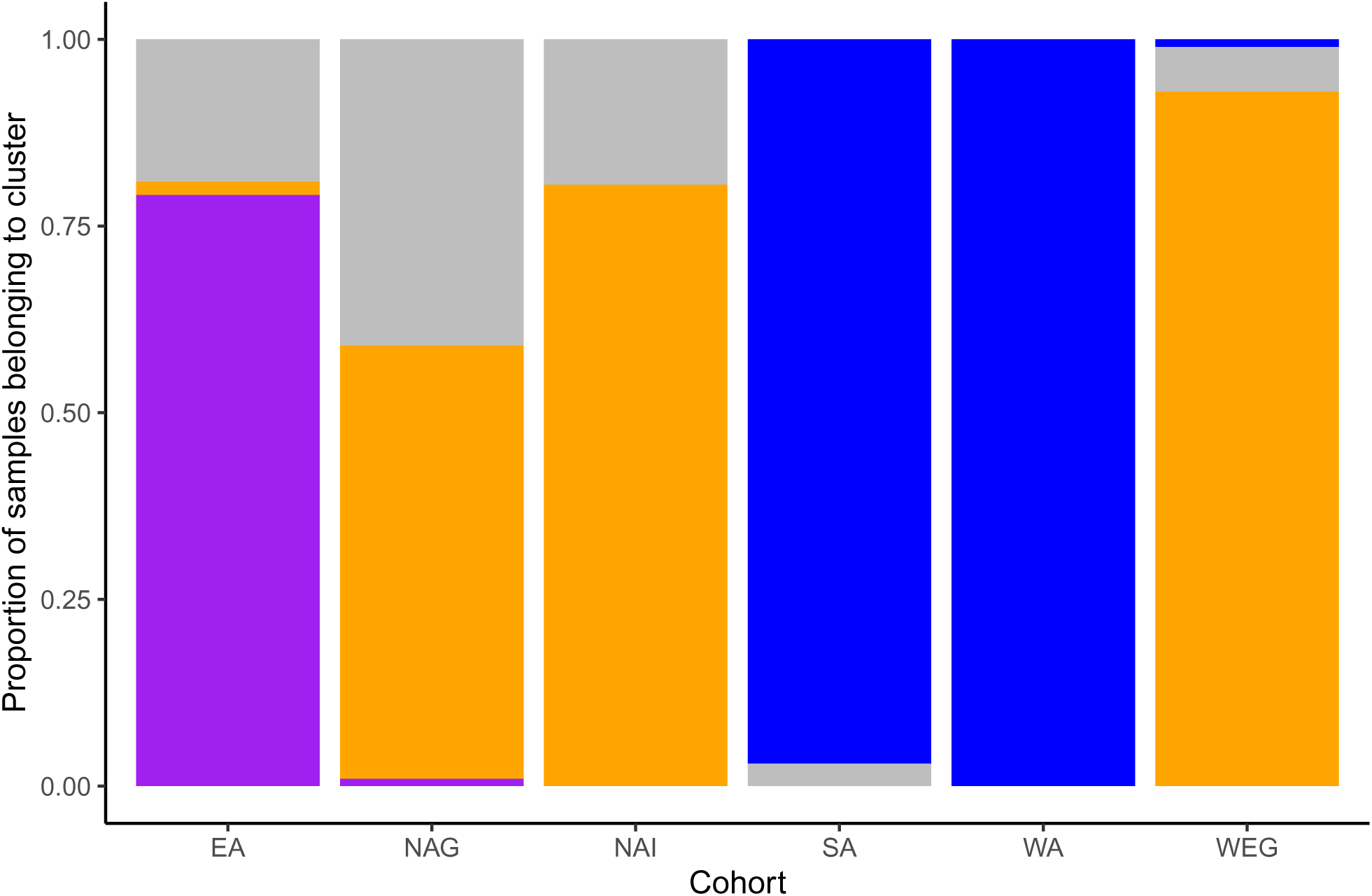
Stacked bar plot of the proportions of samples belonging to the hdbscan clusters from each cohort.

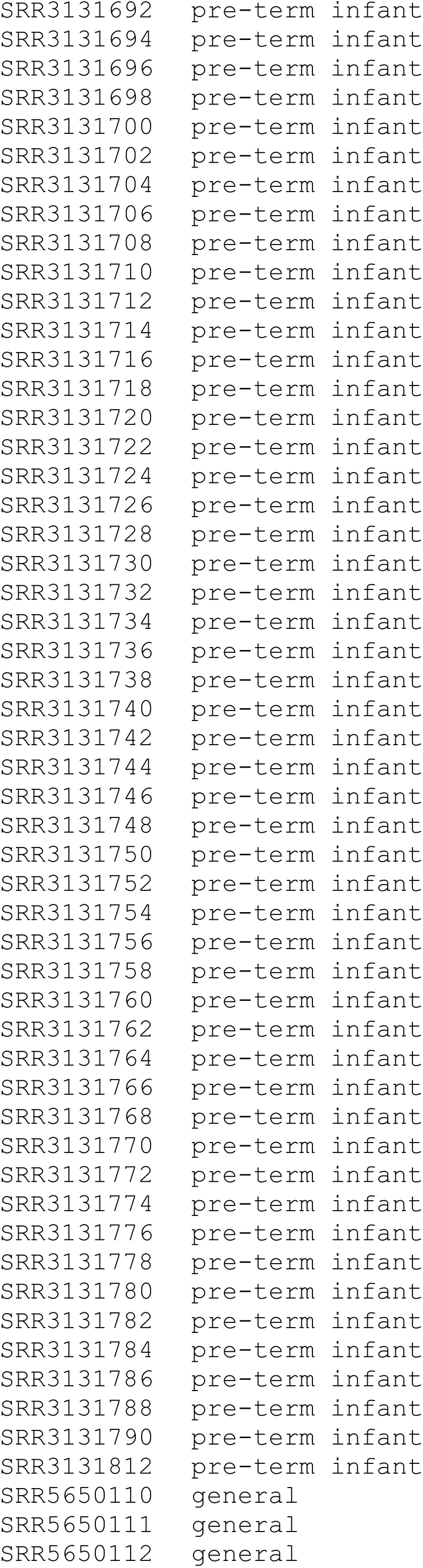
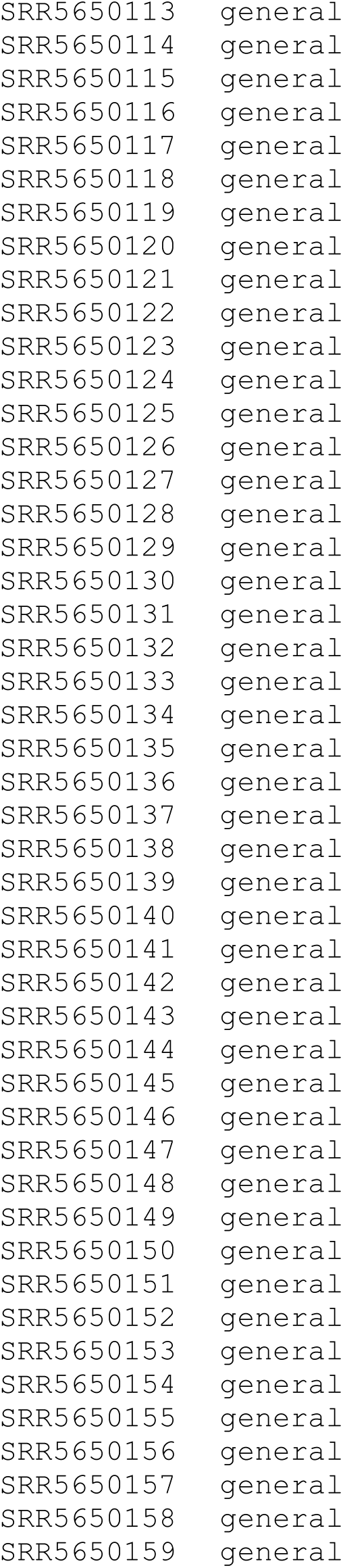
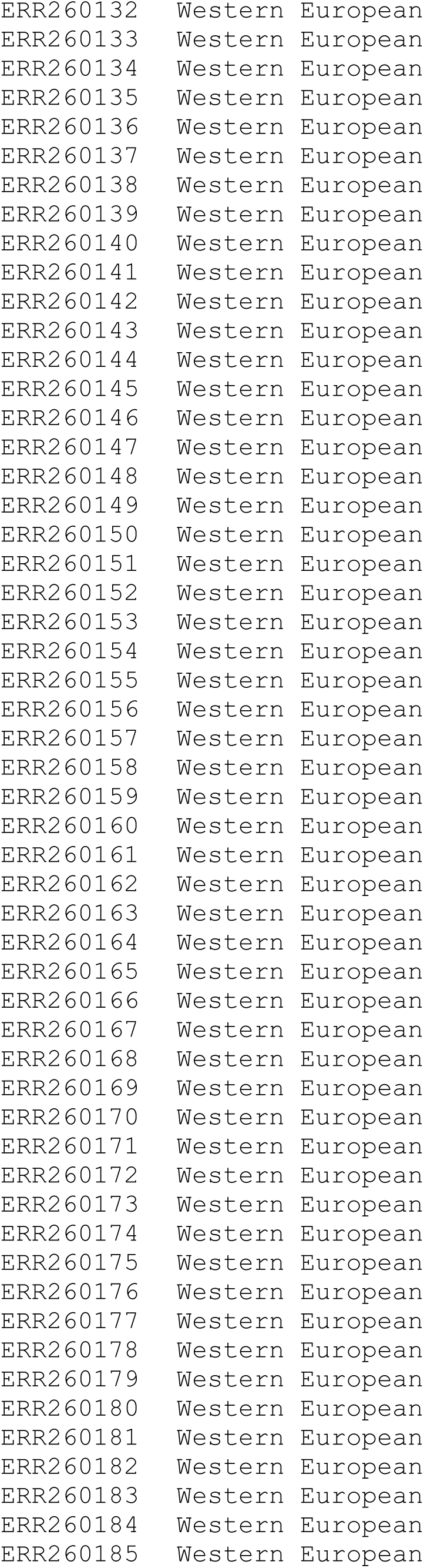
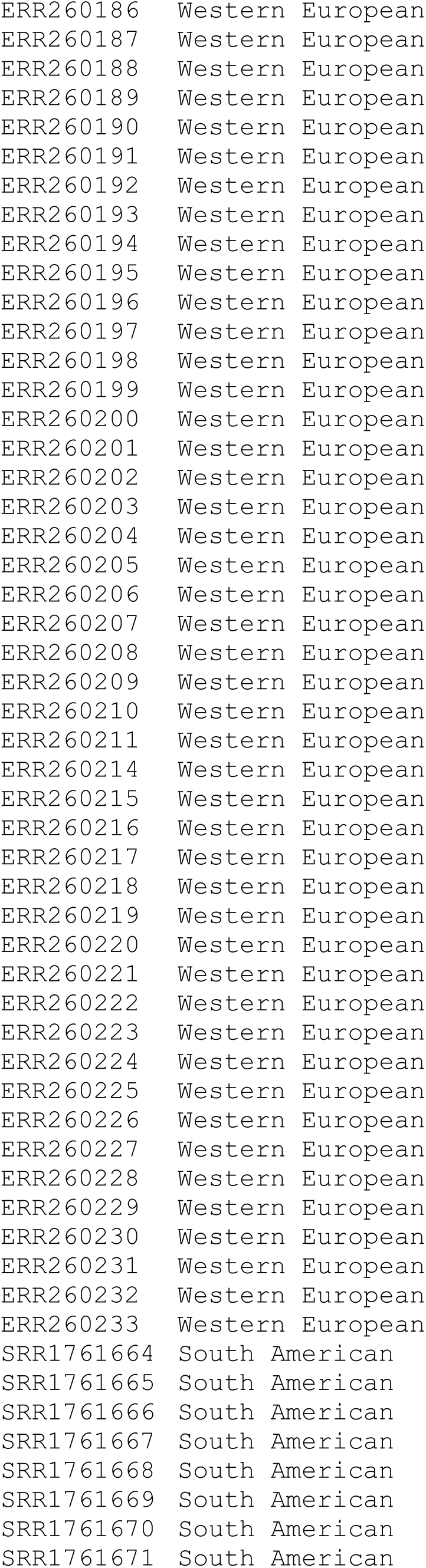
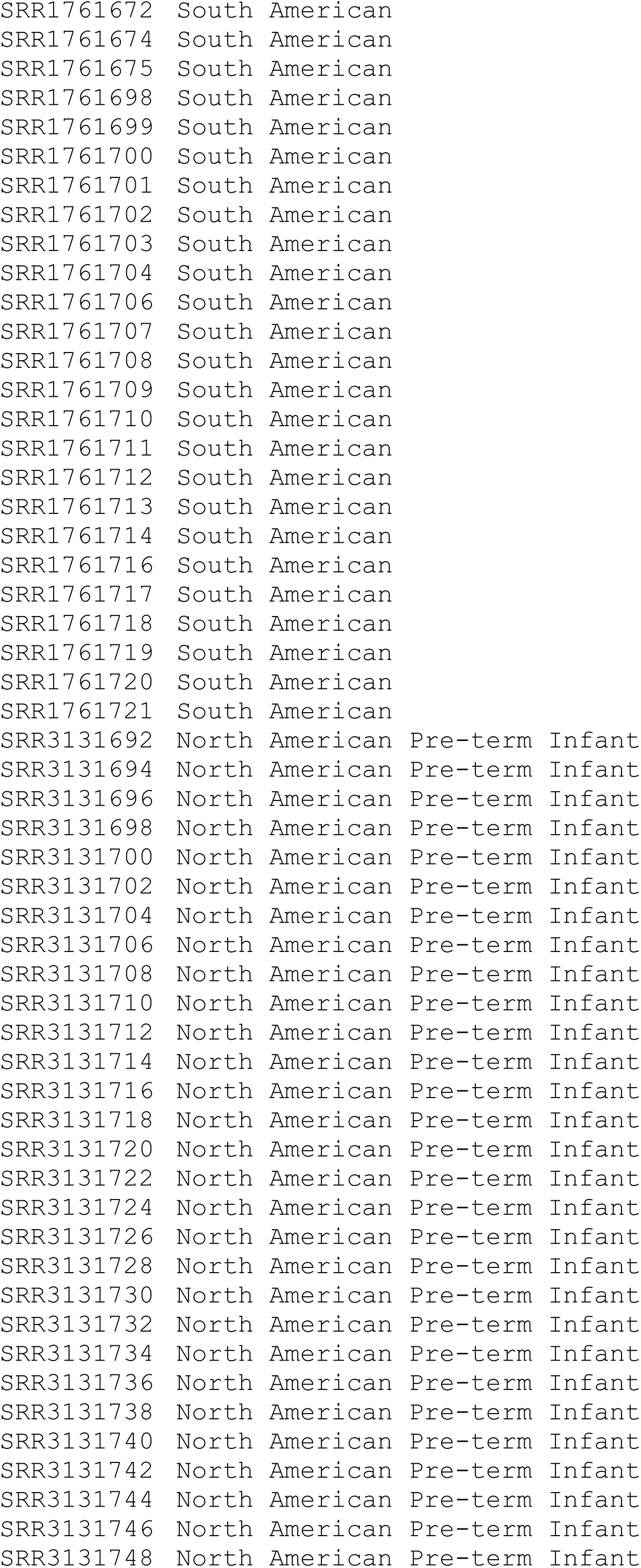
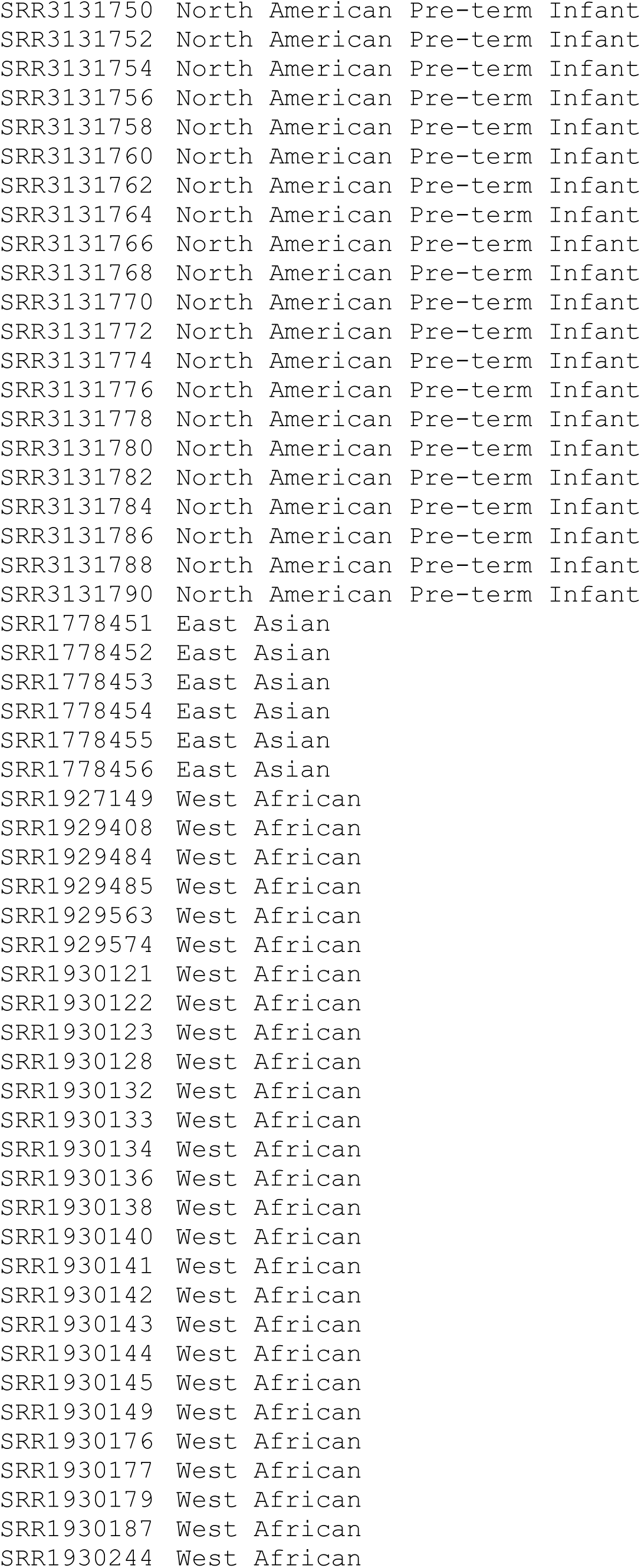
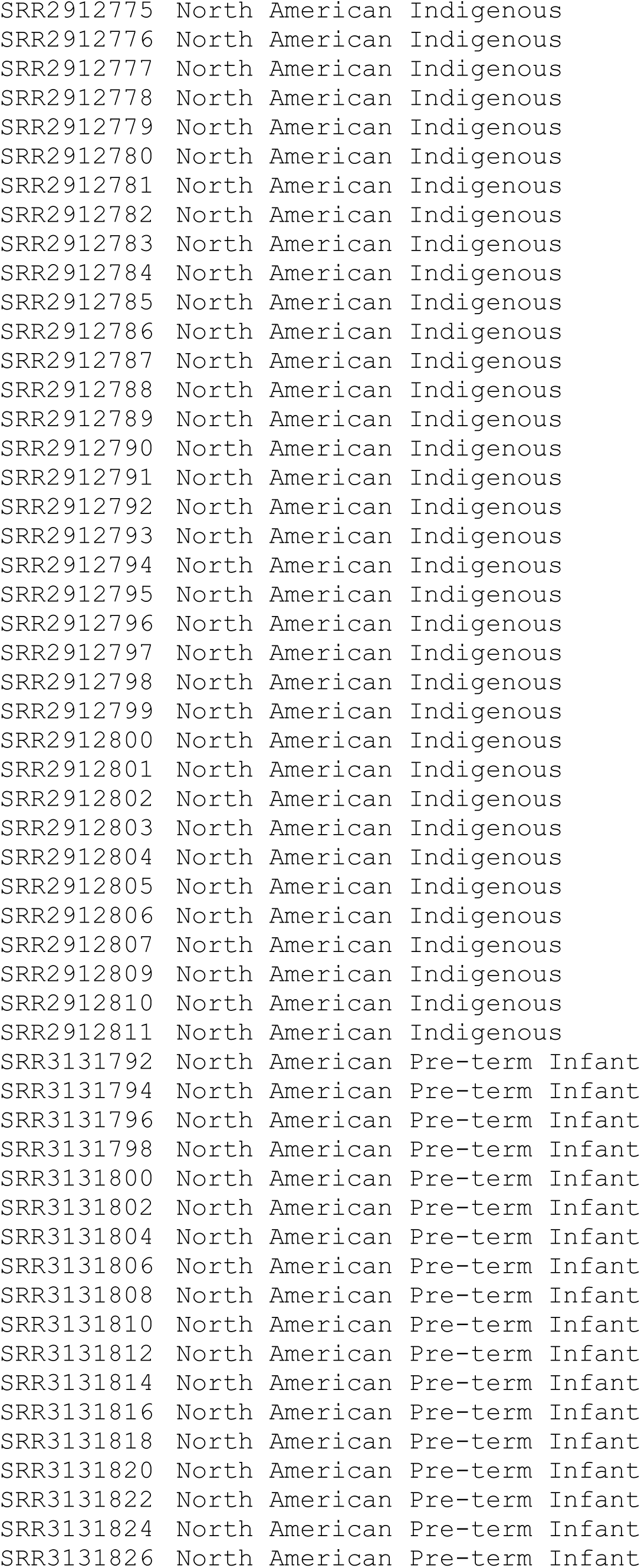
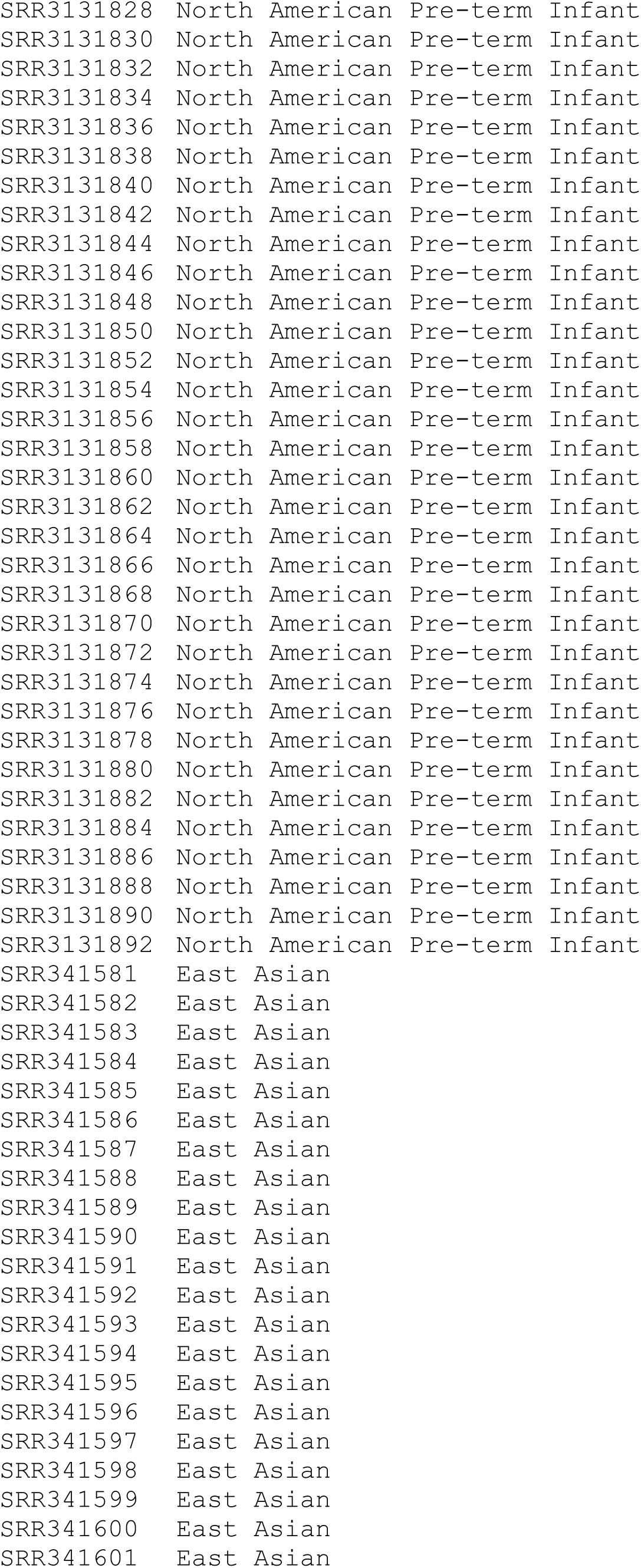
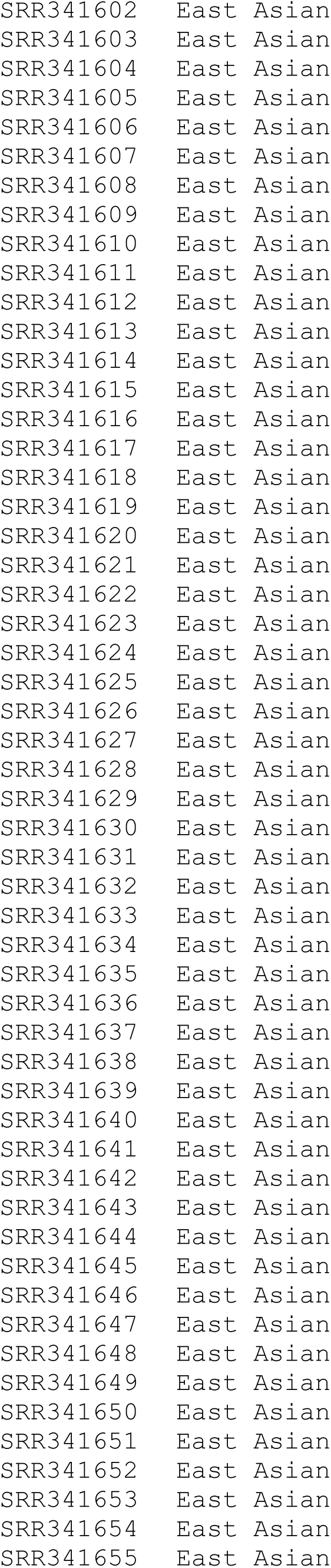
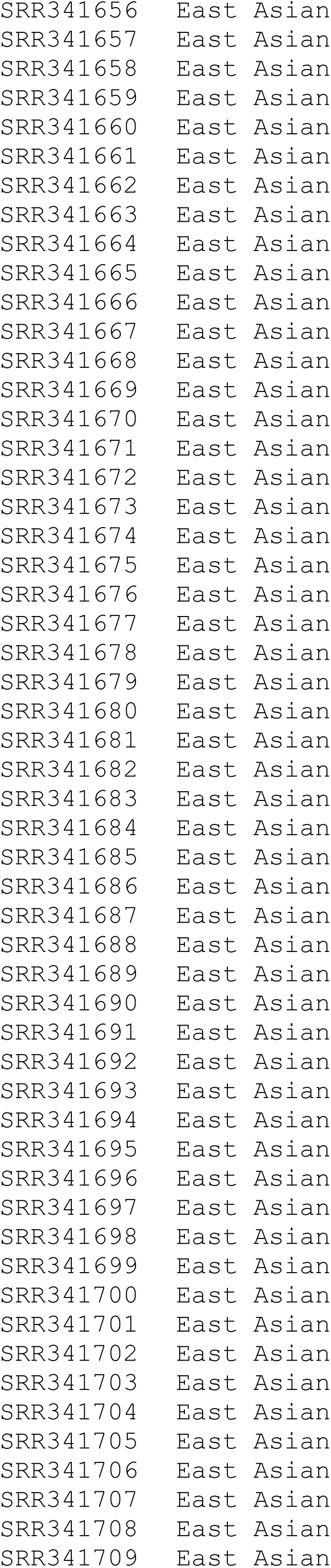
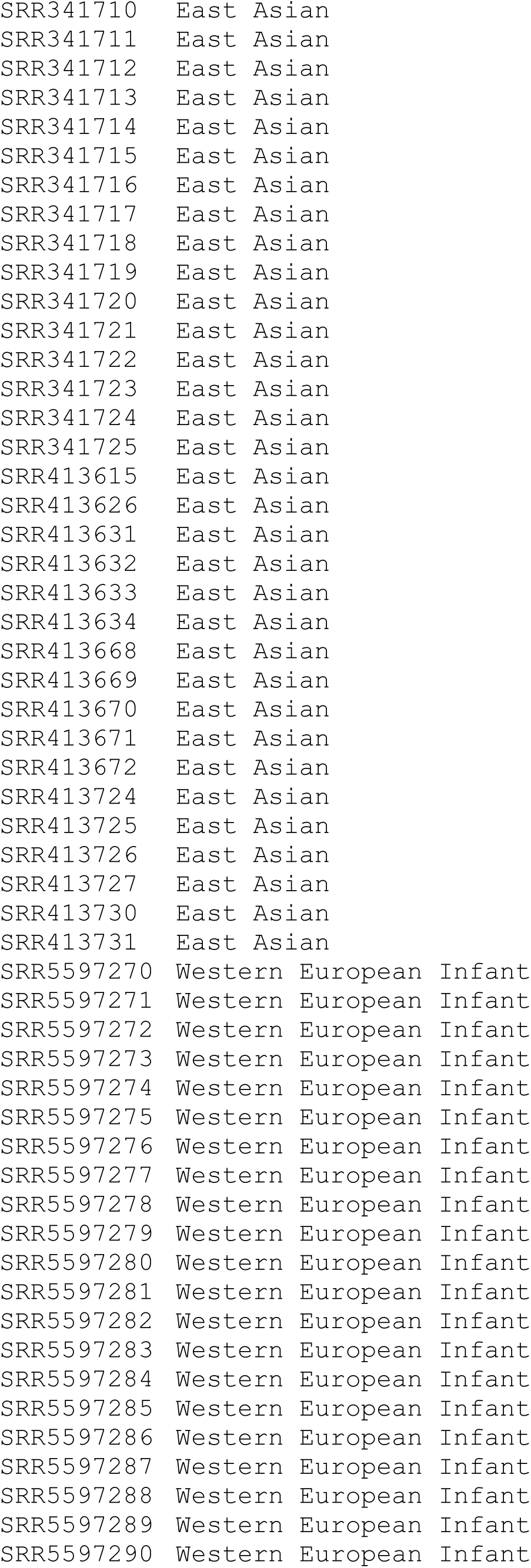
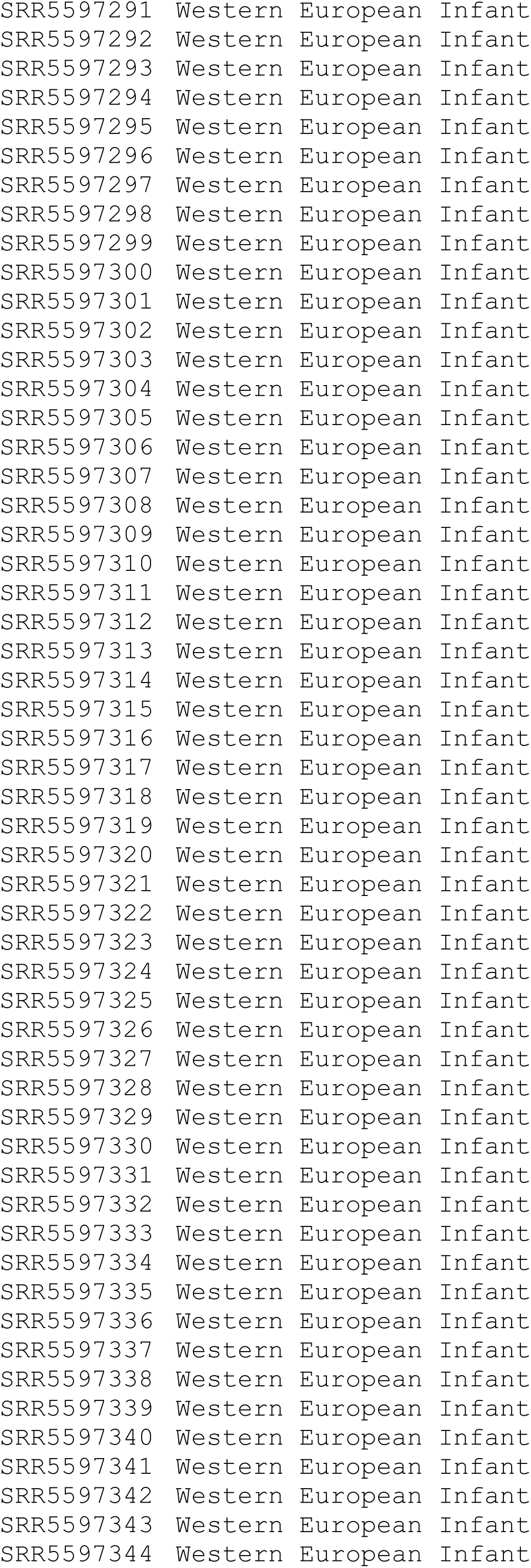
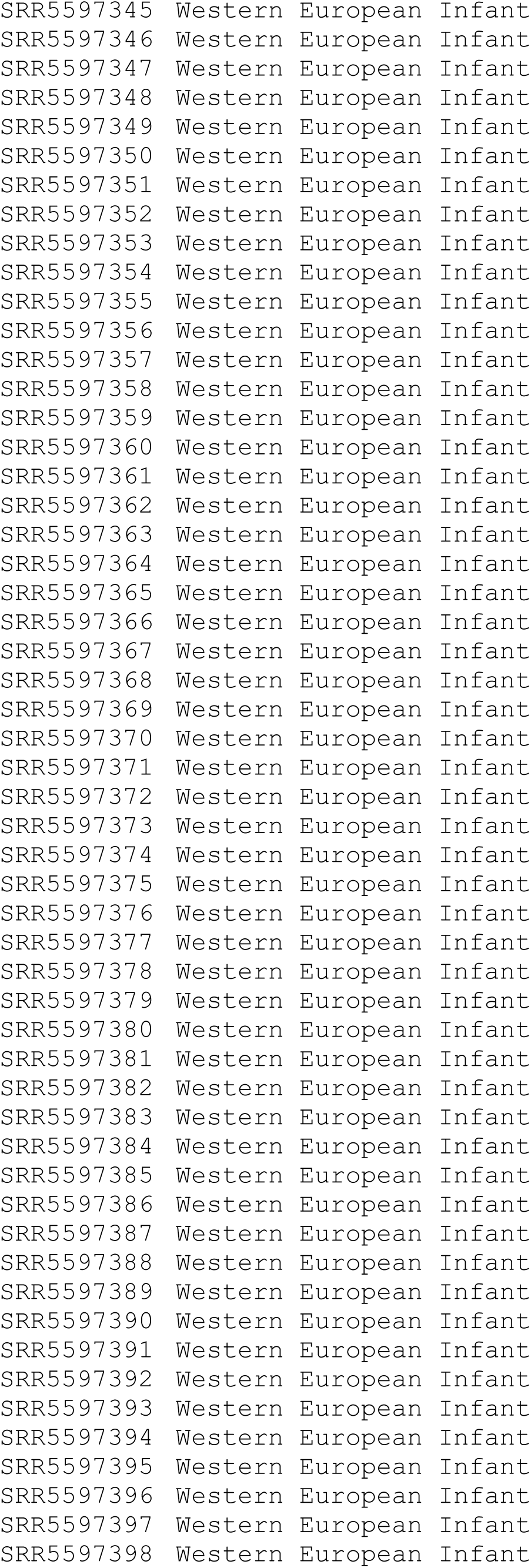
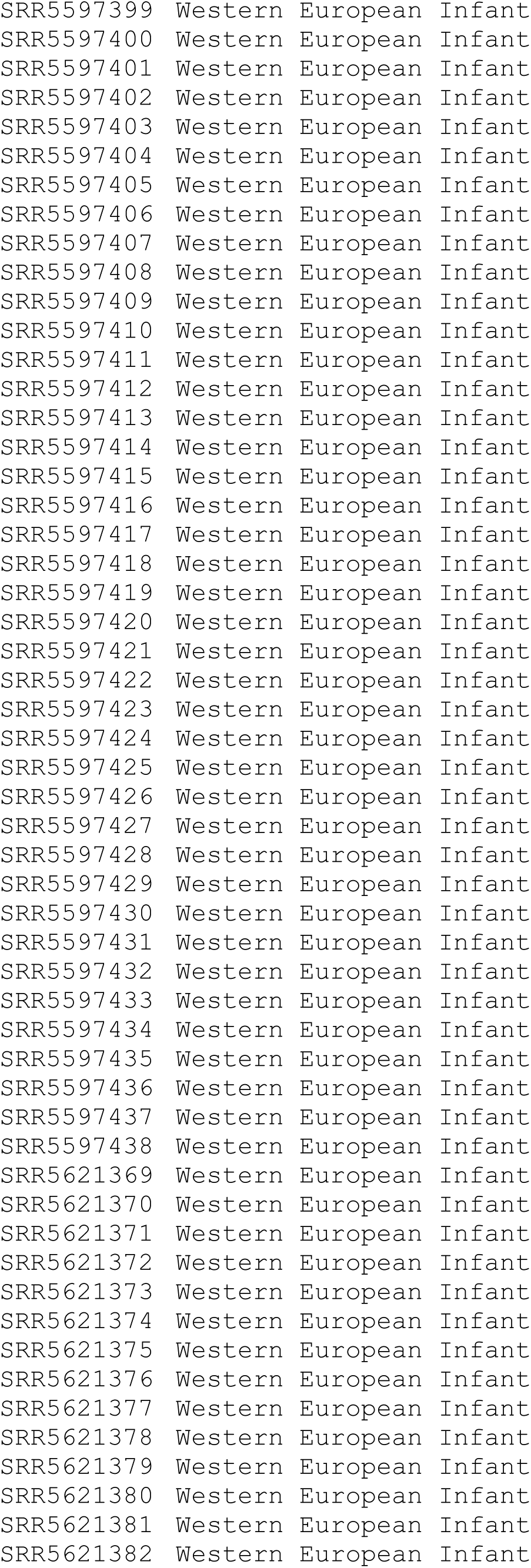
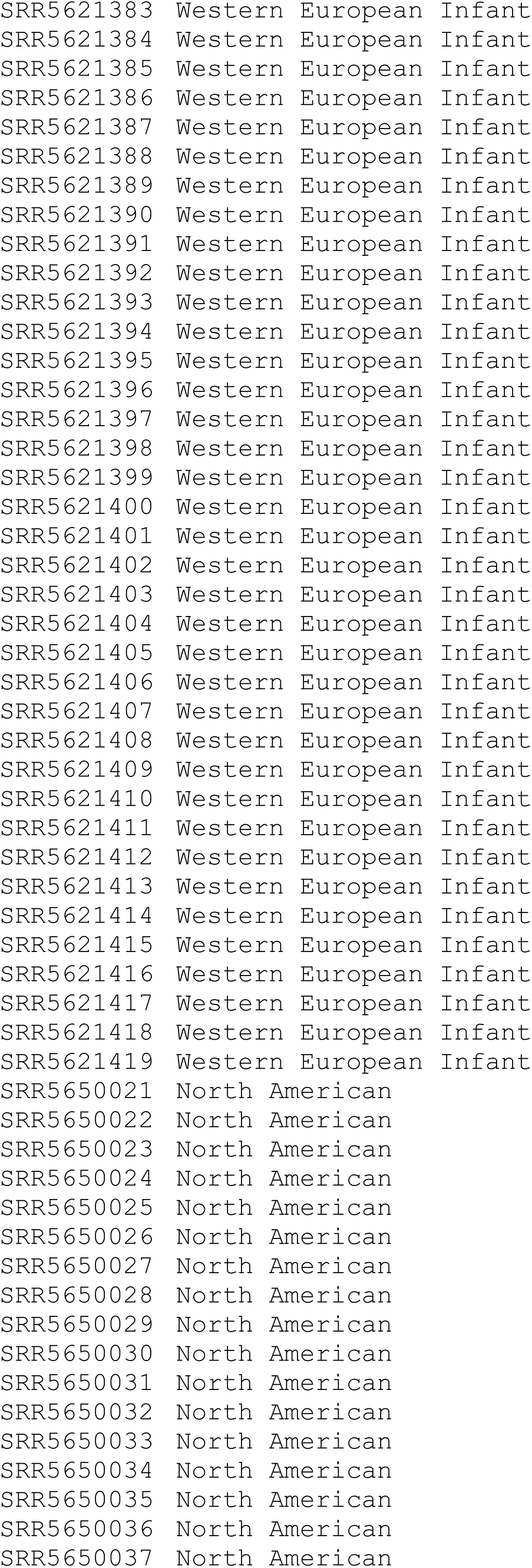
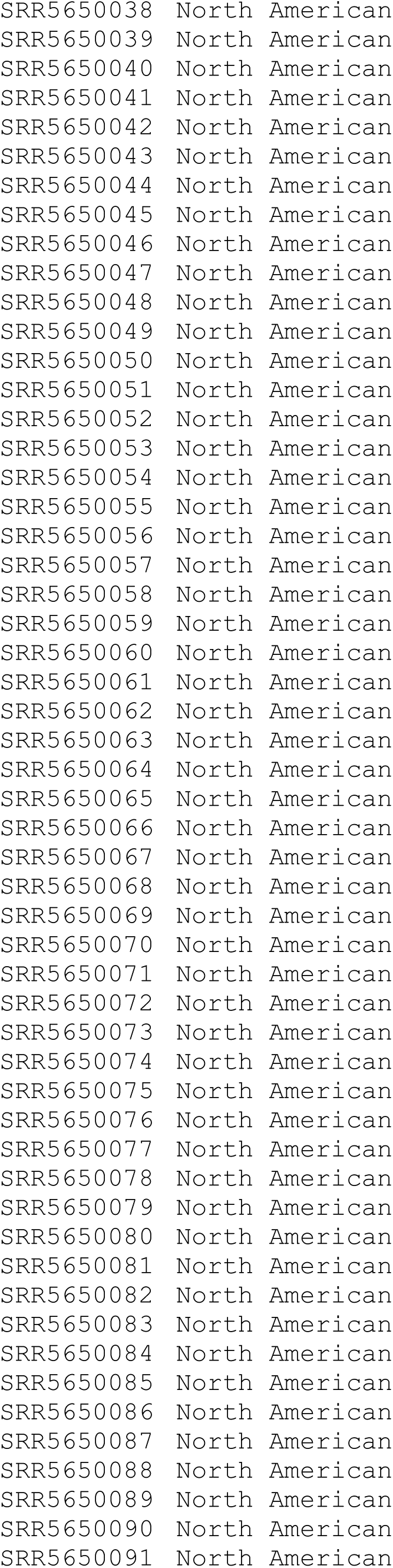
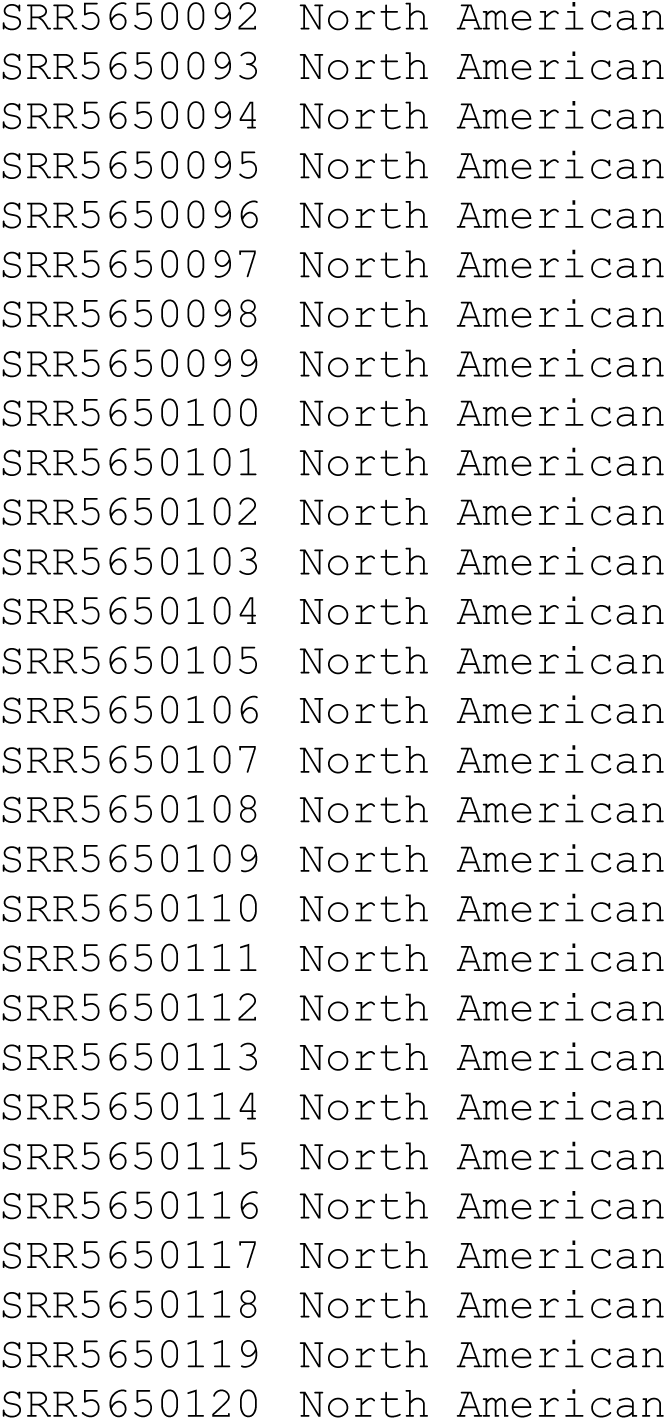
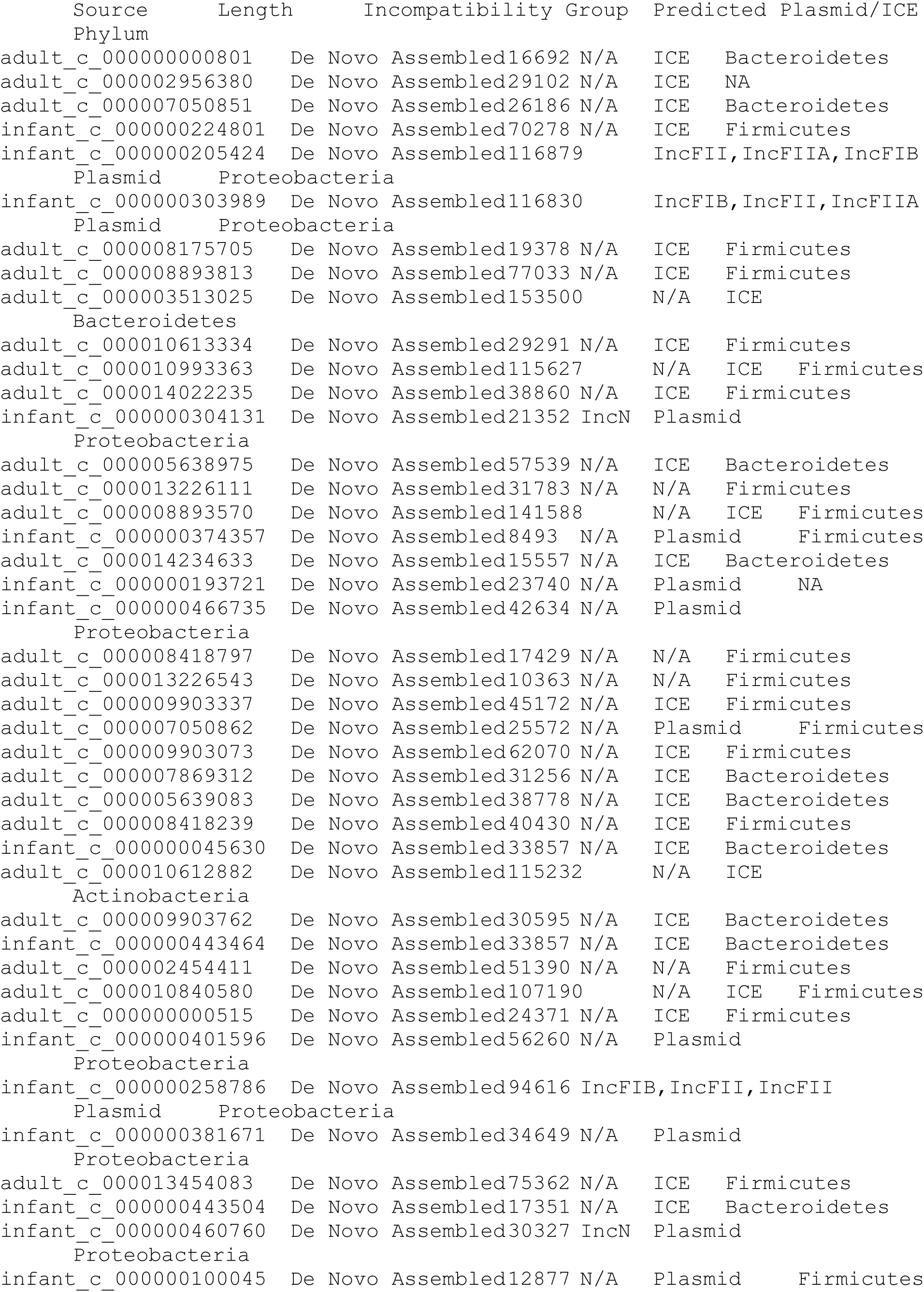
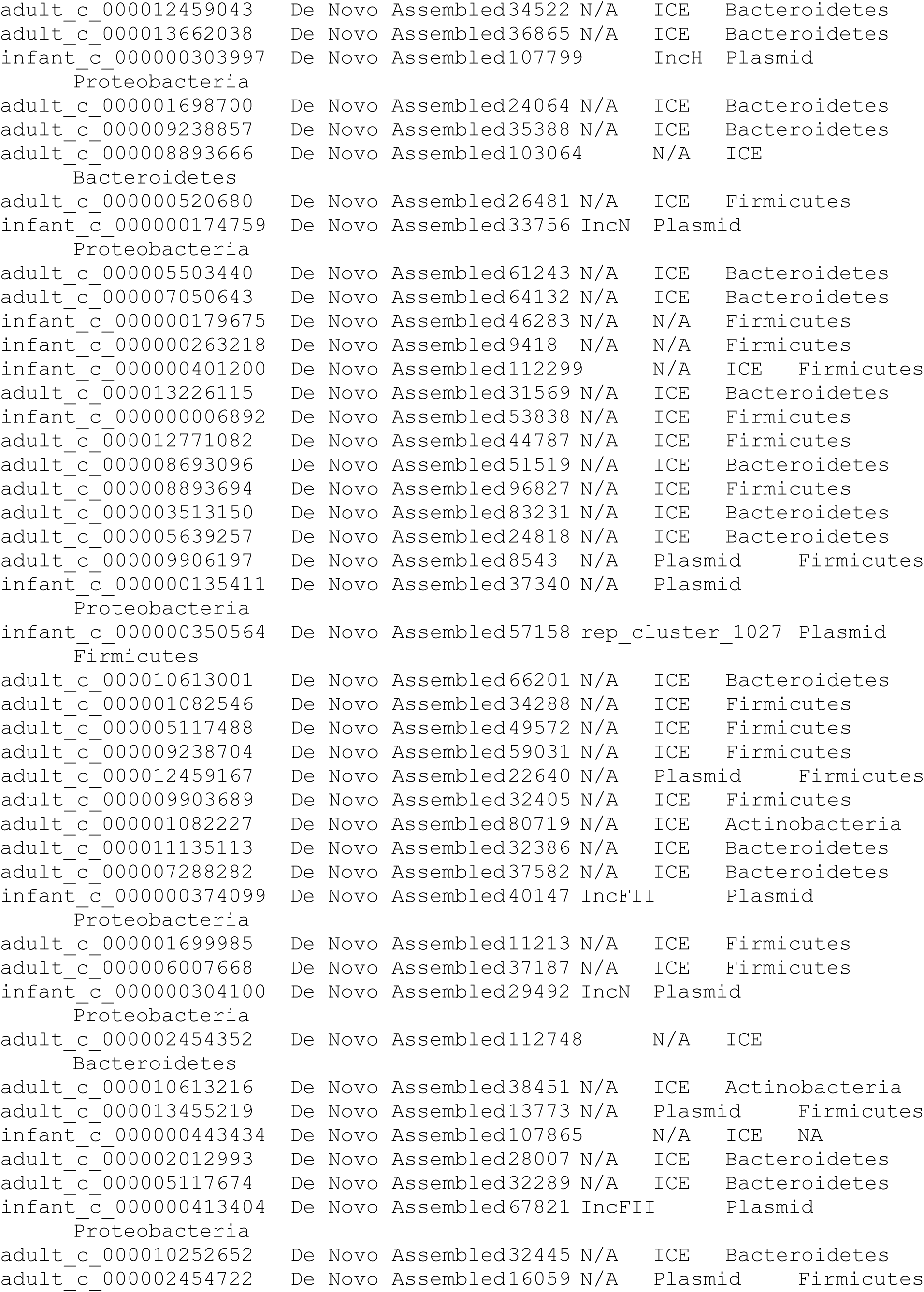
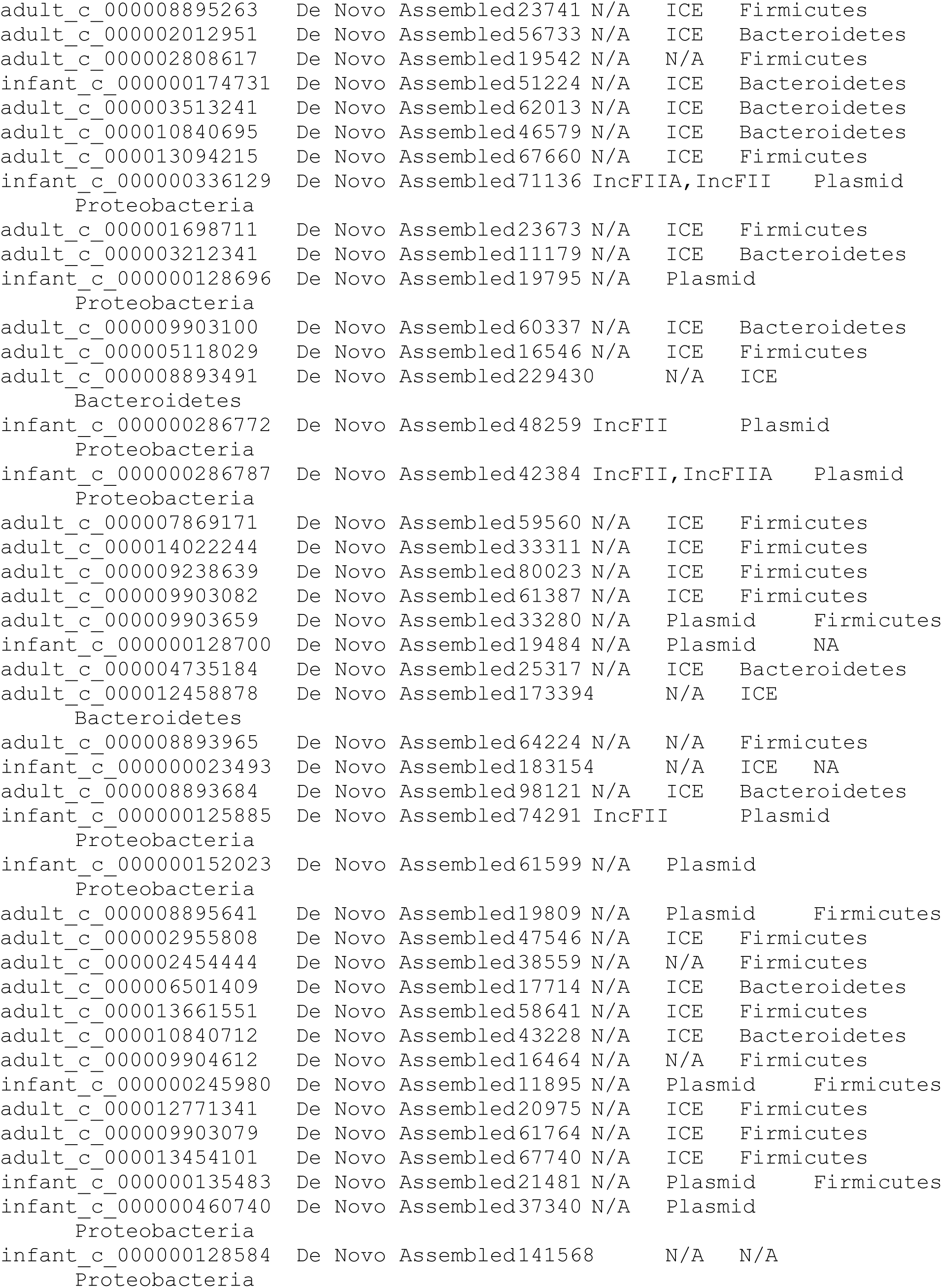
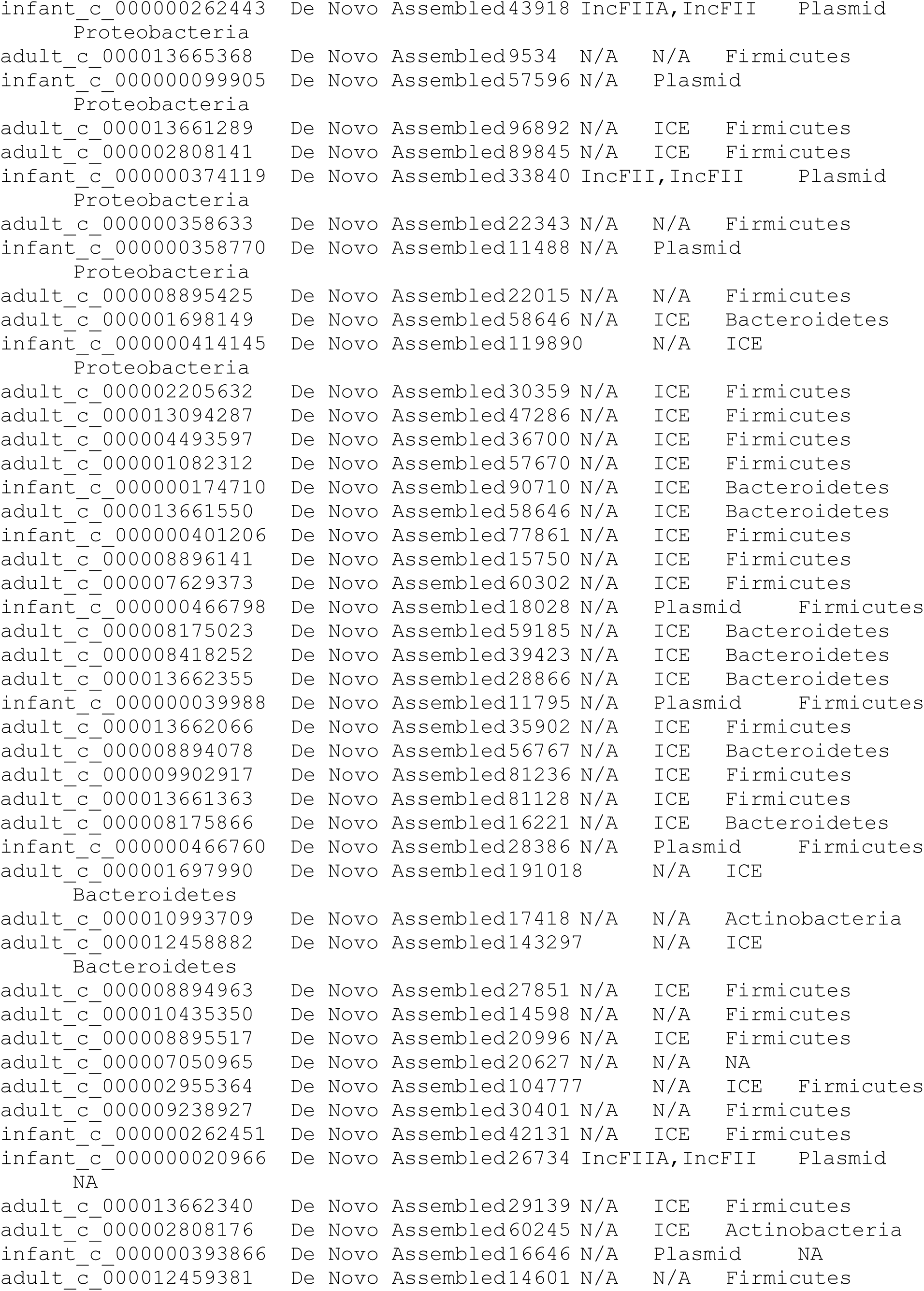
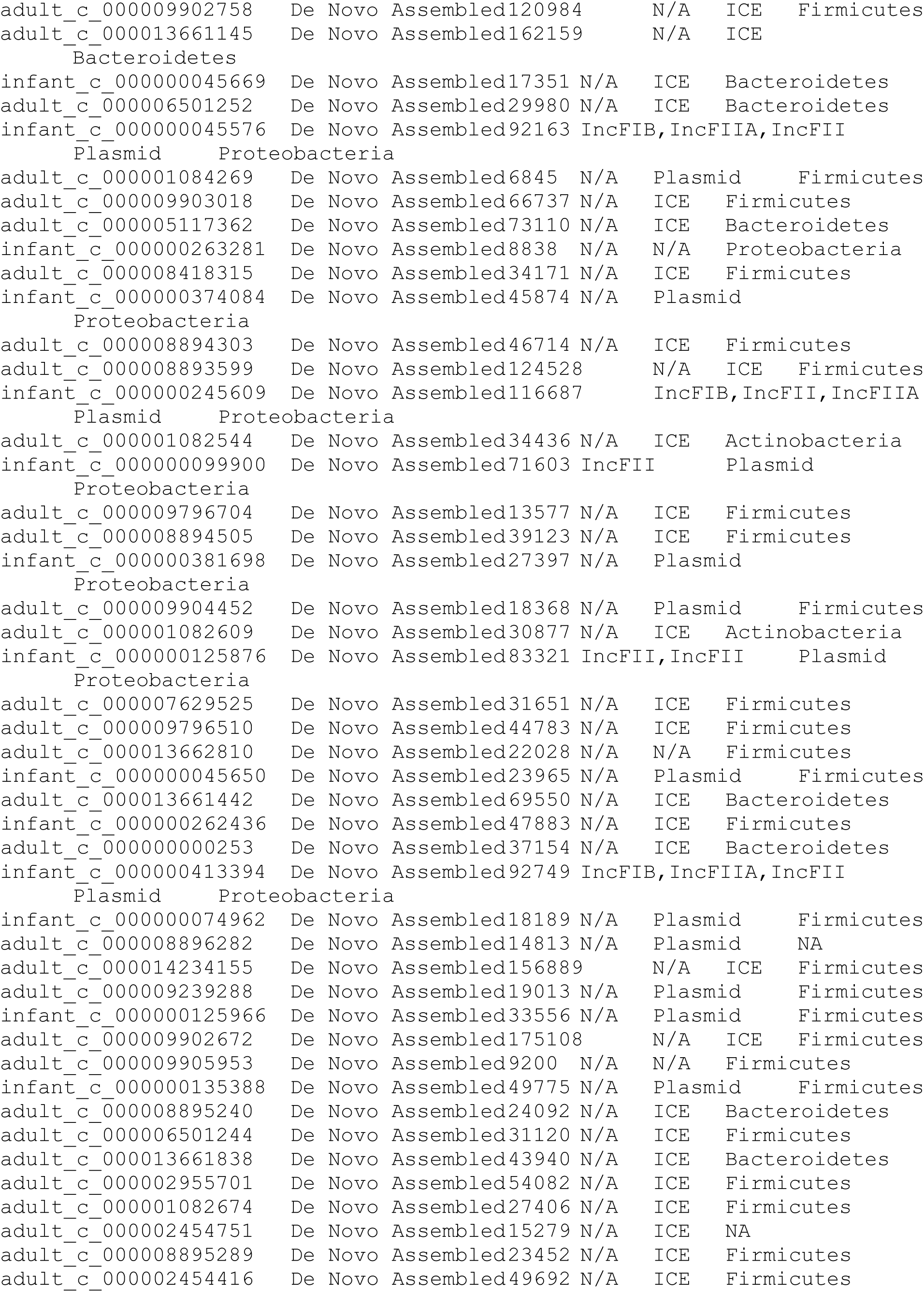
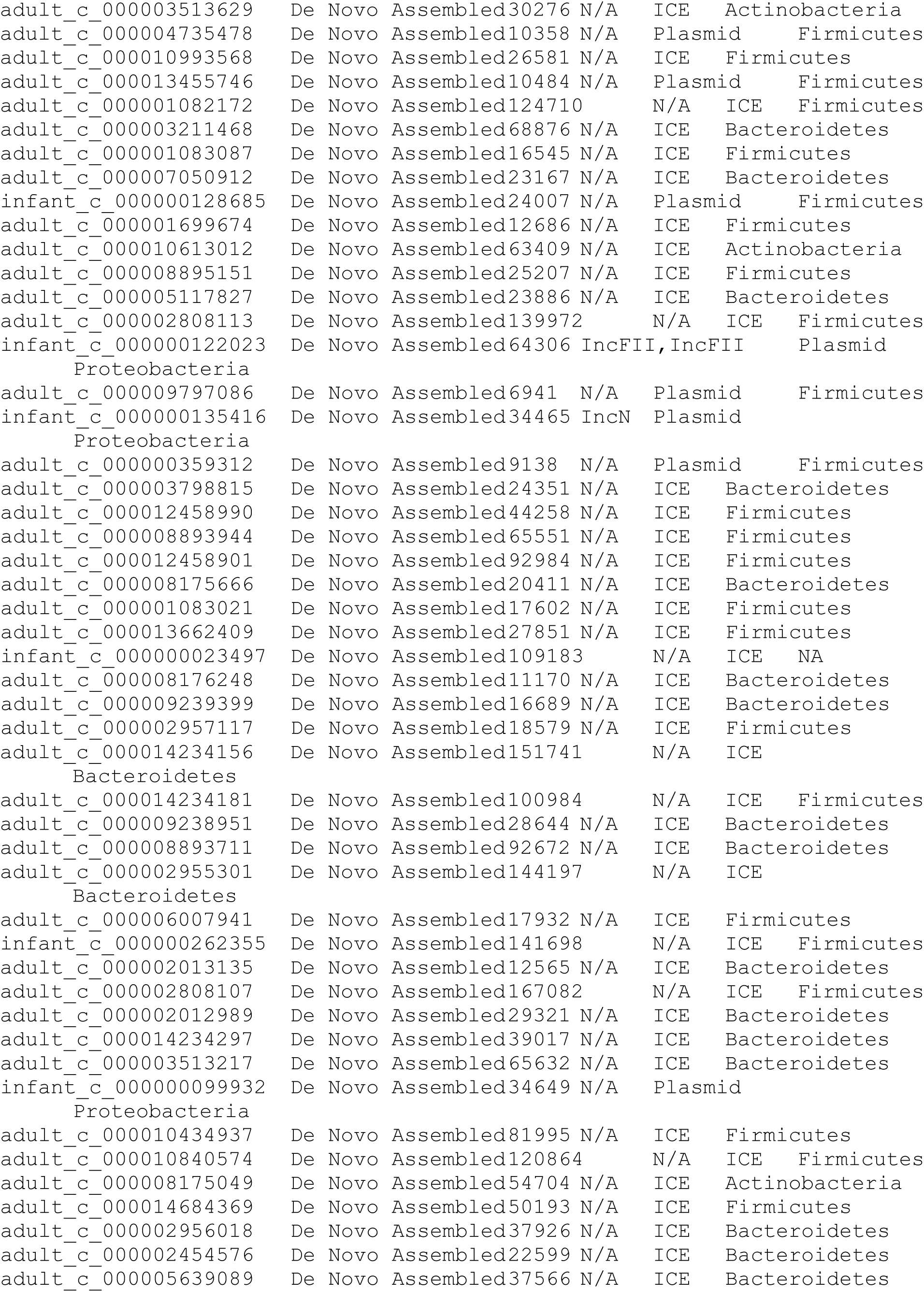
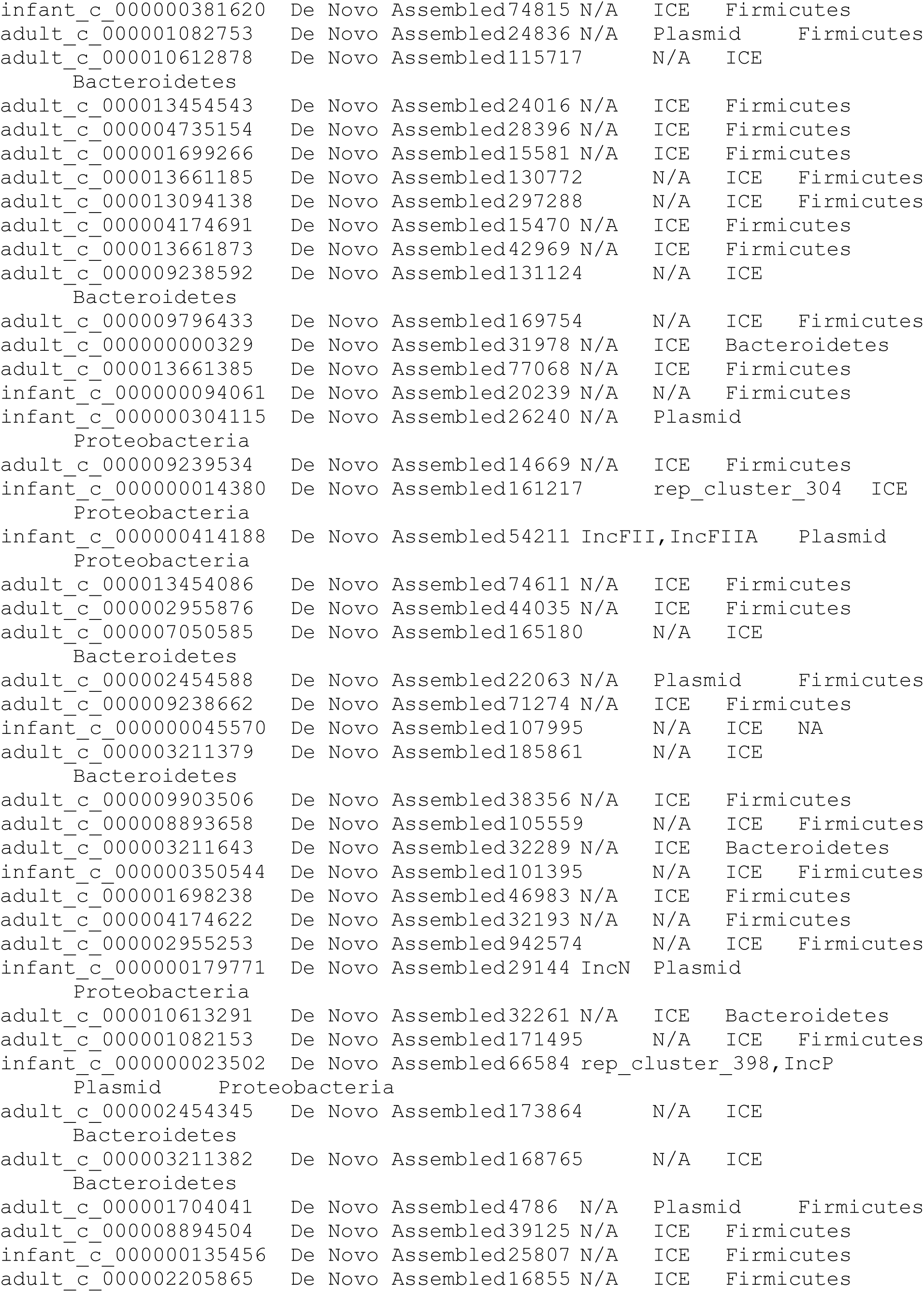
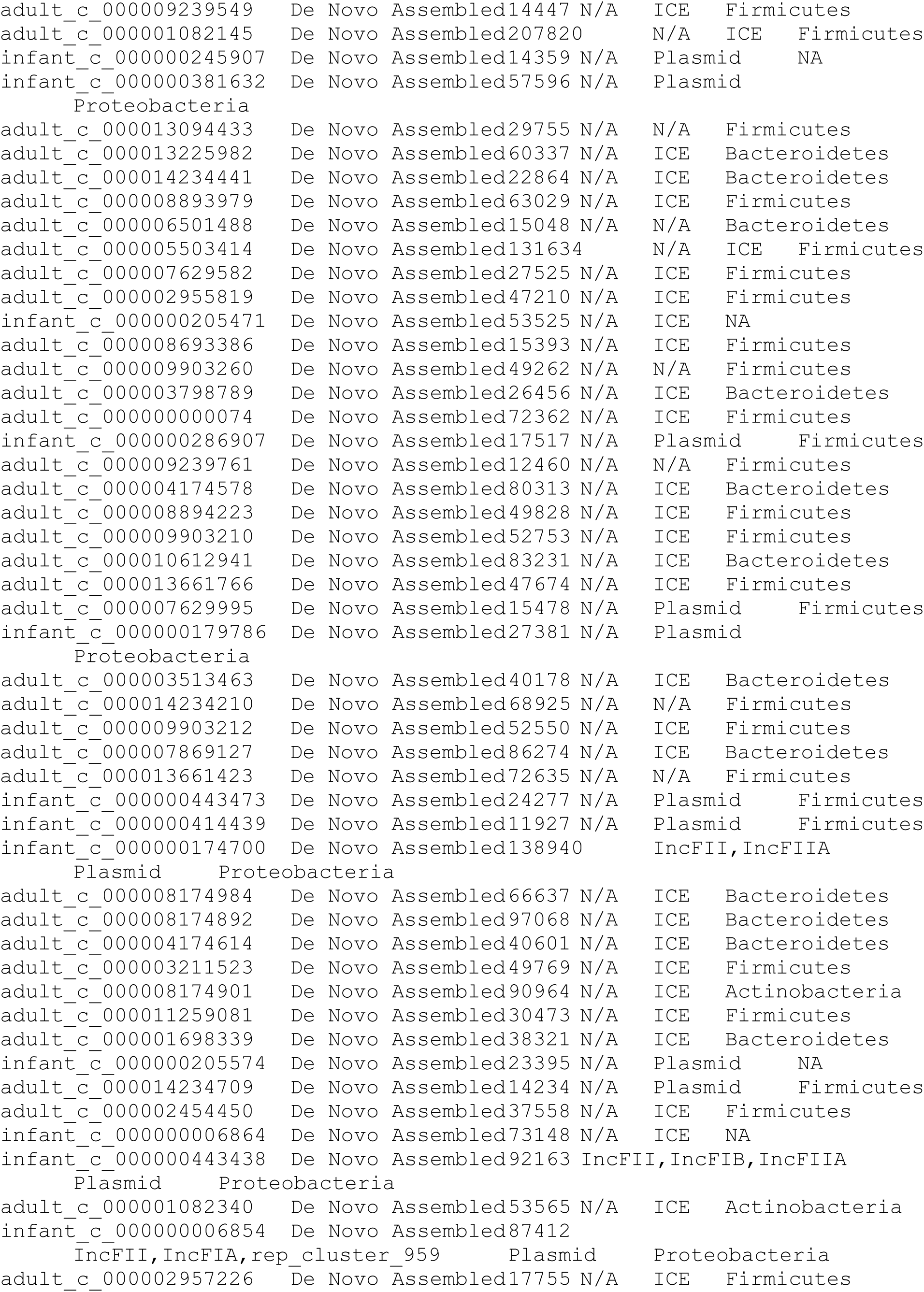
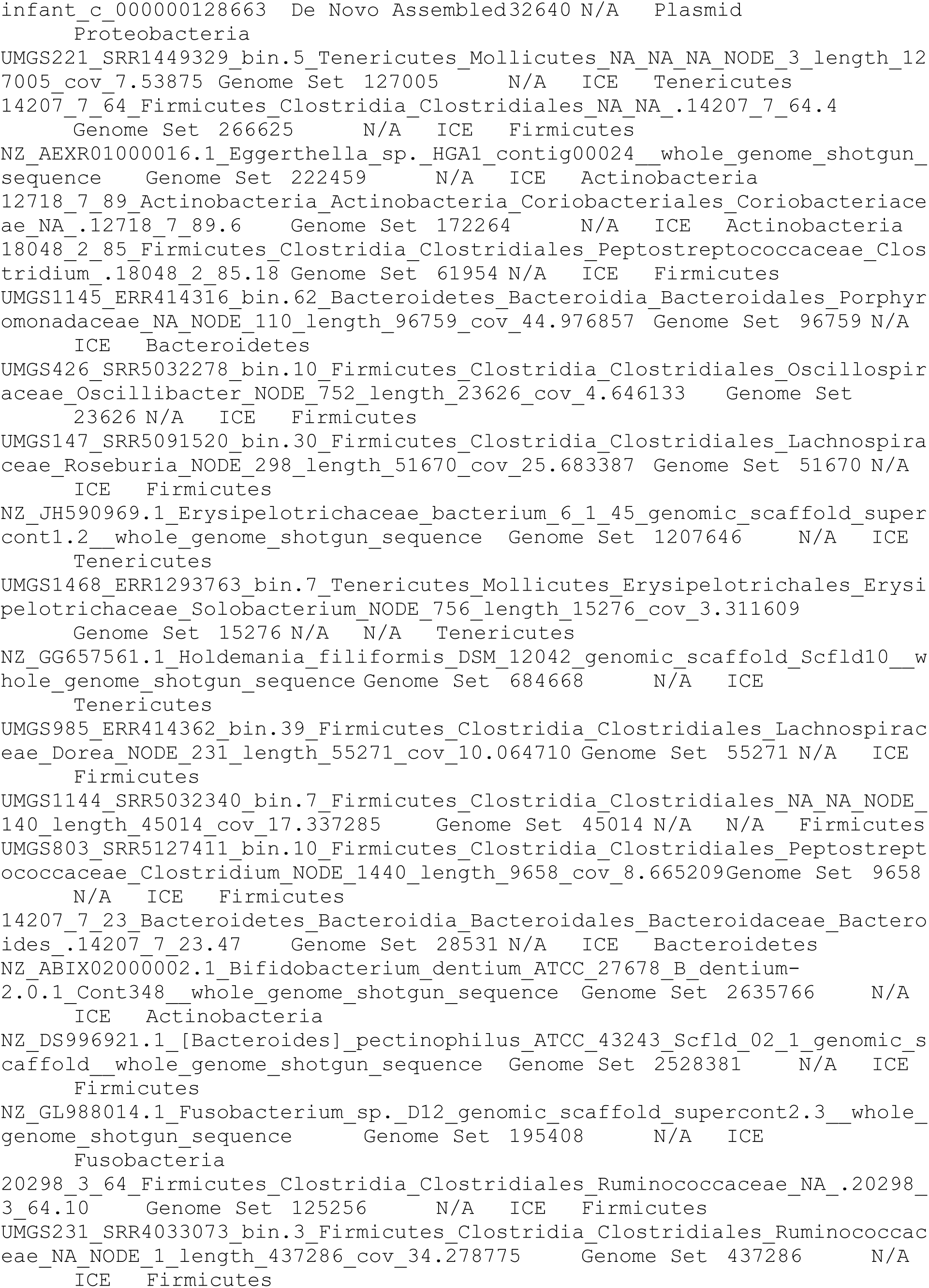
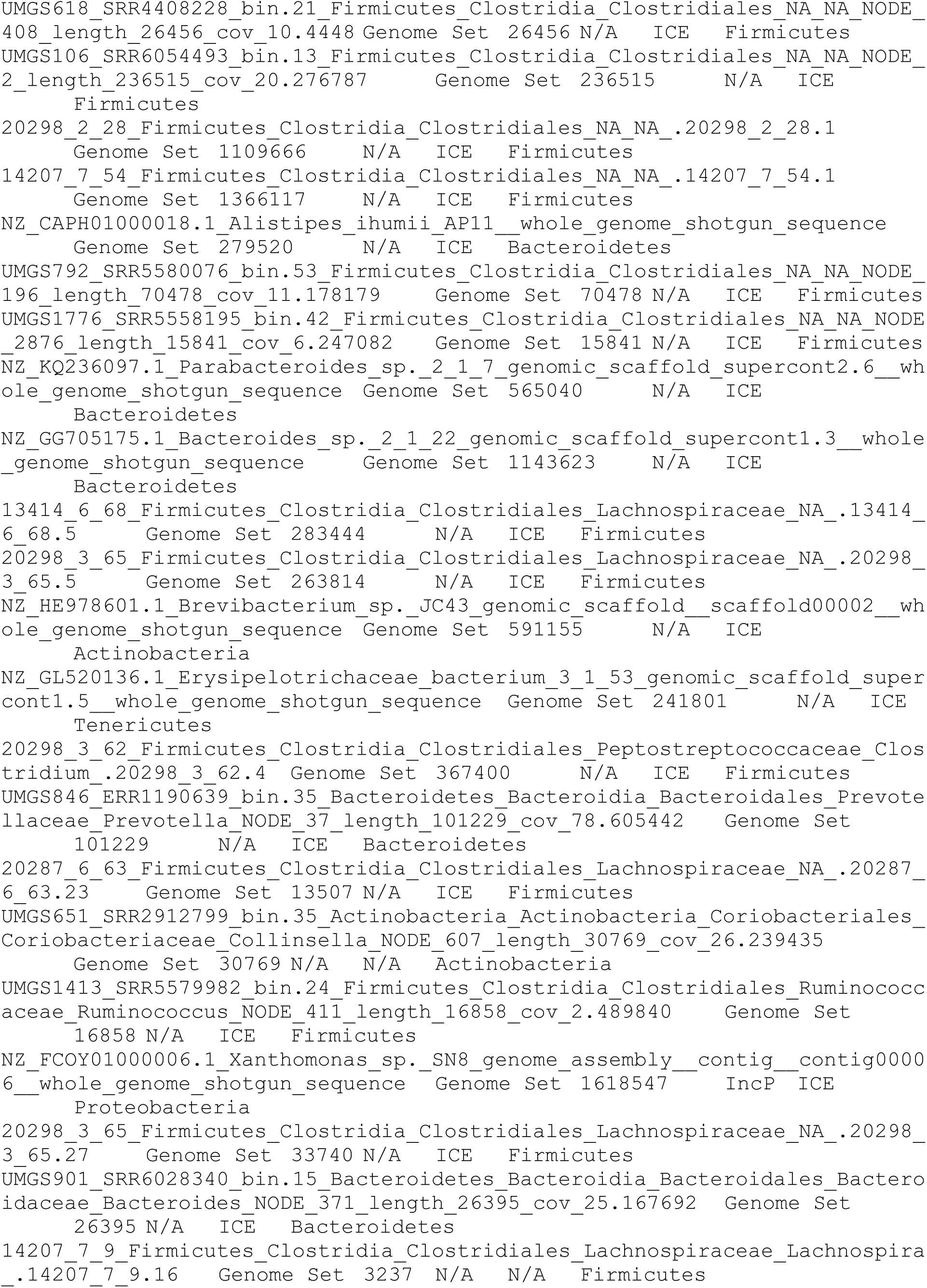
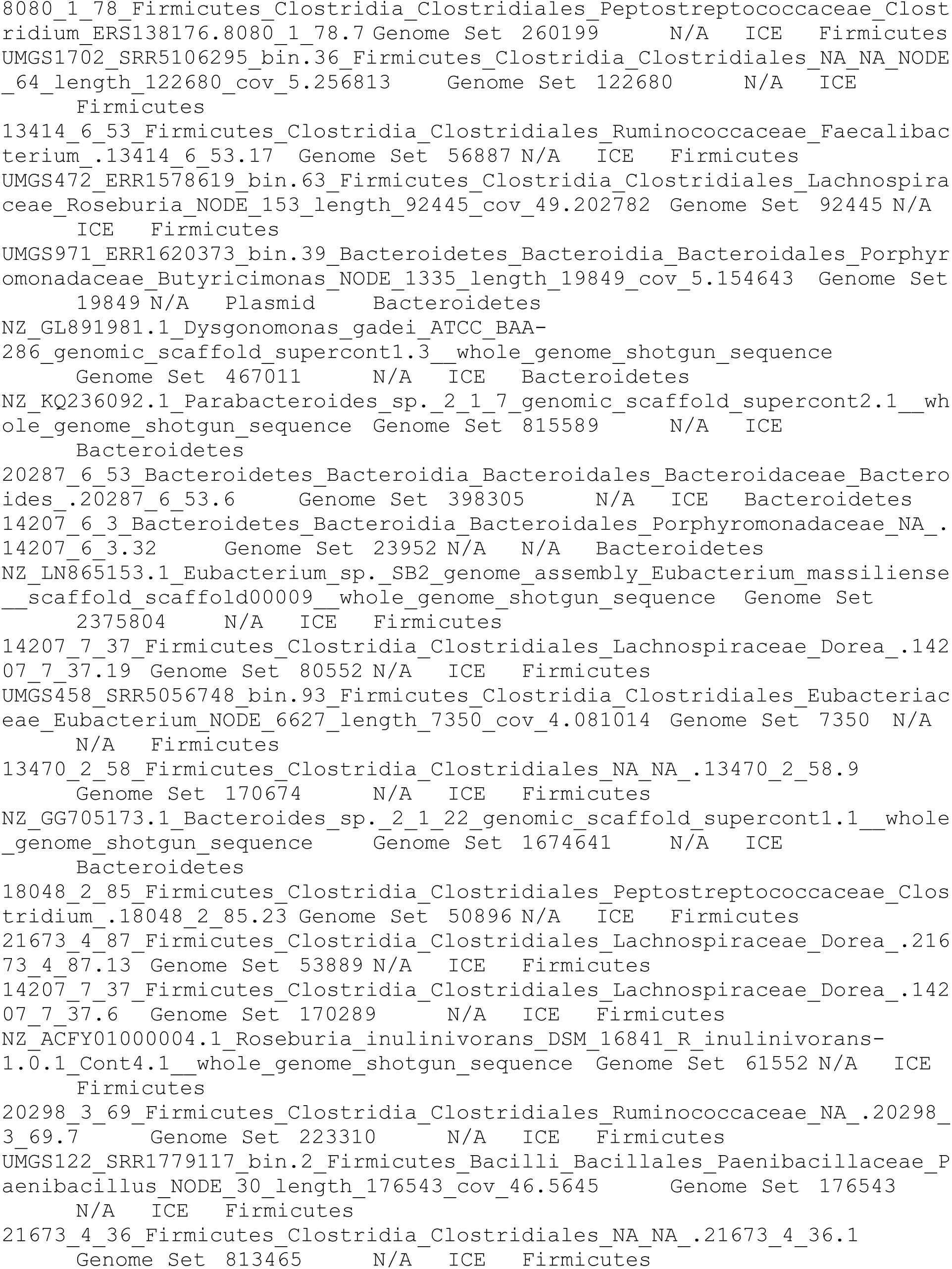
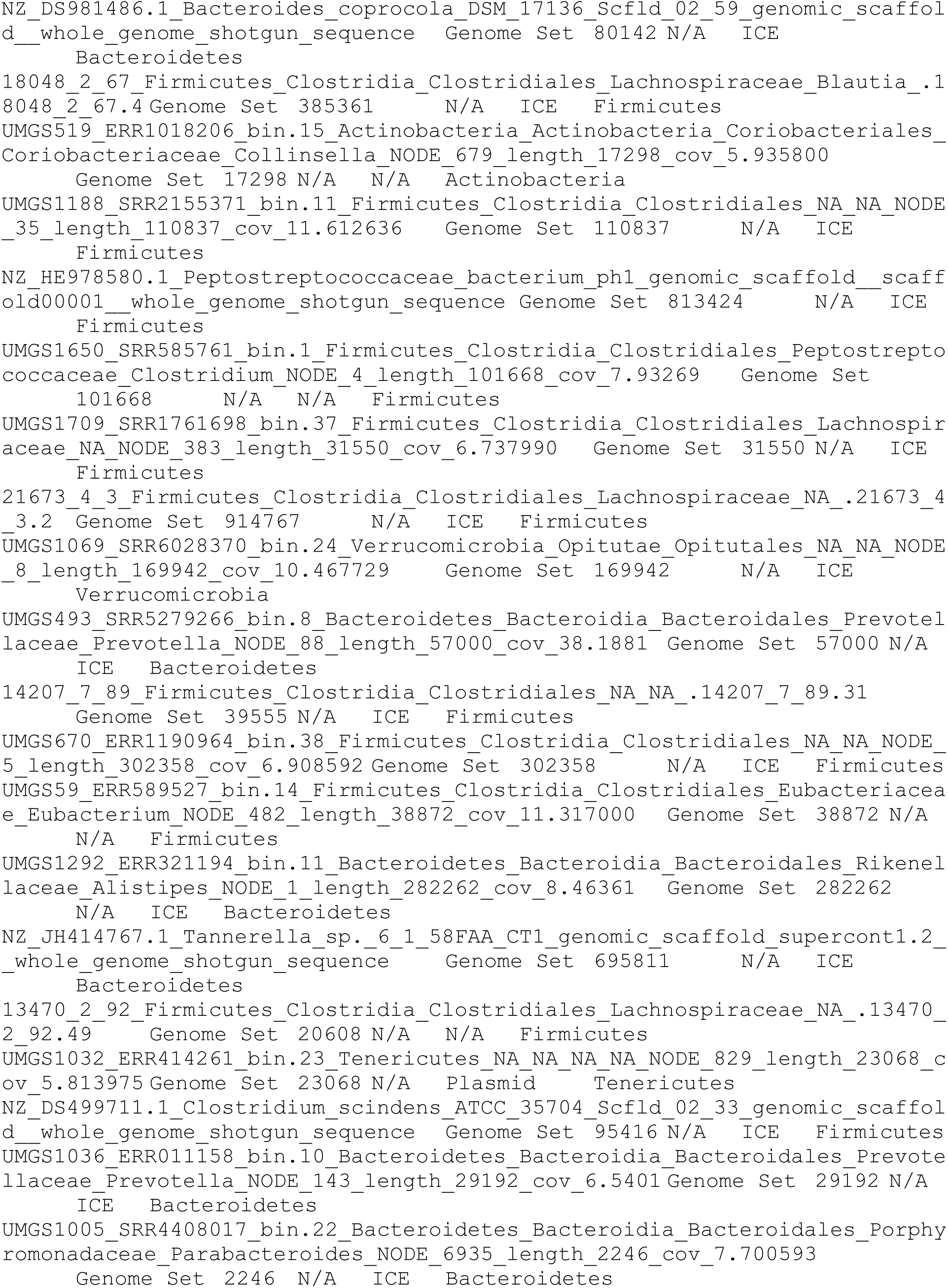
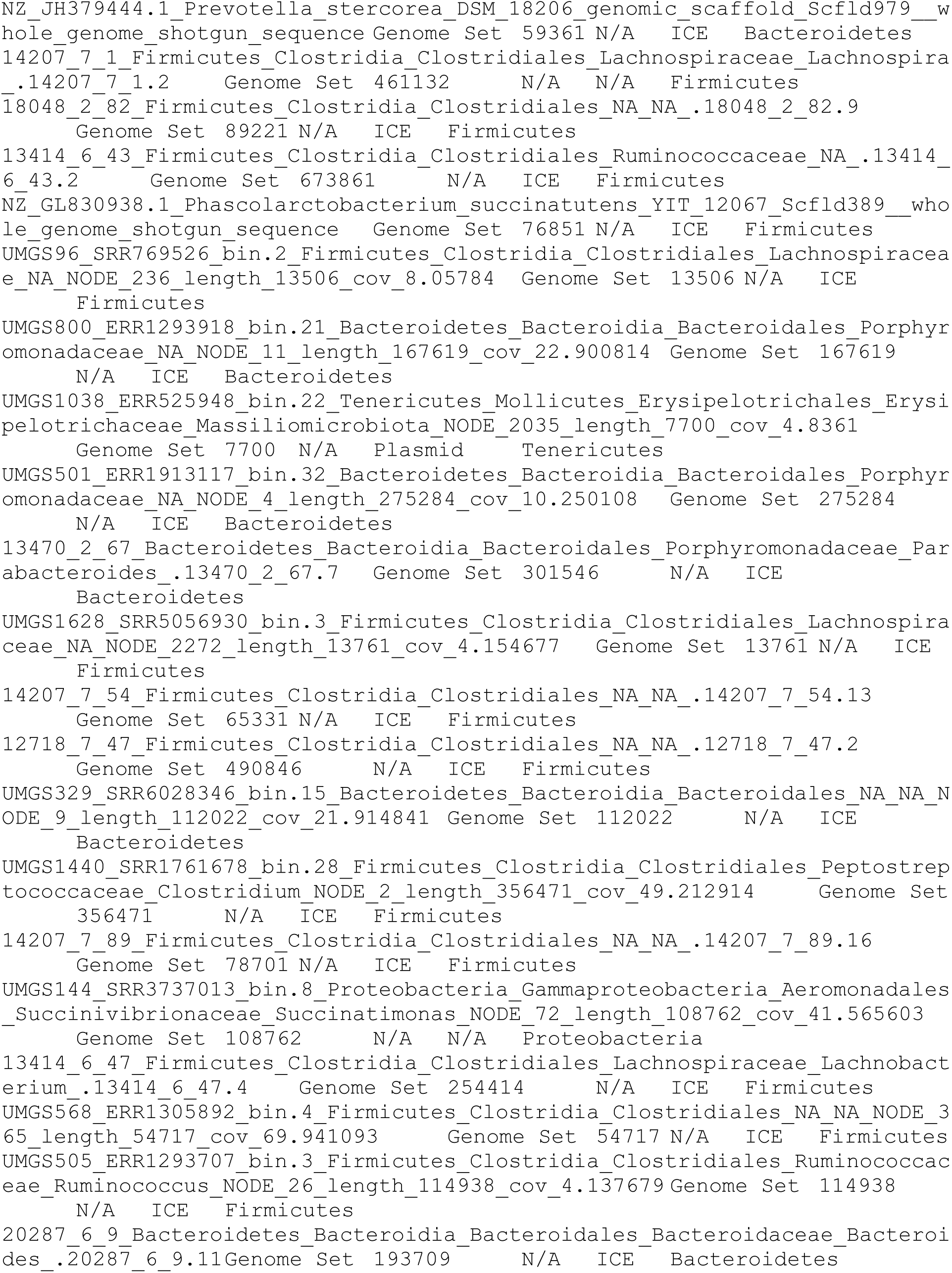
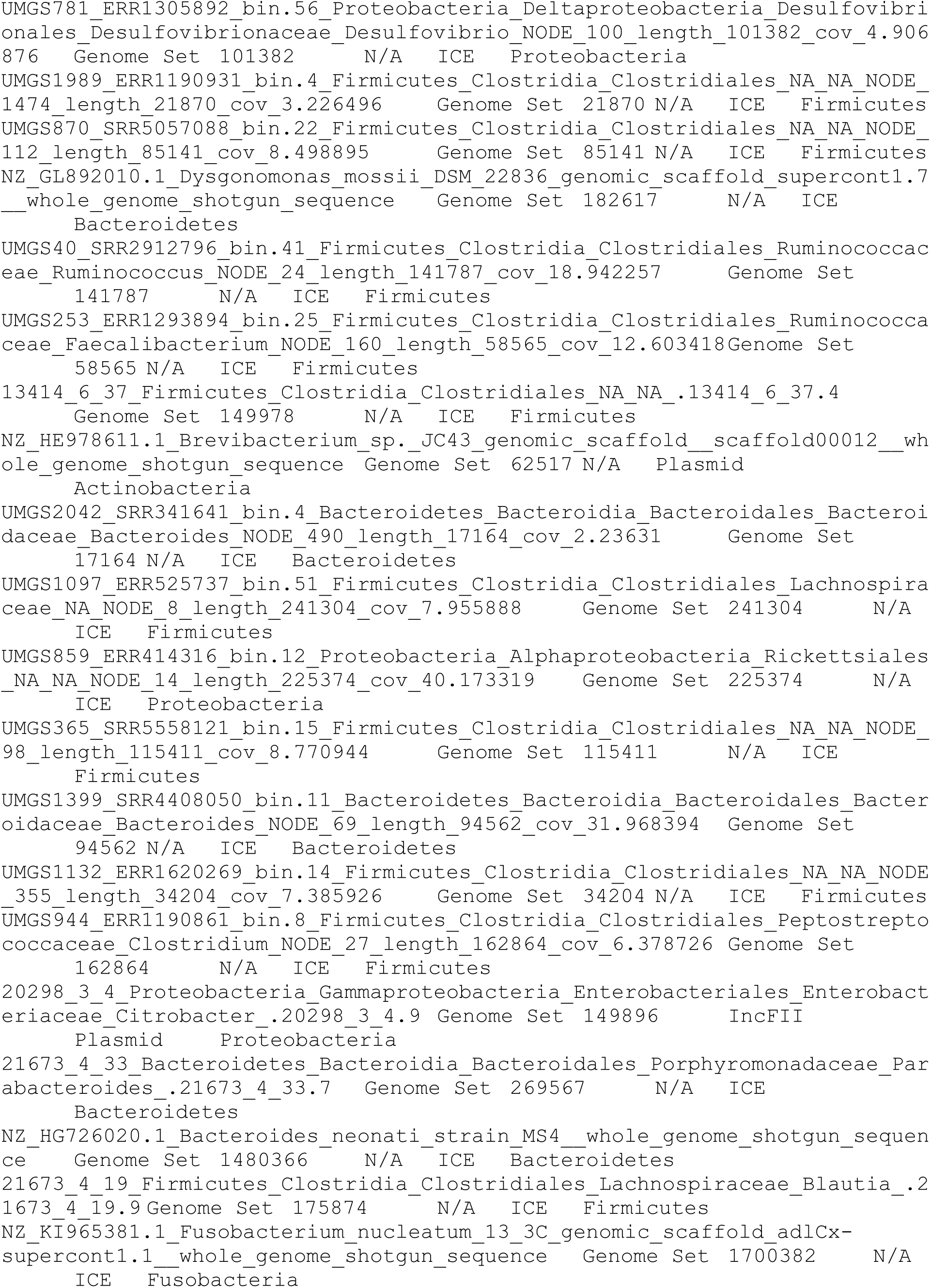
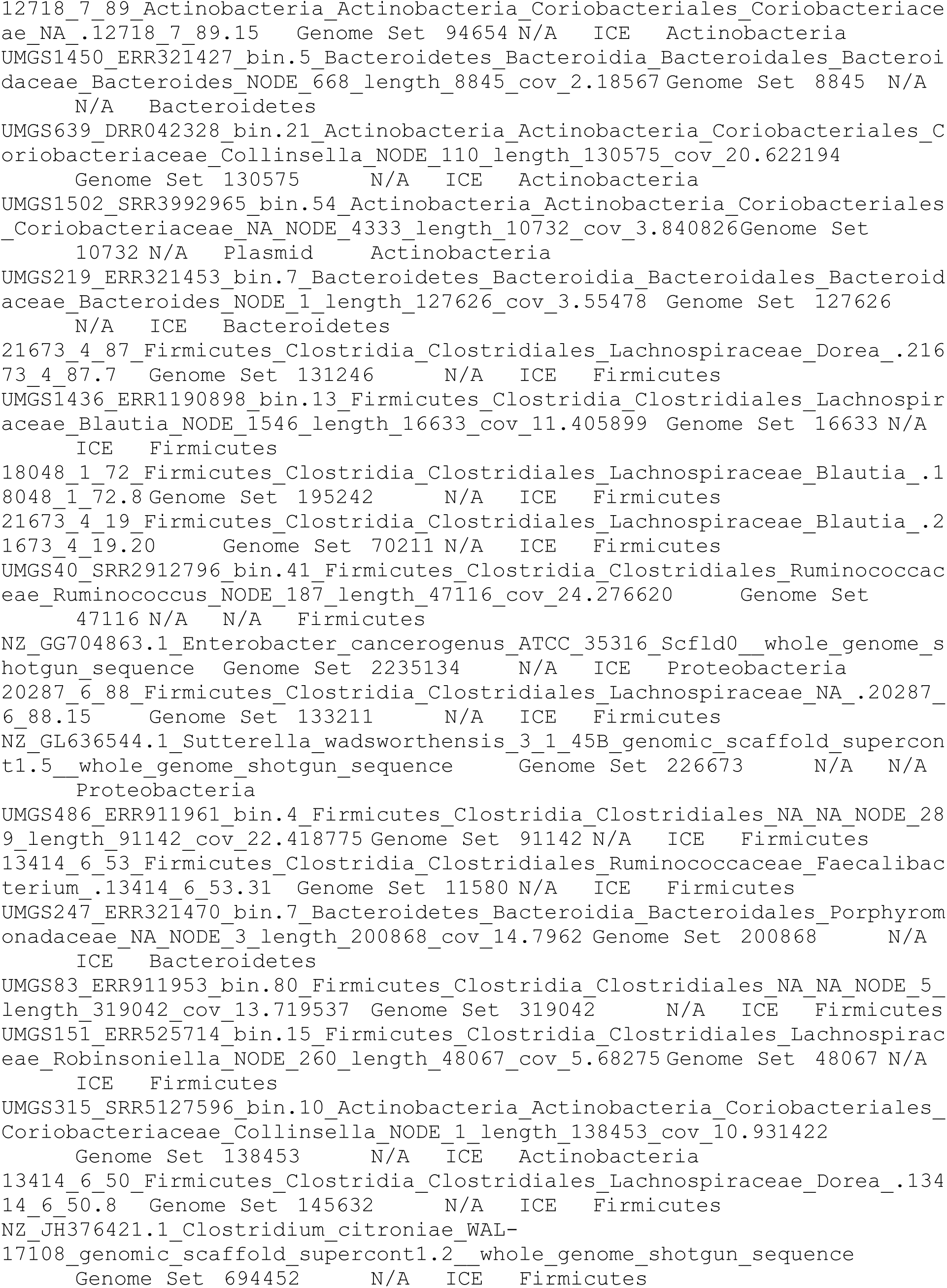
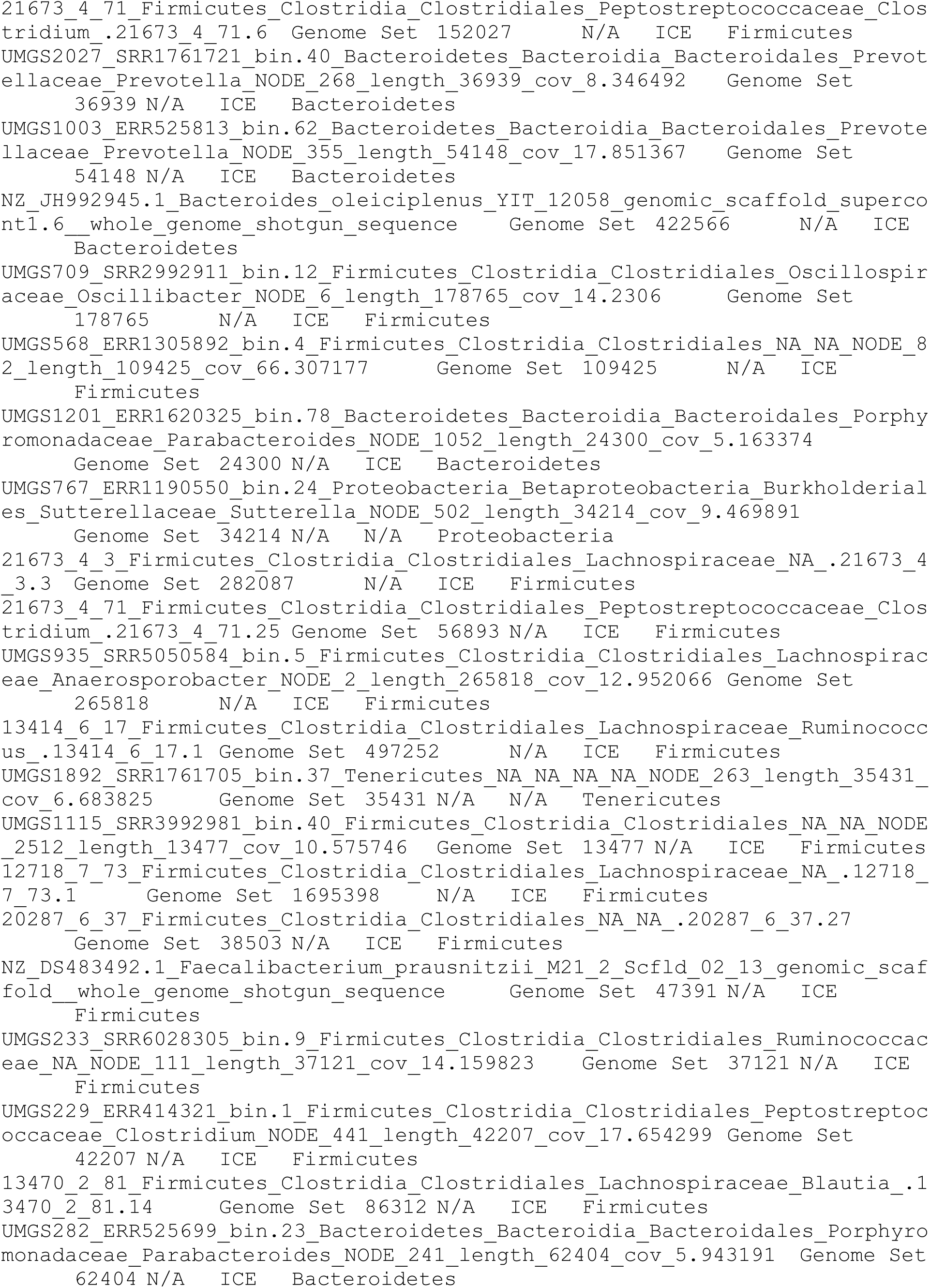
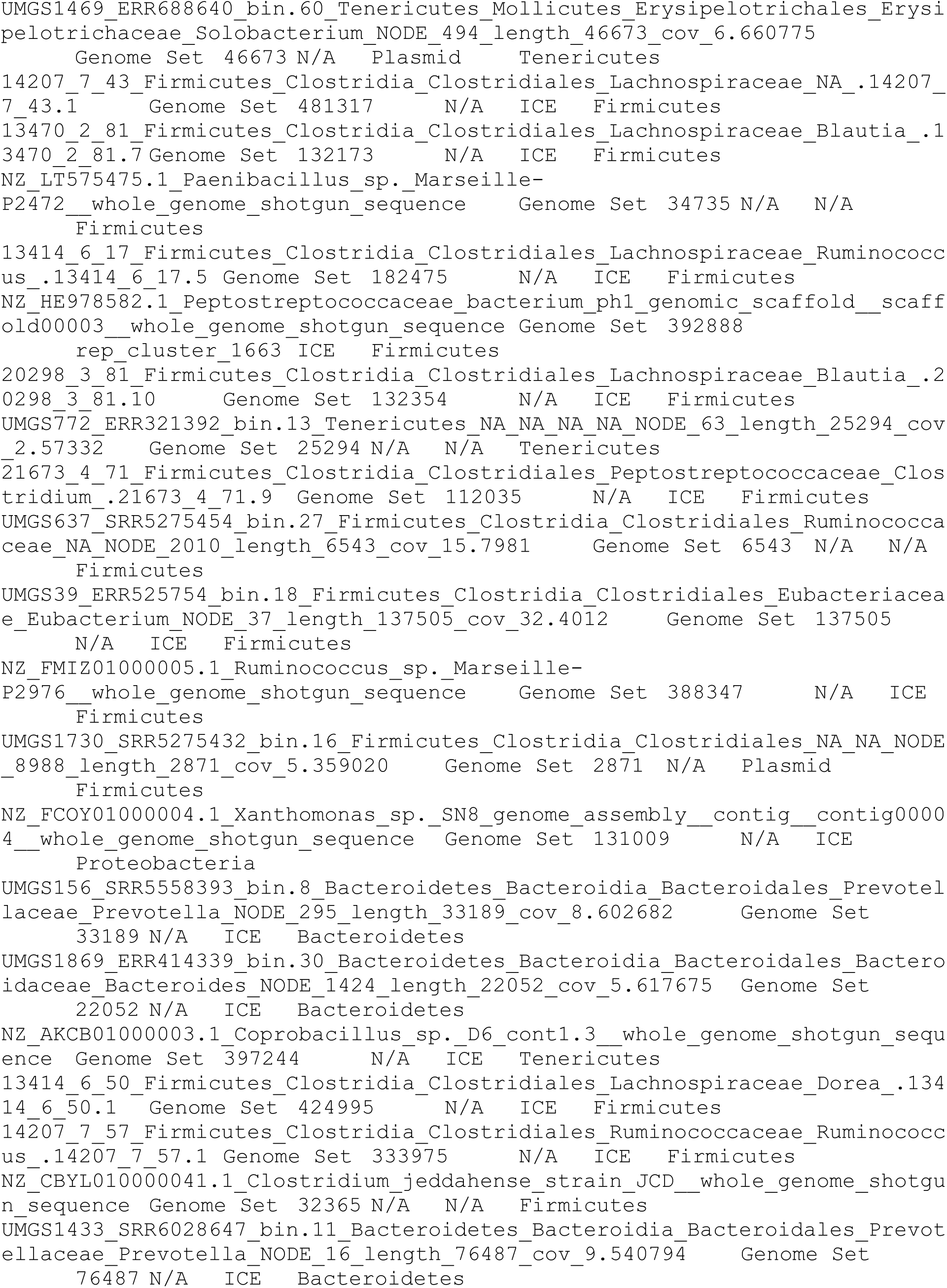
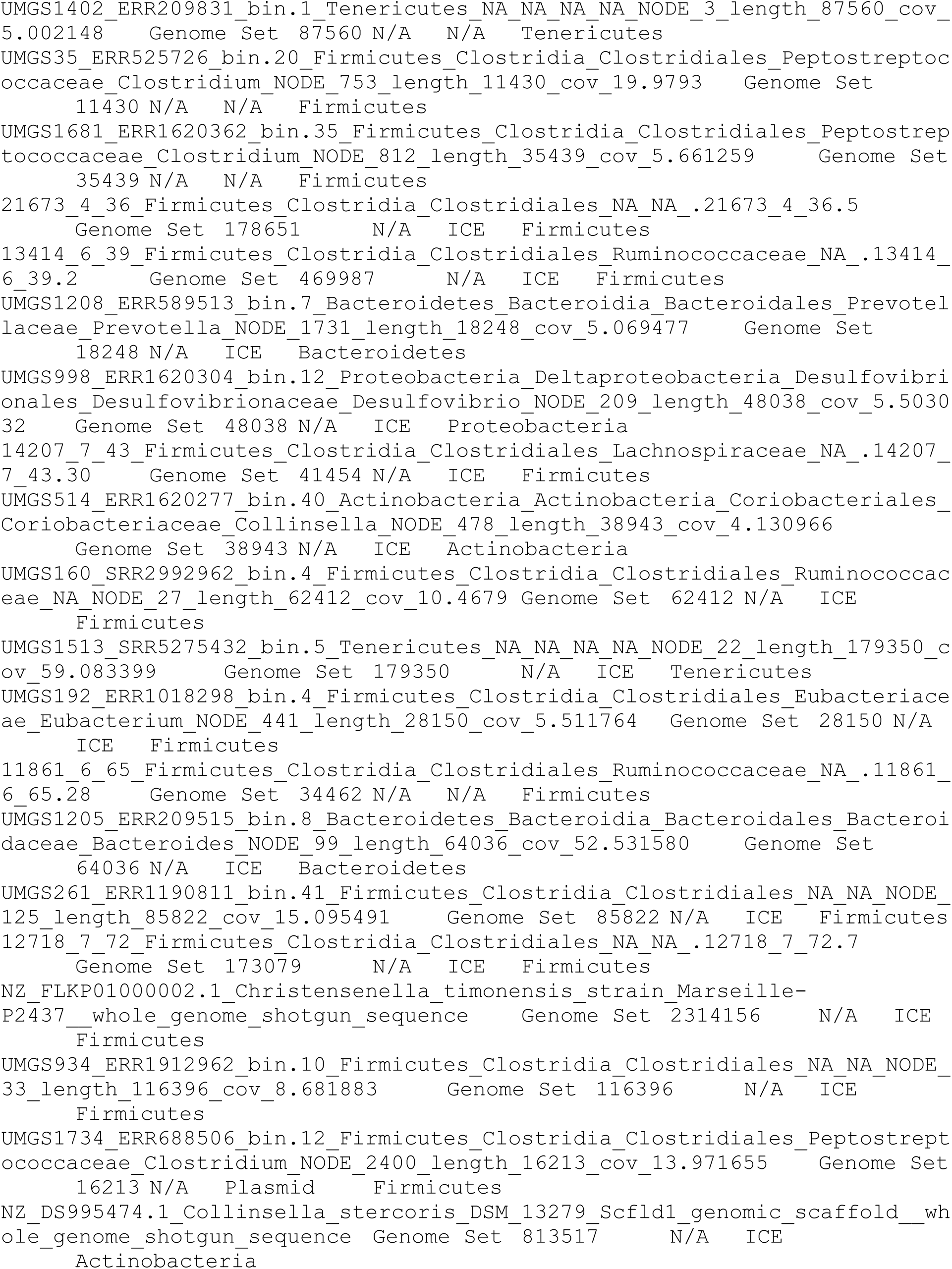
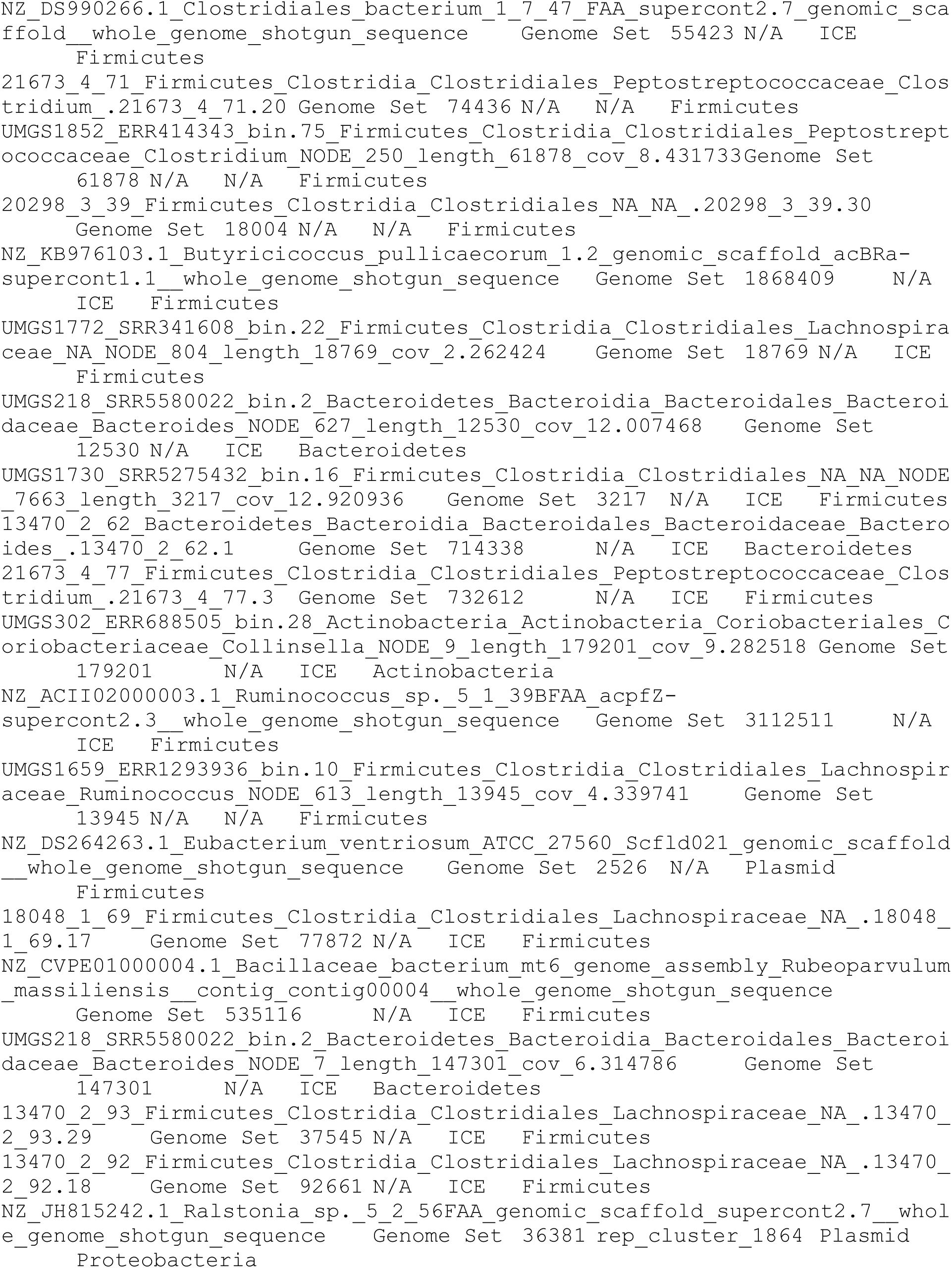
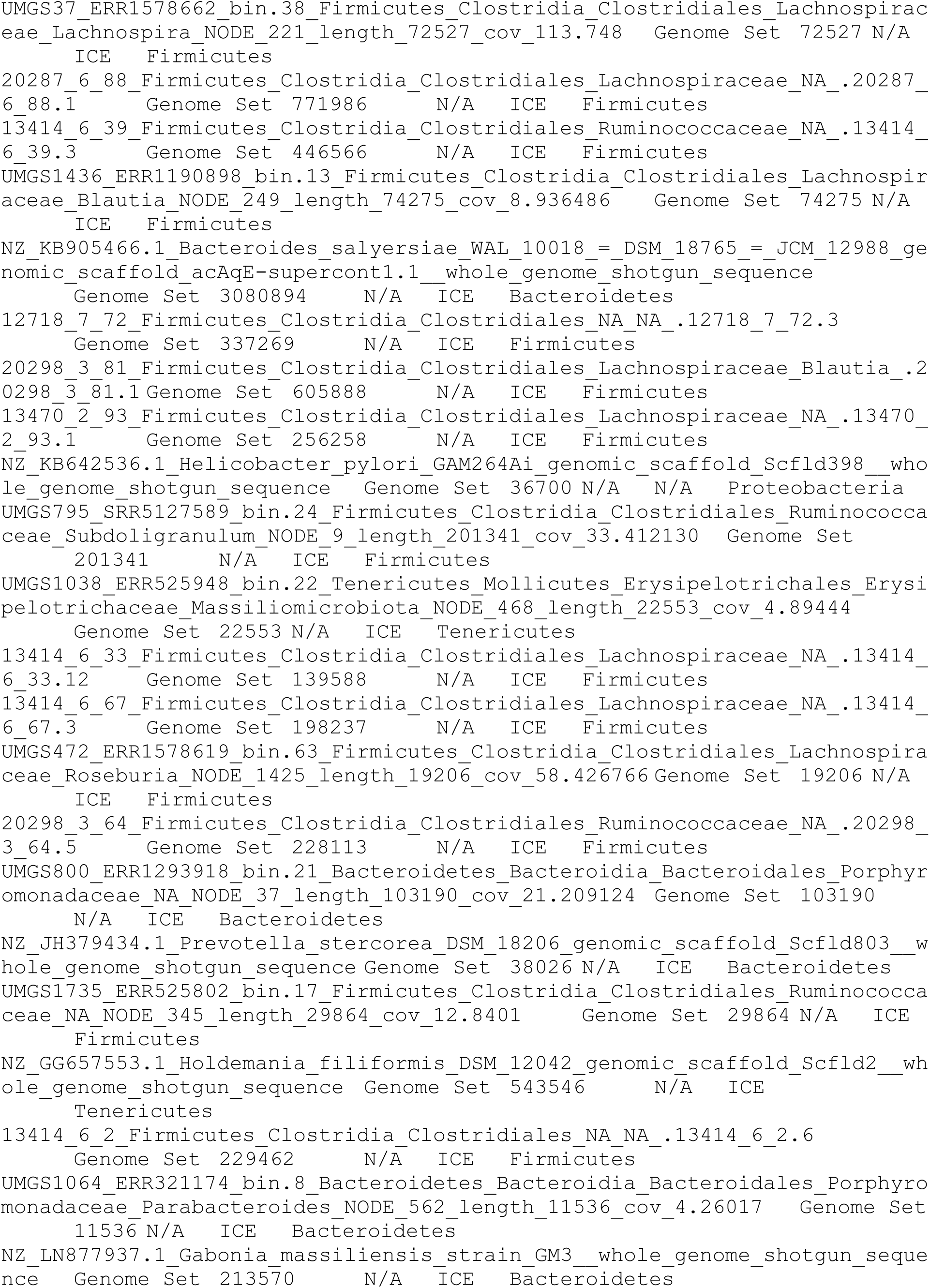
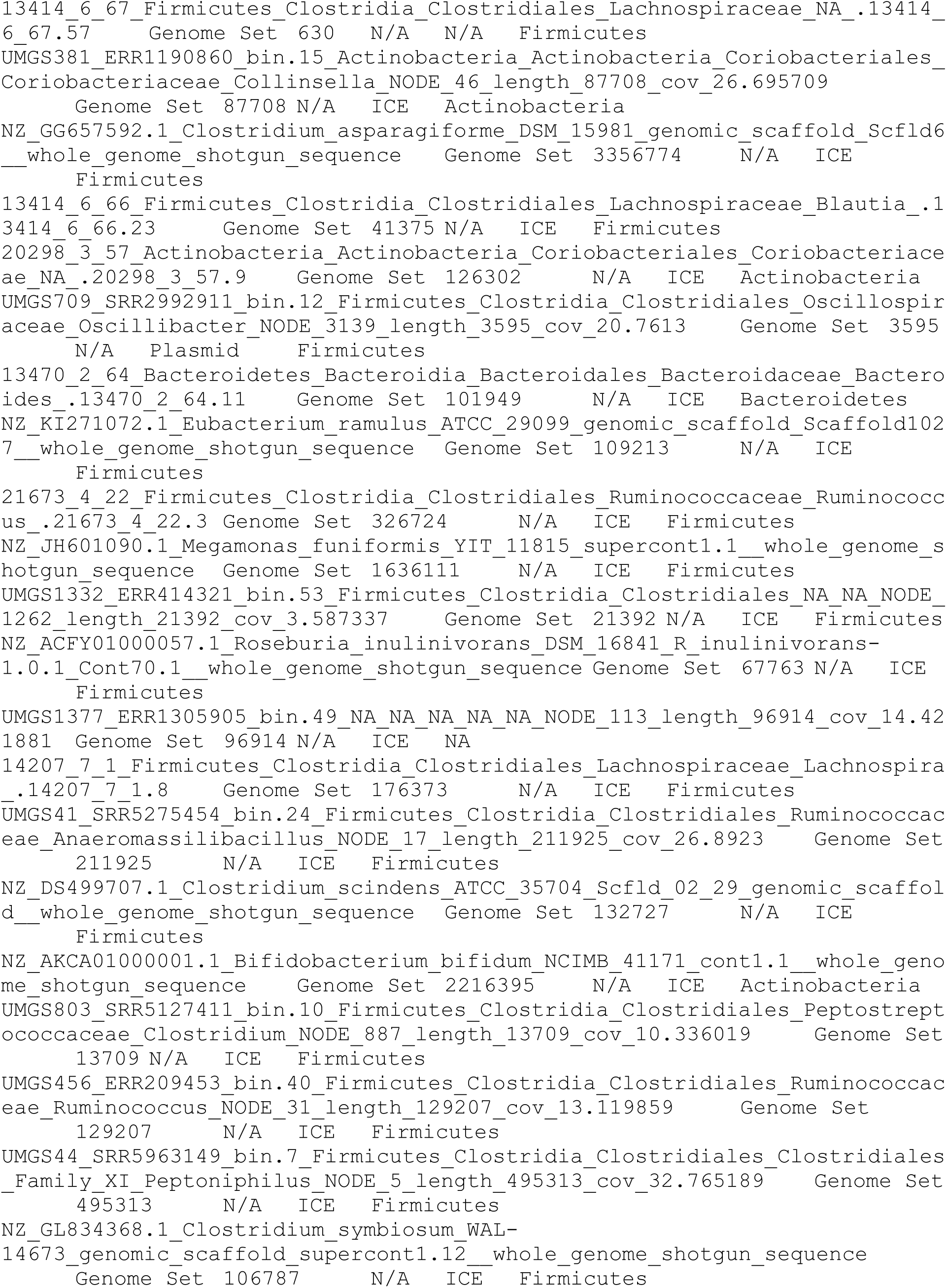
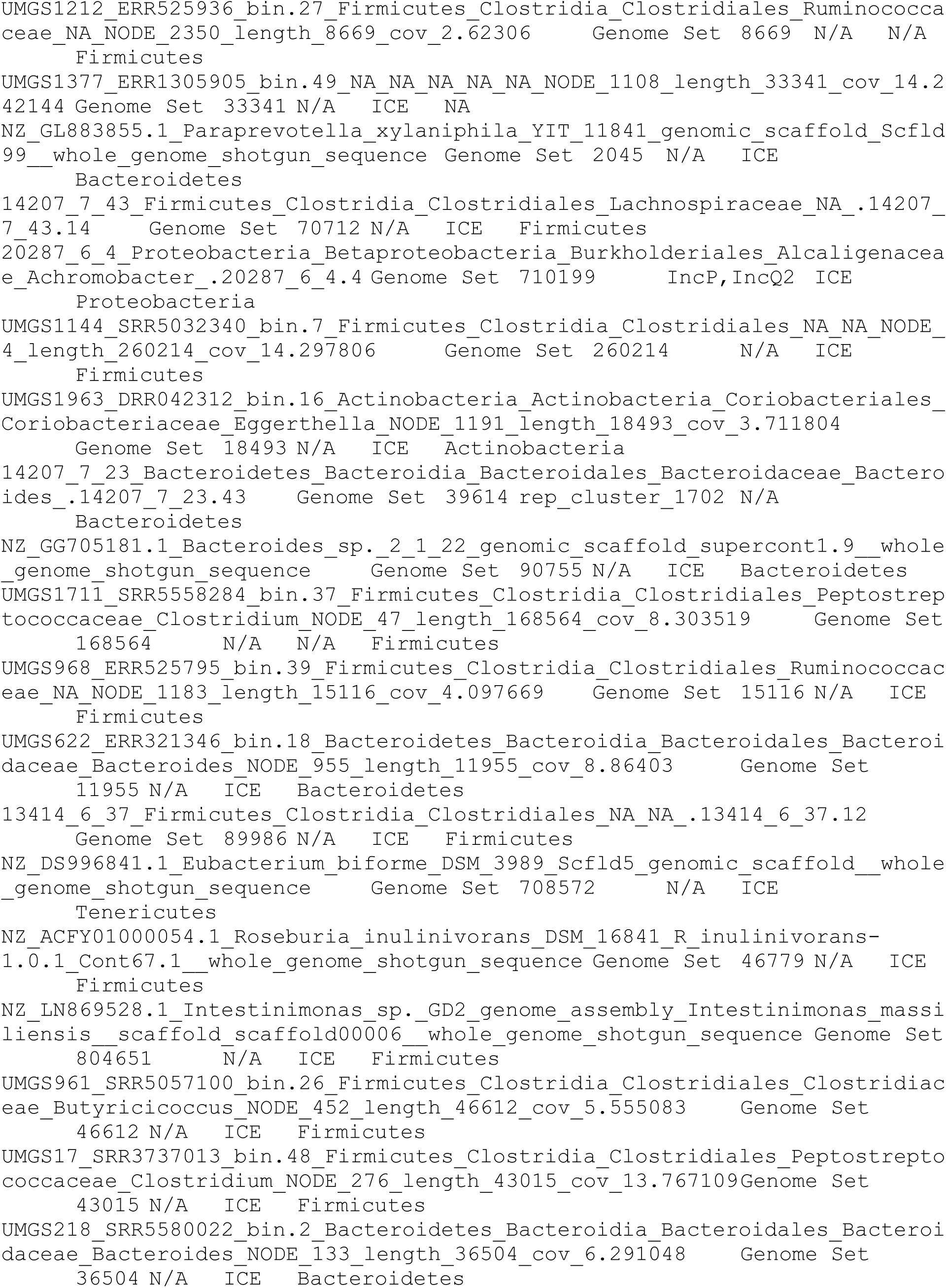
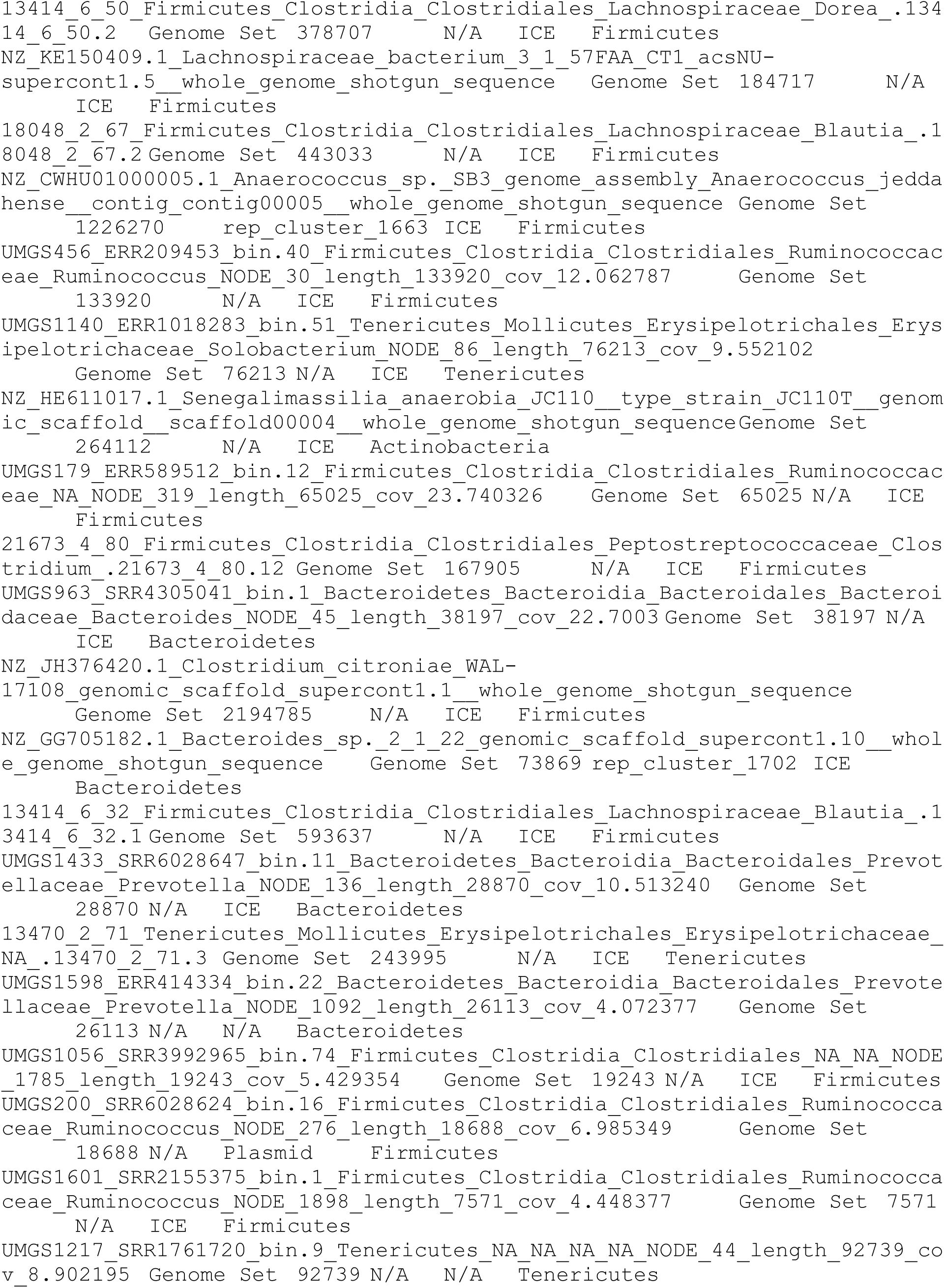
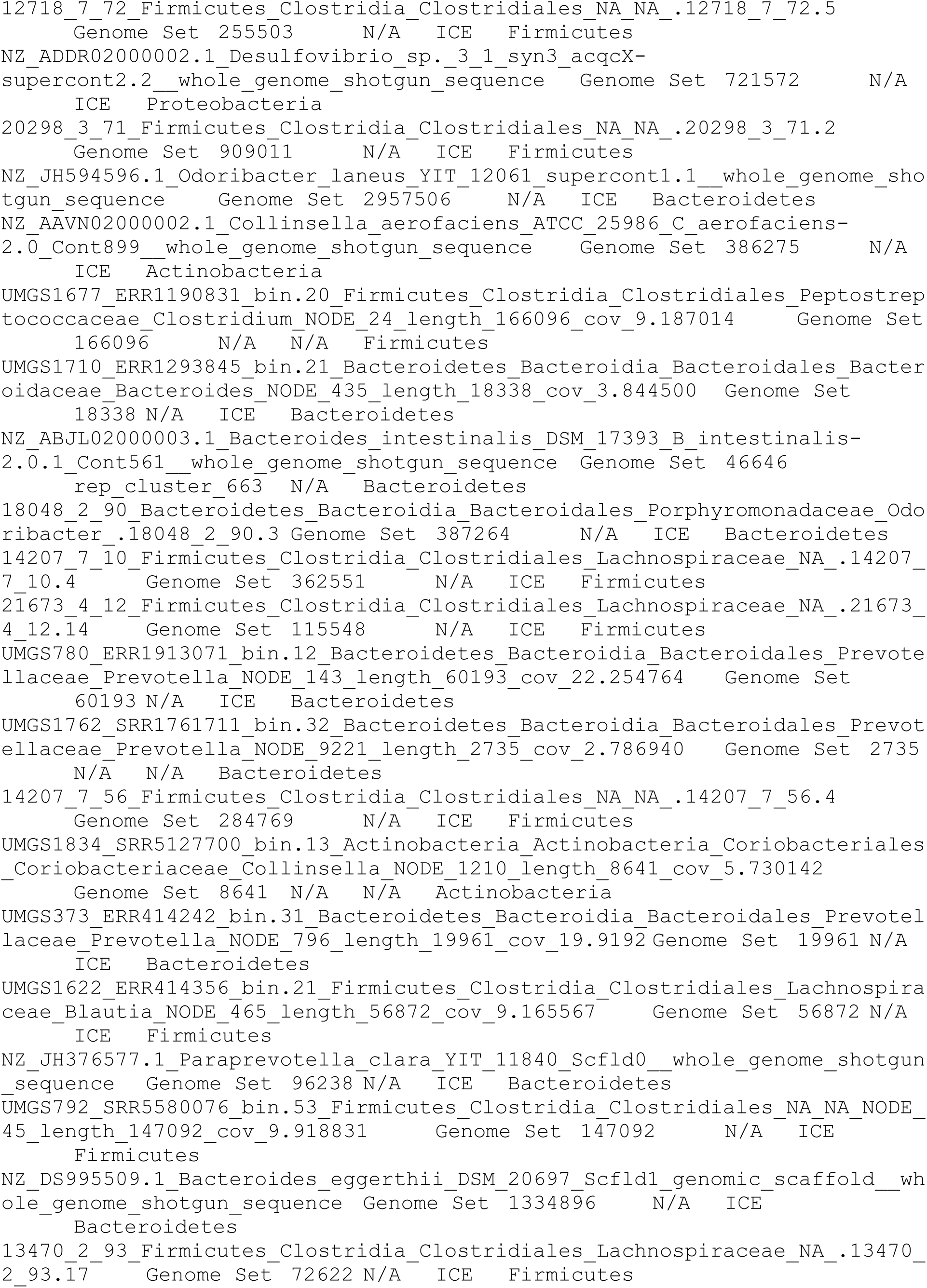
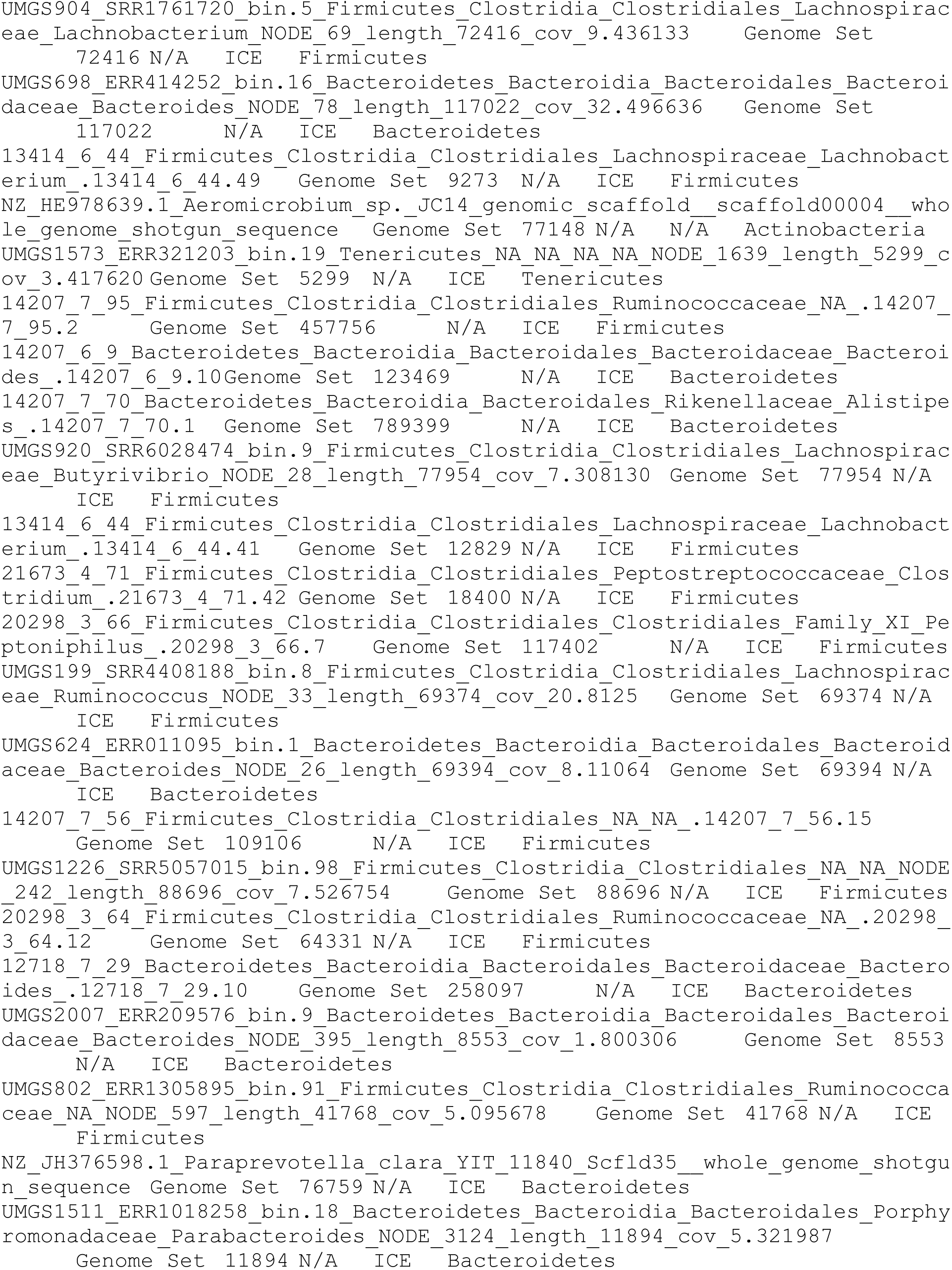
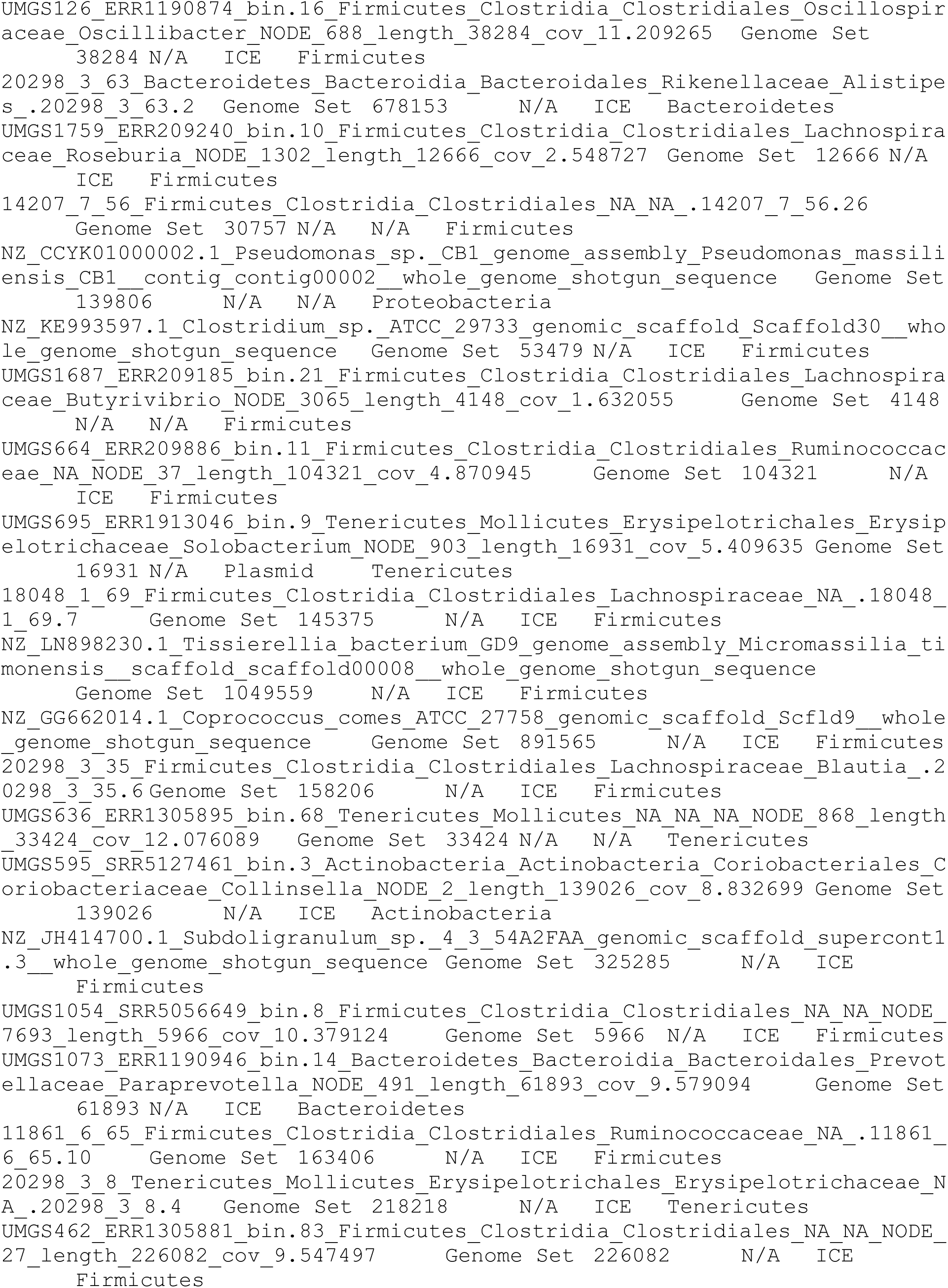
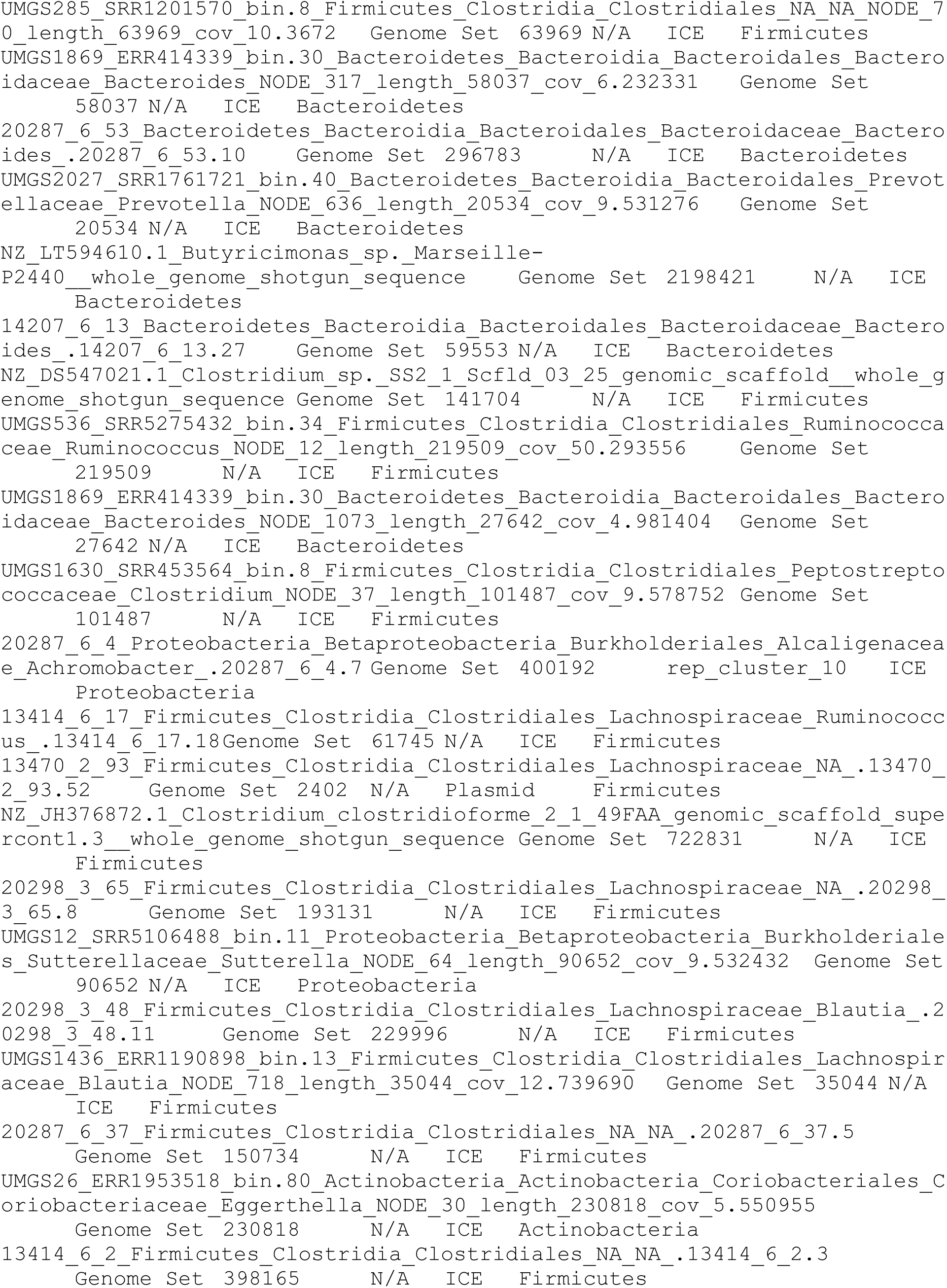
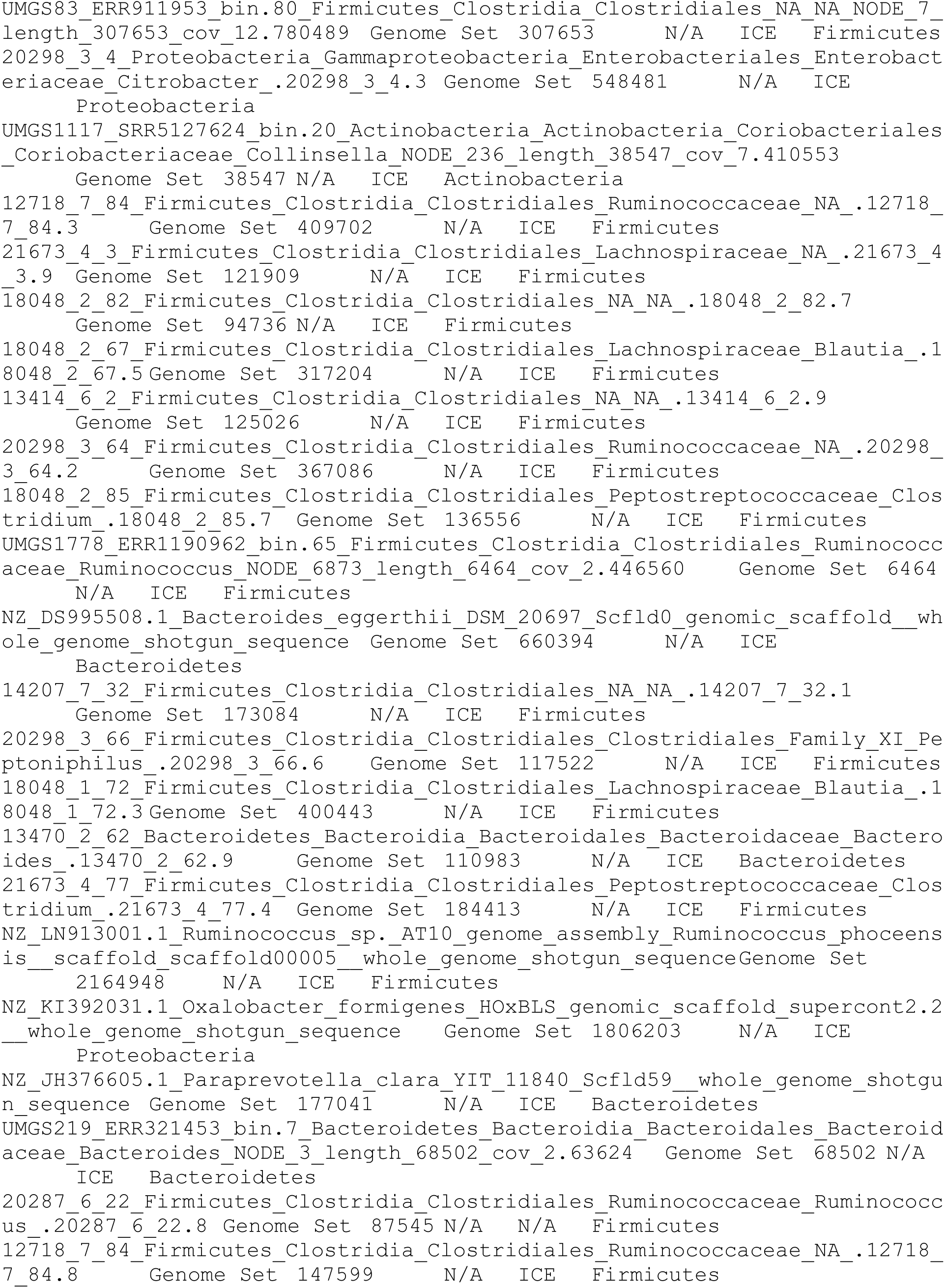
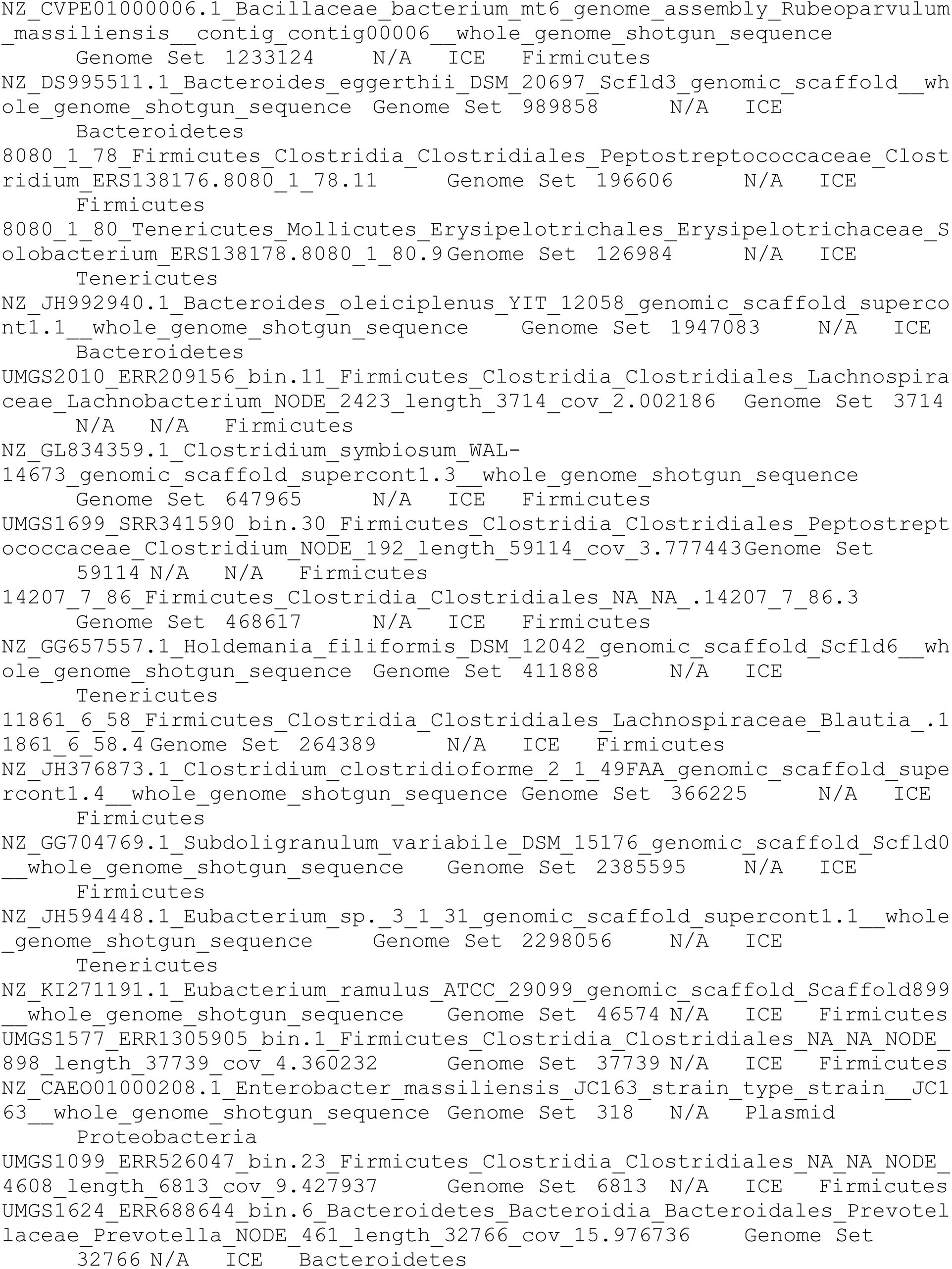
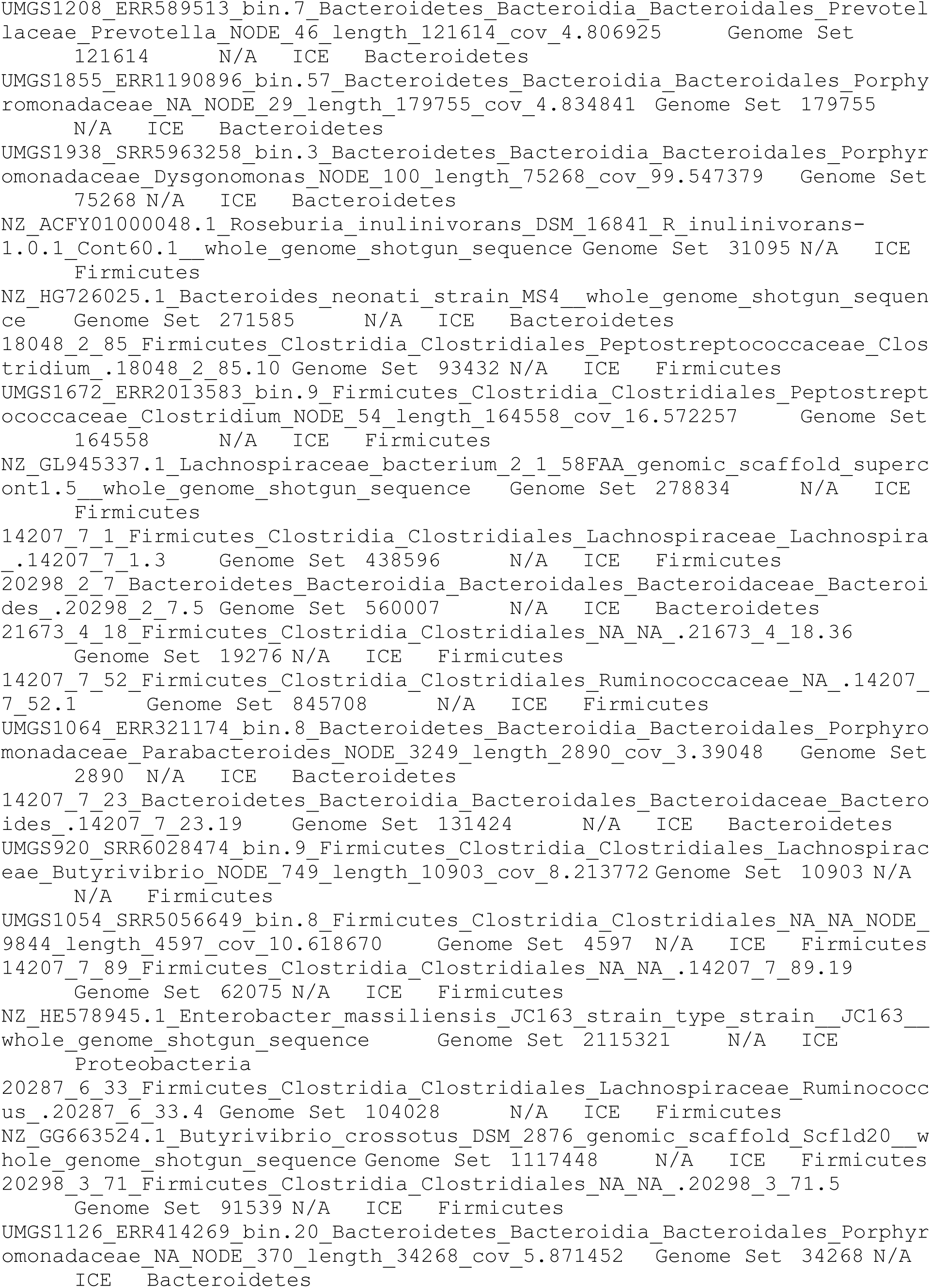
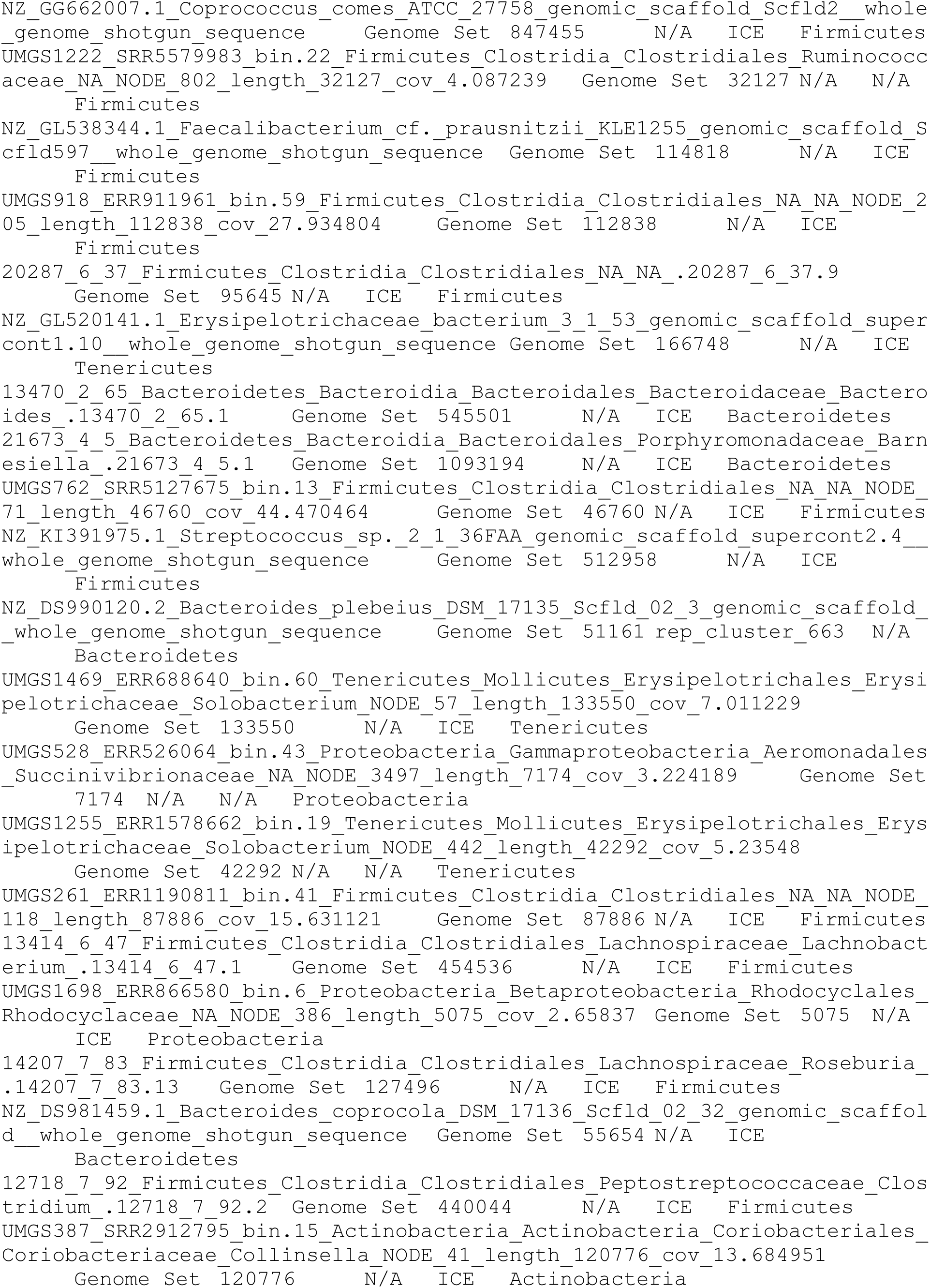
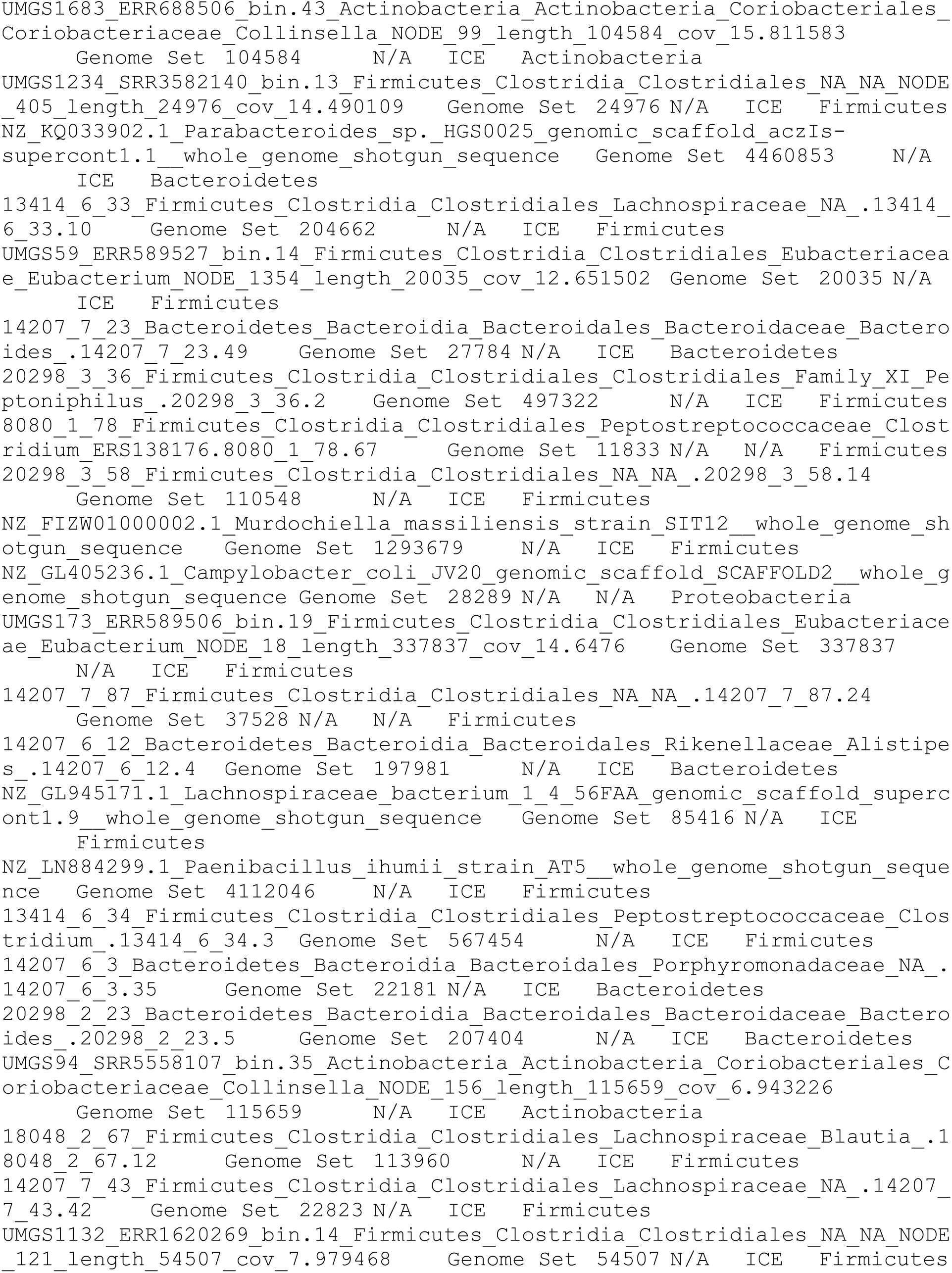
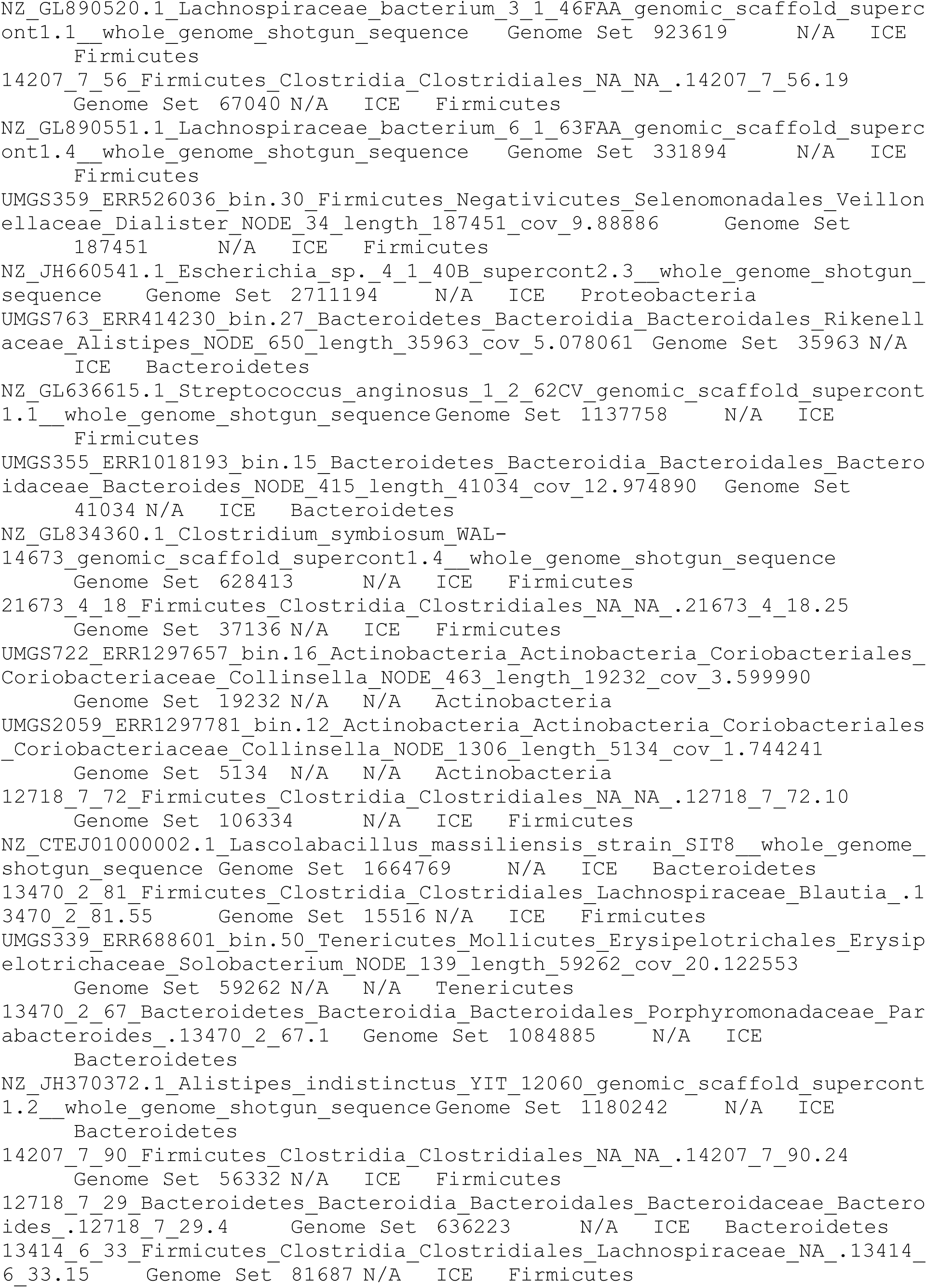
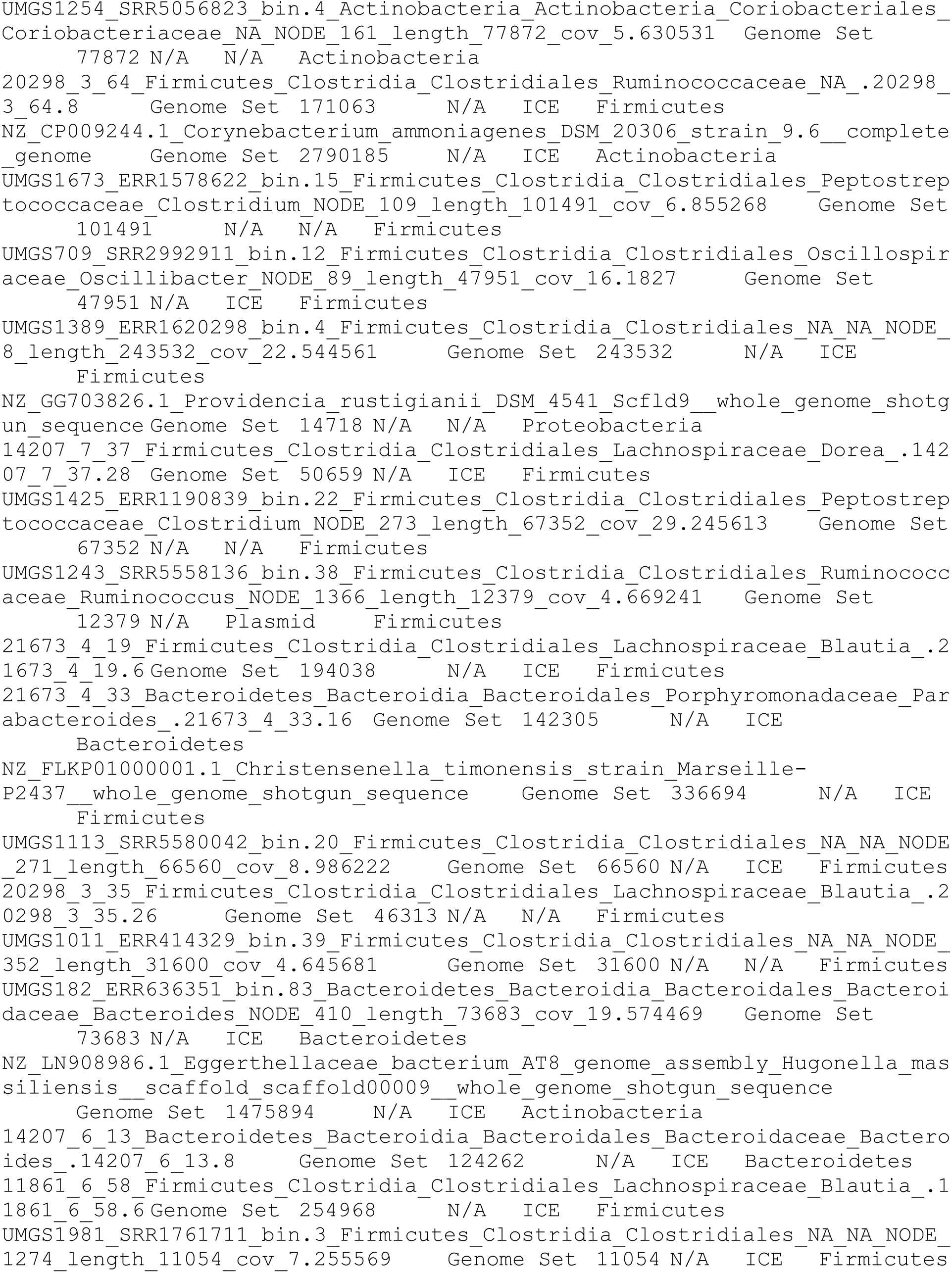
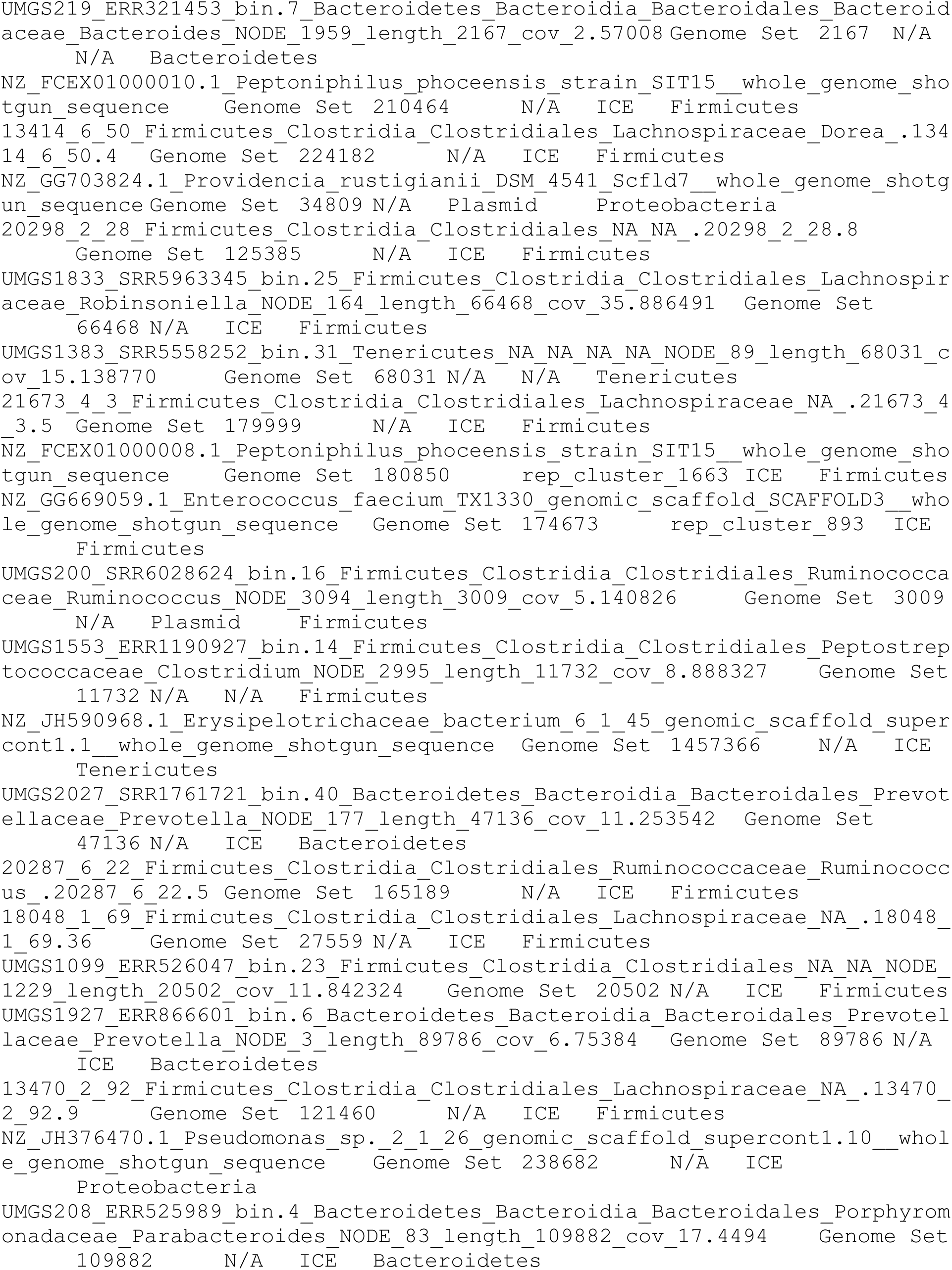
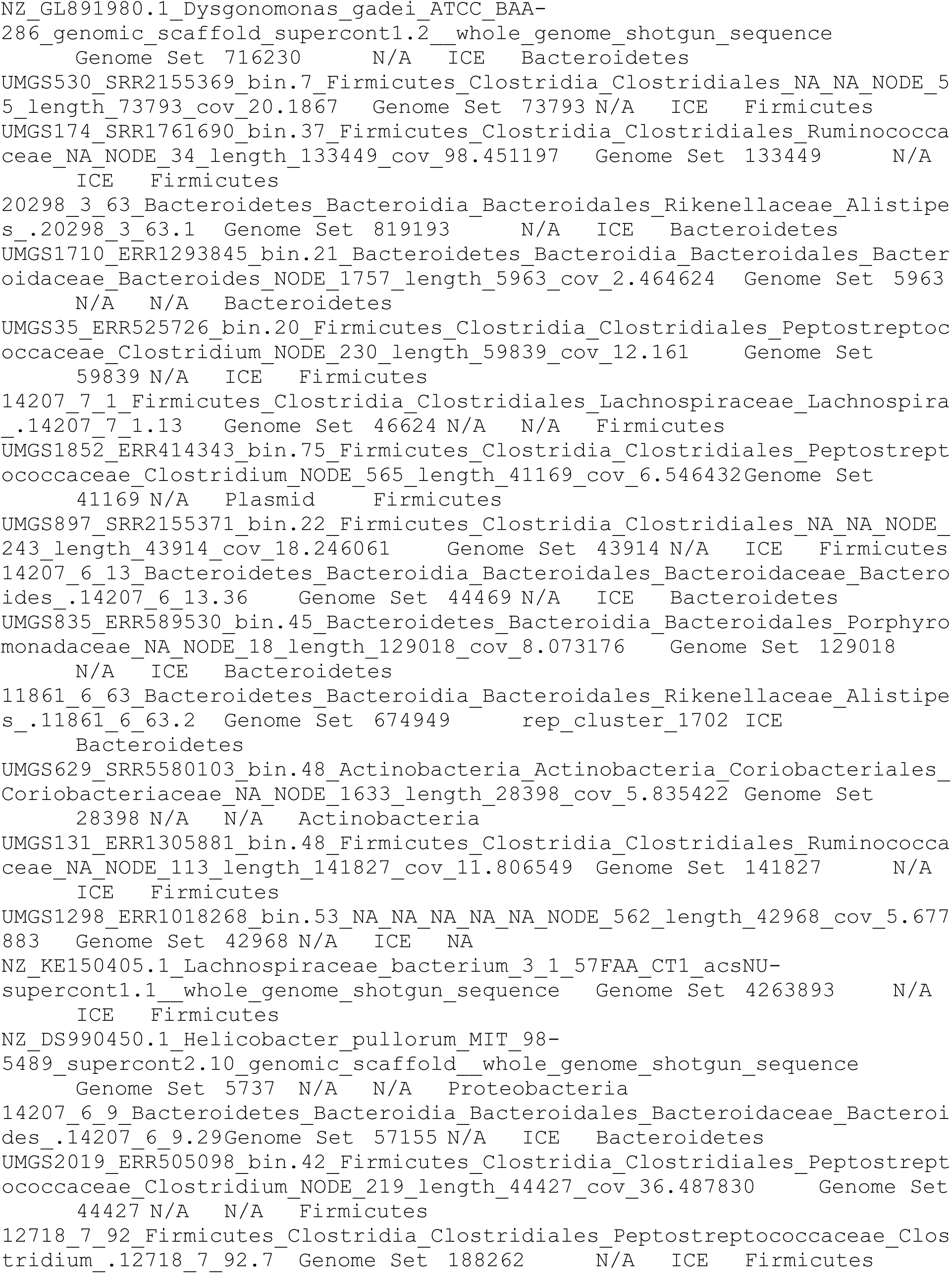
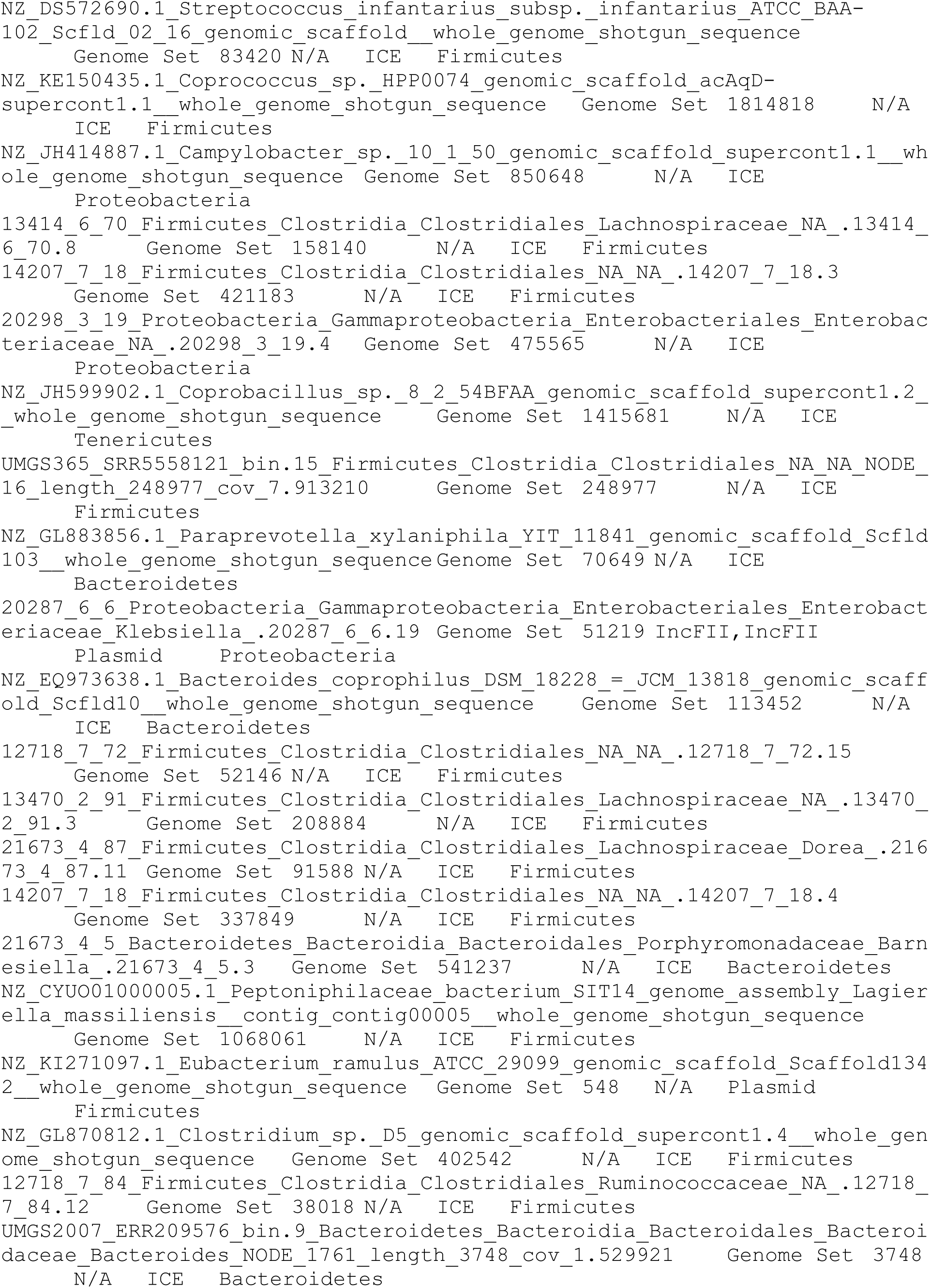
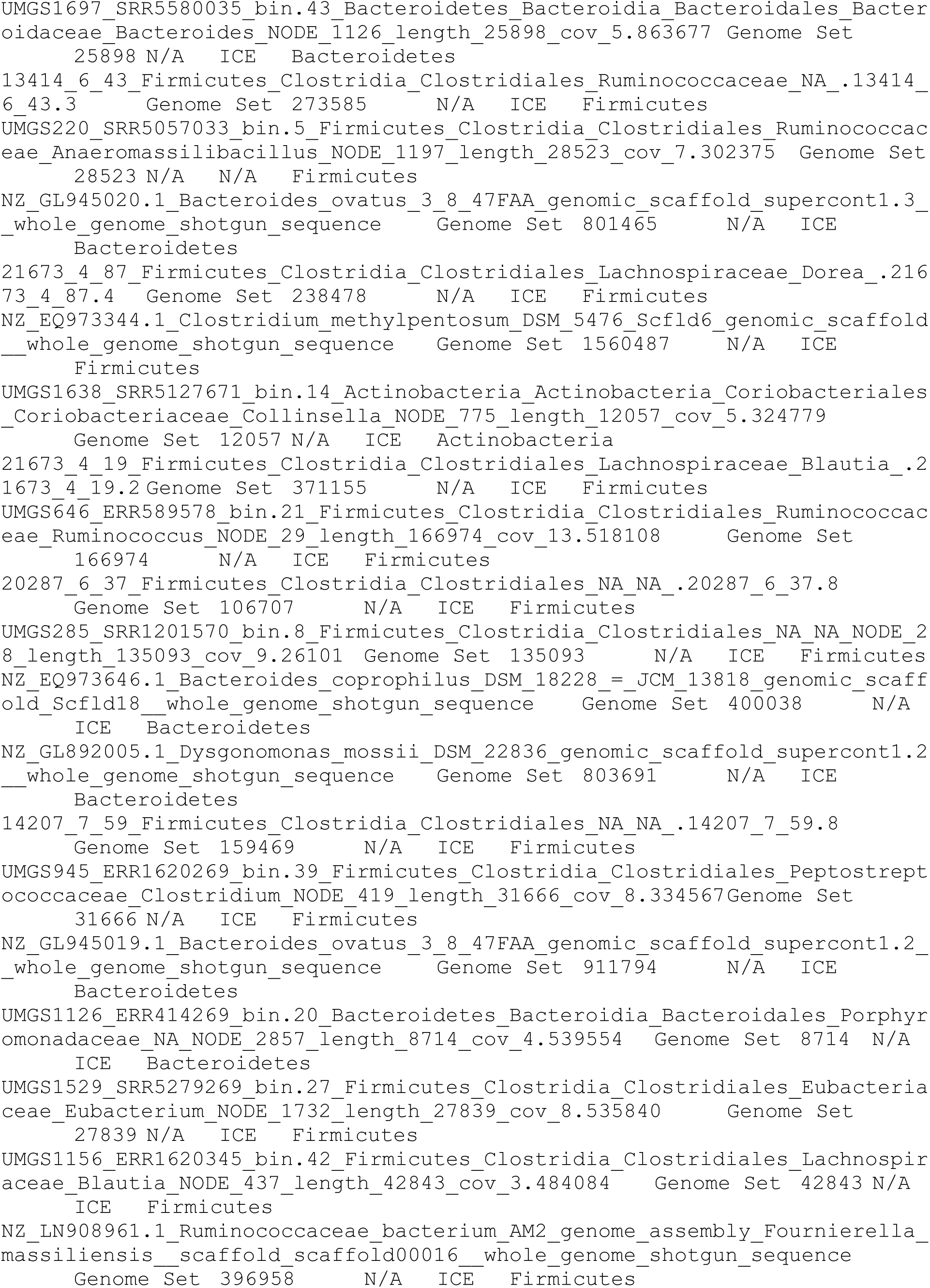
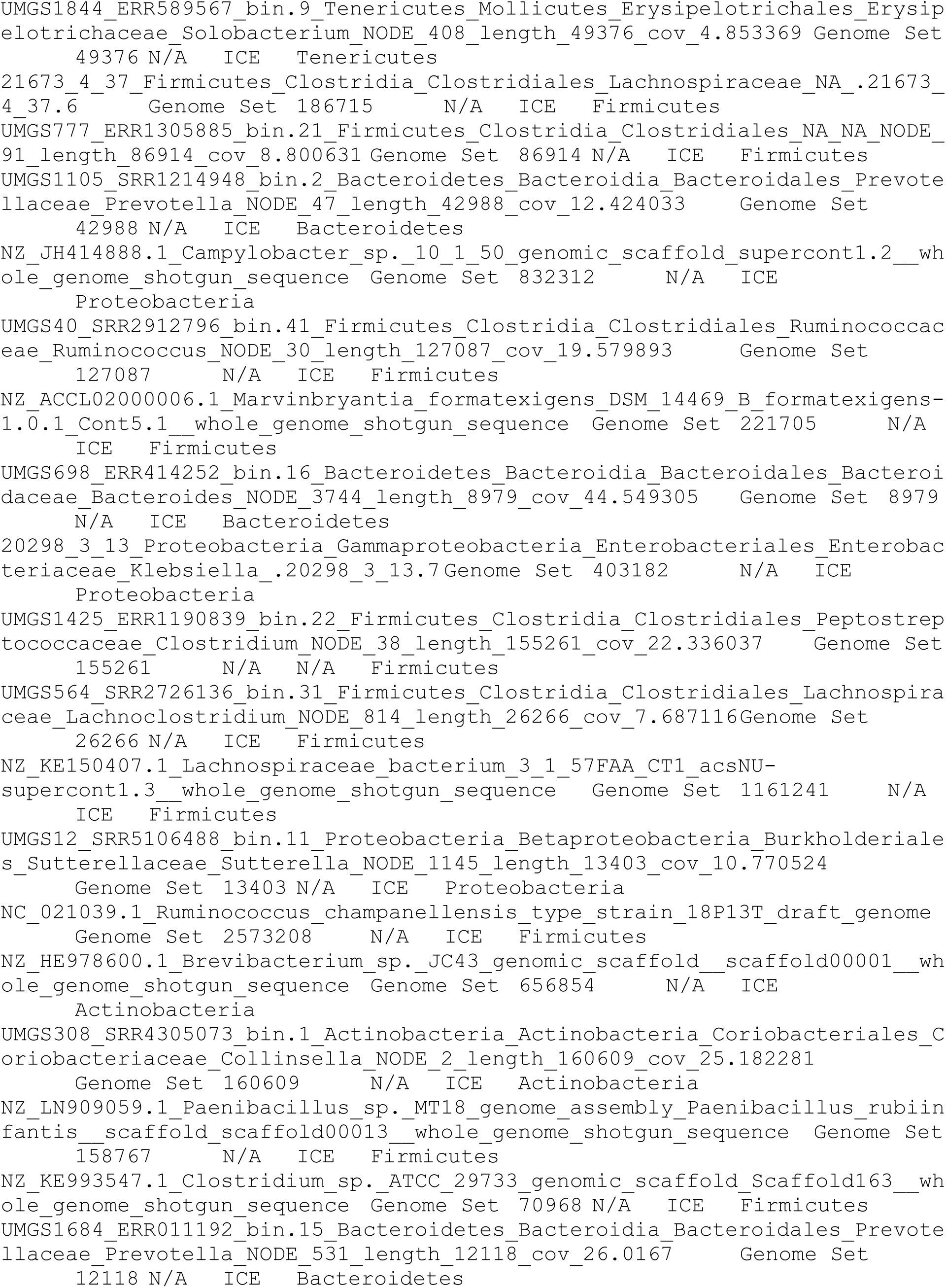
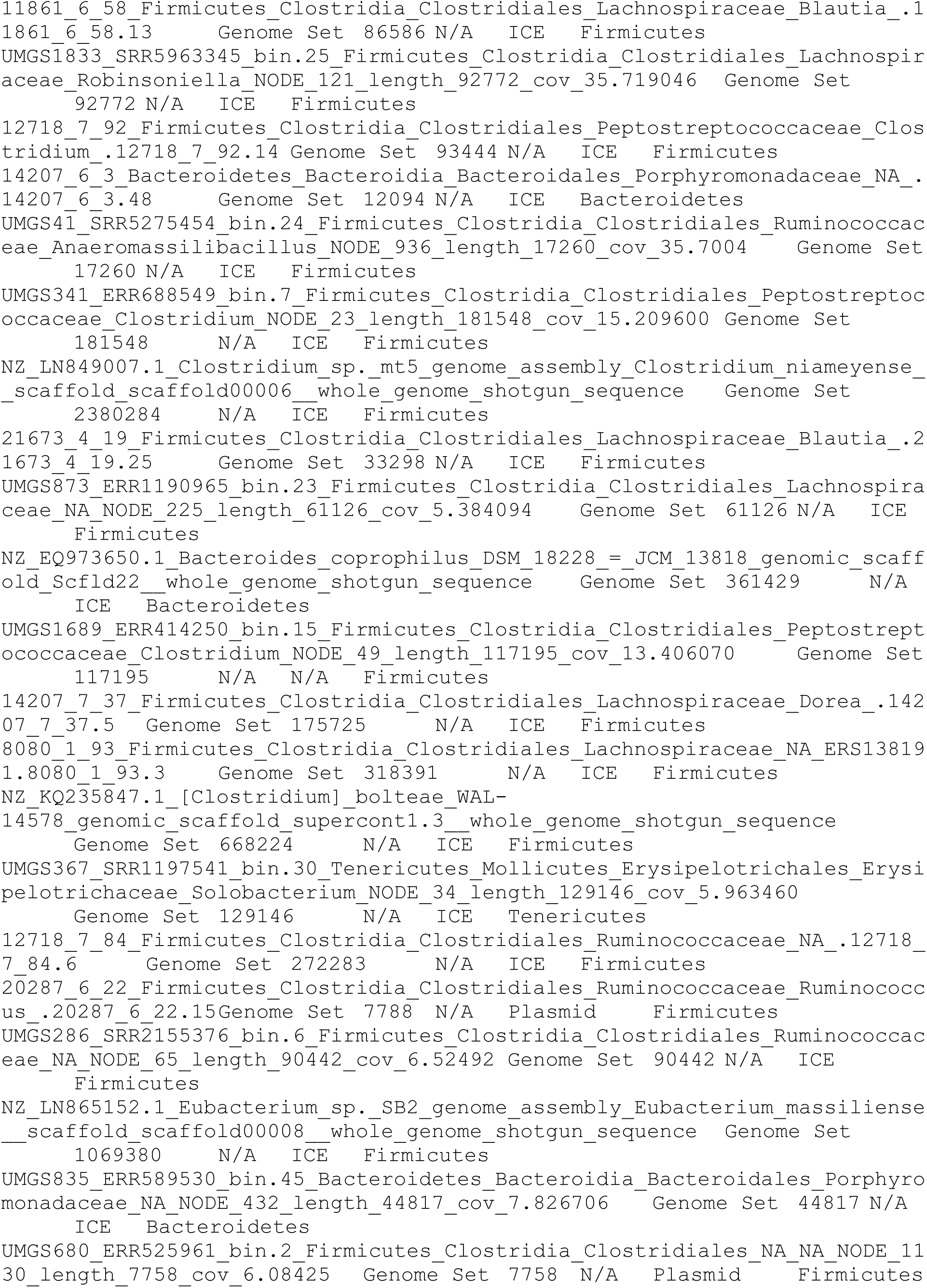
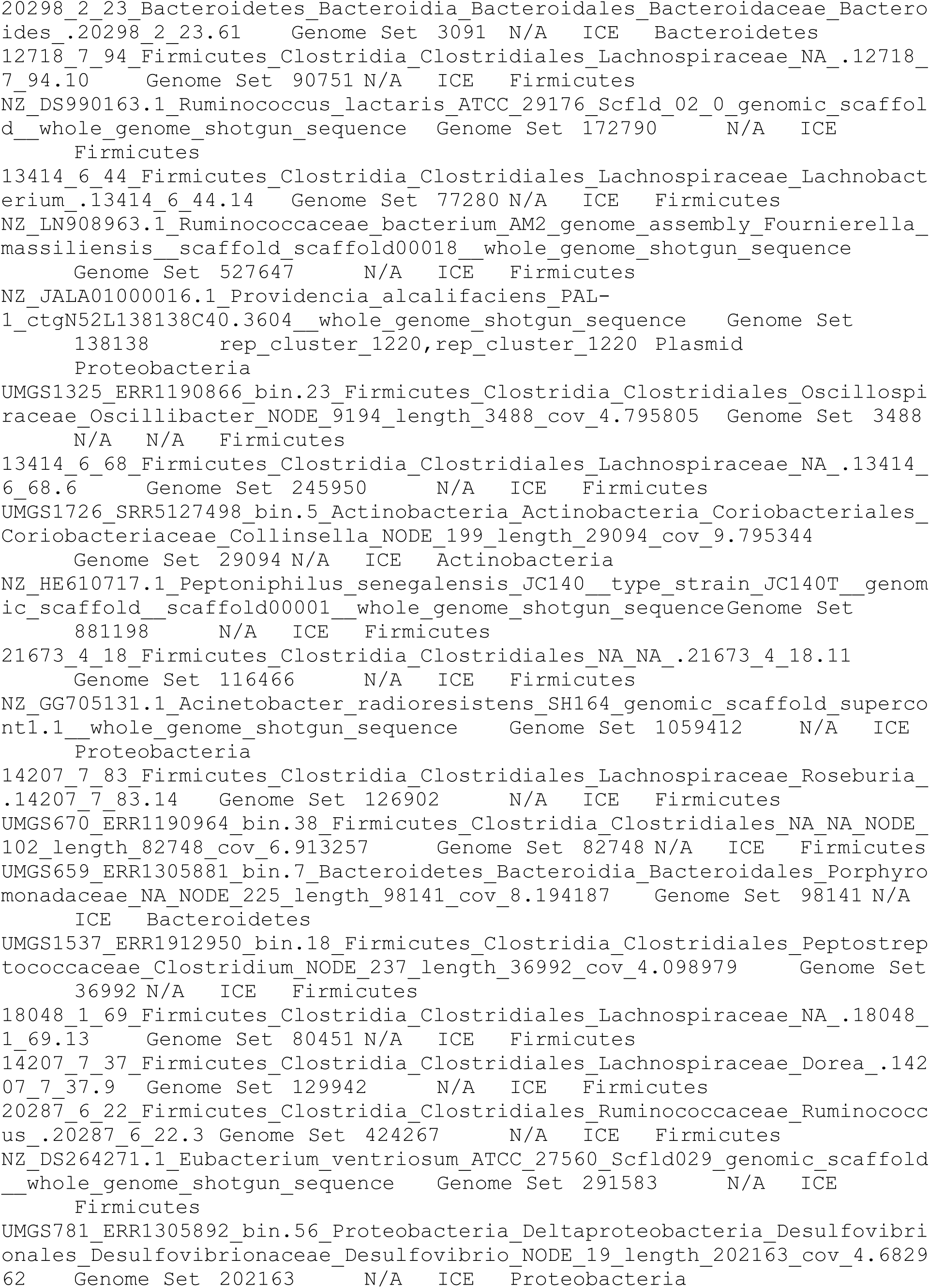
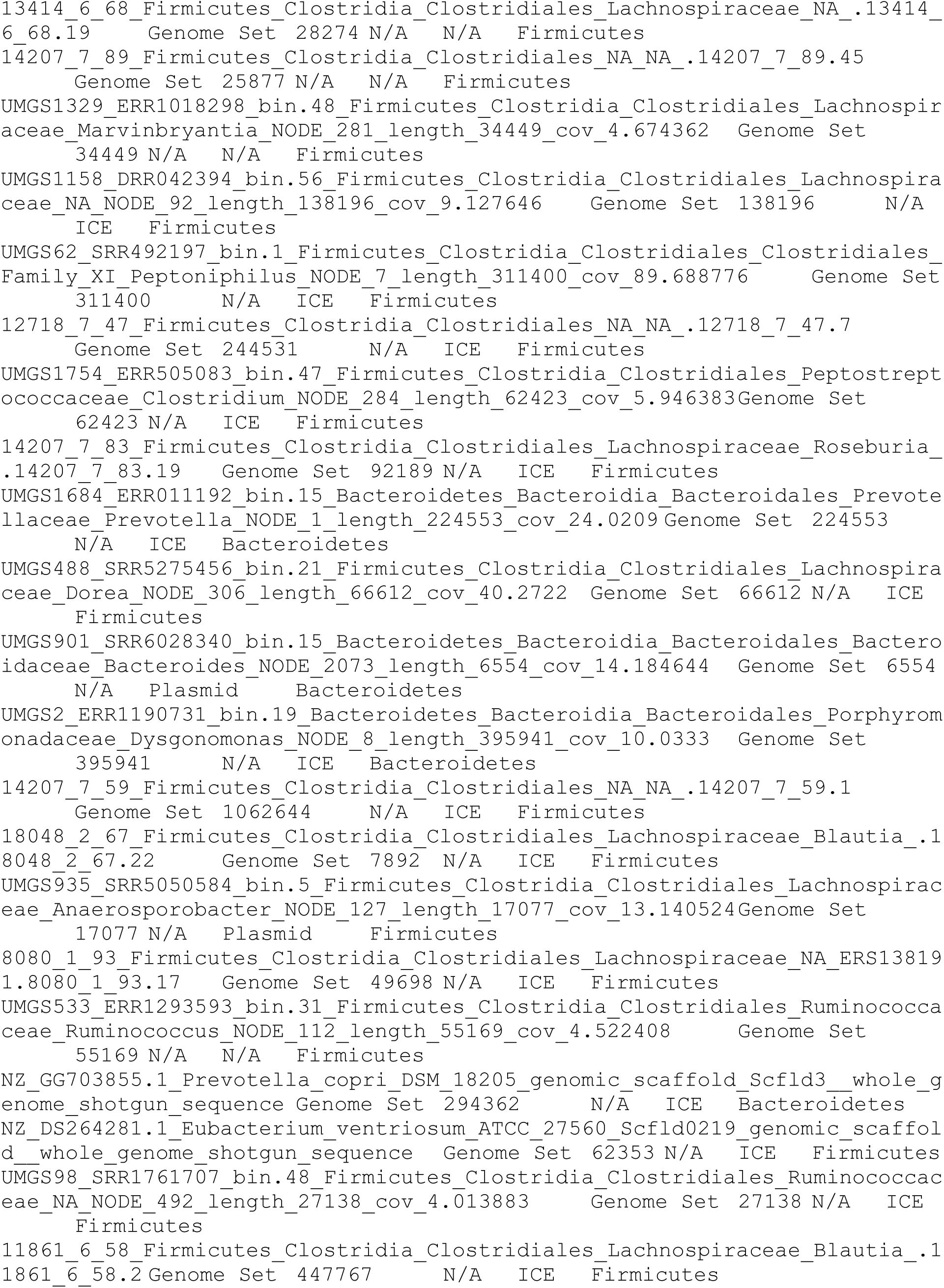
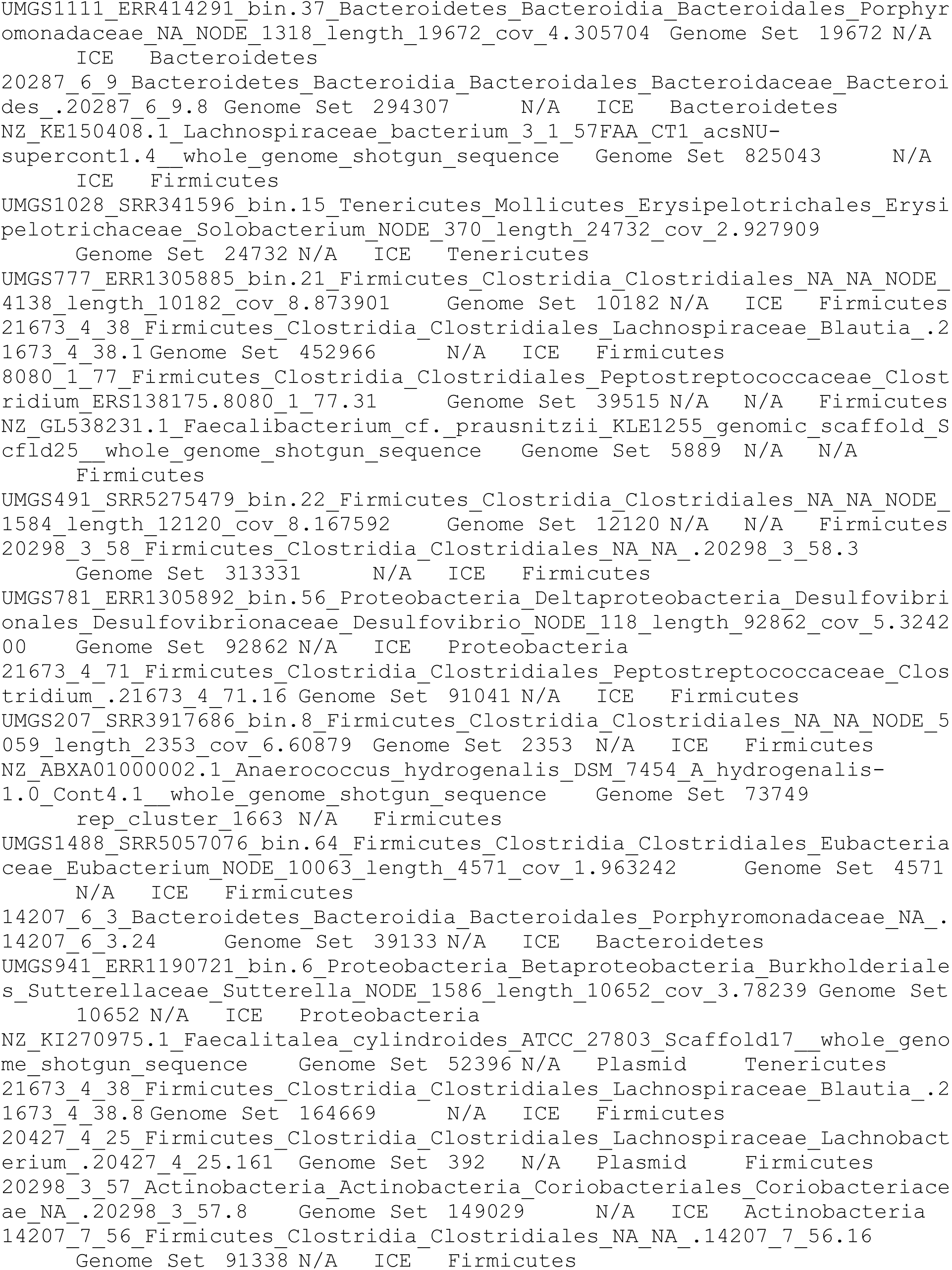
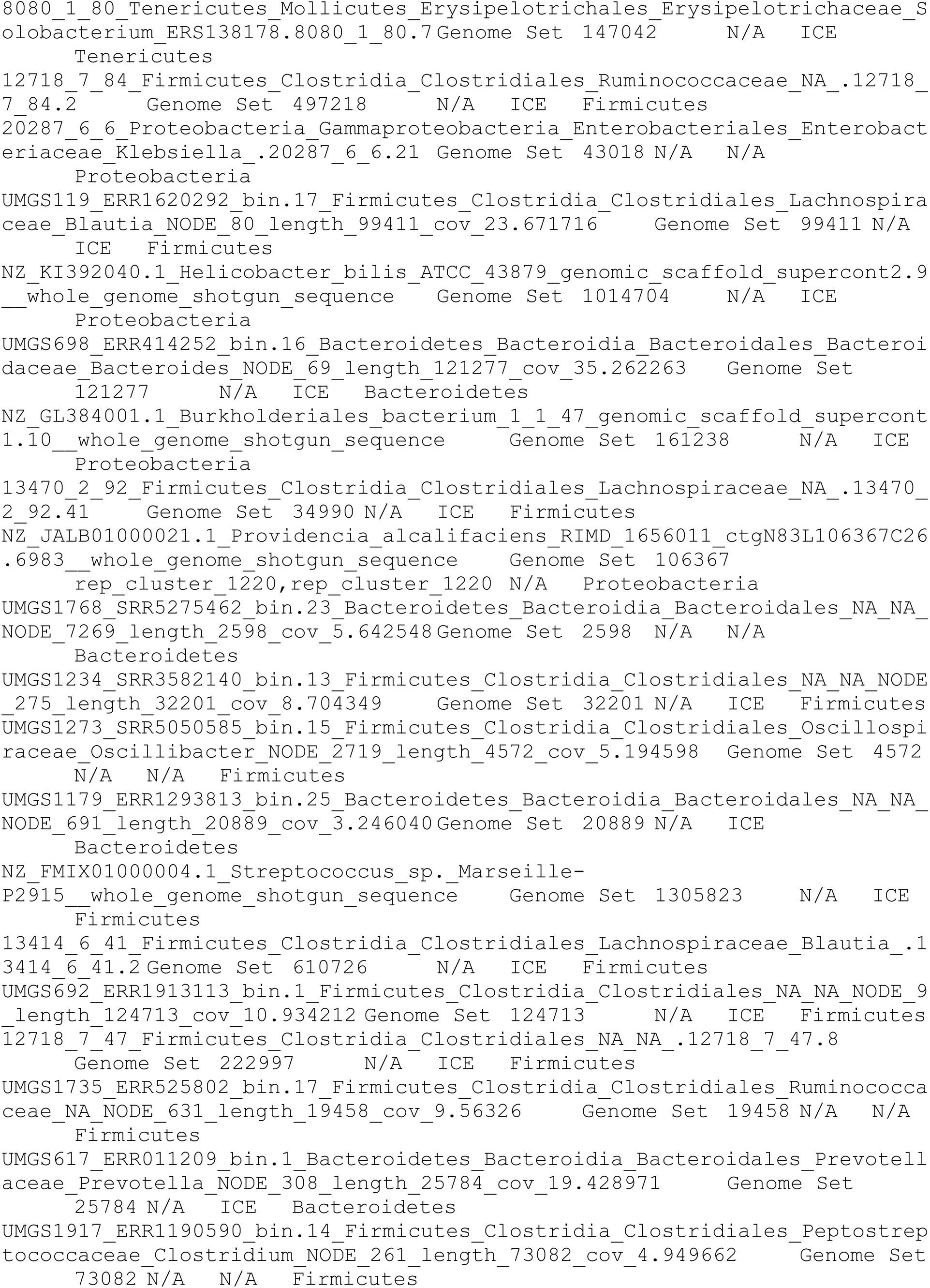
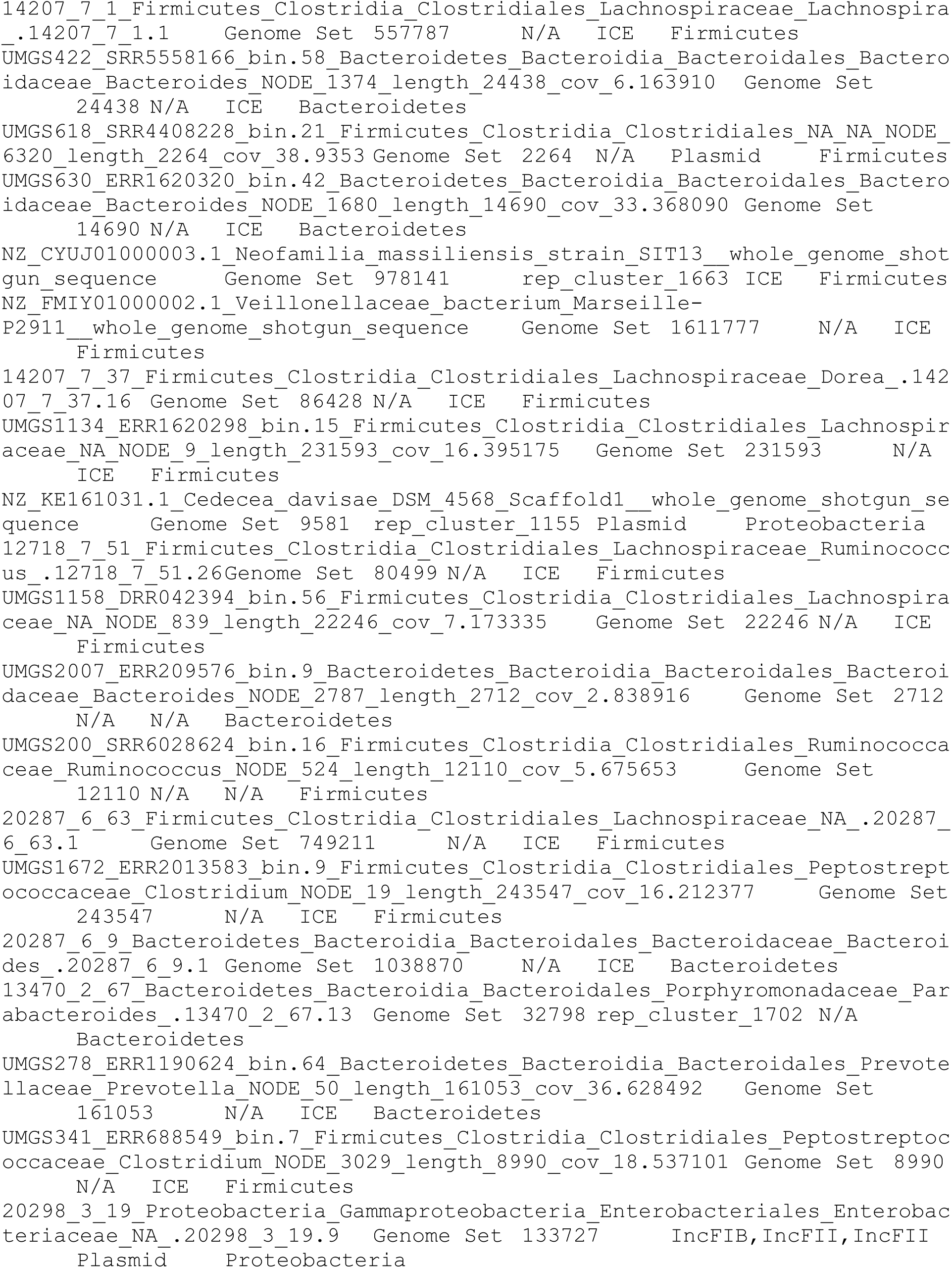
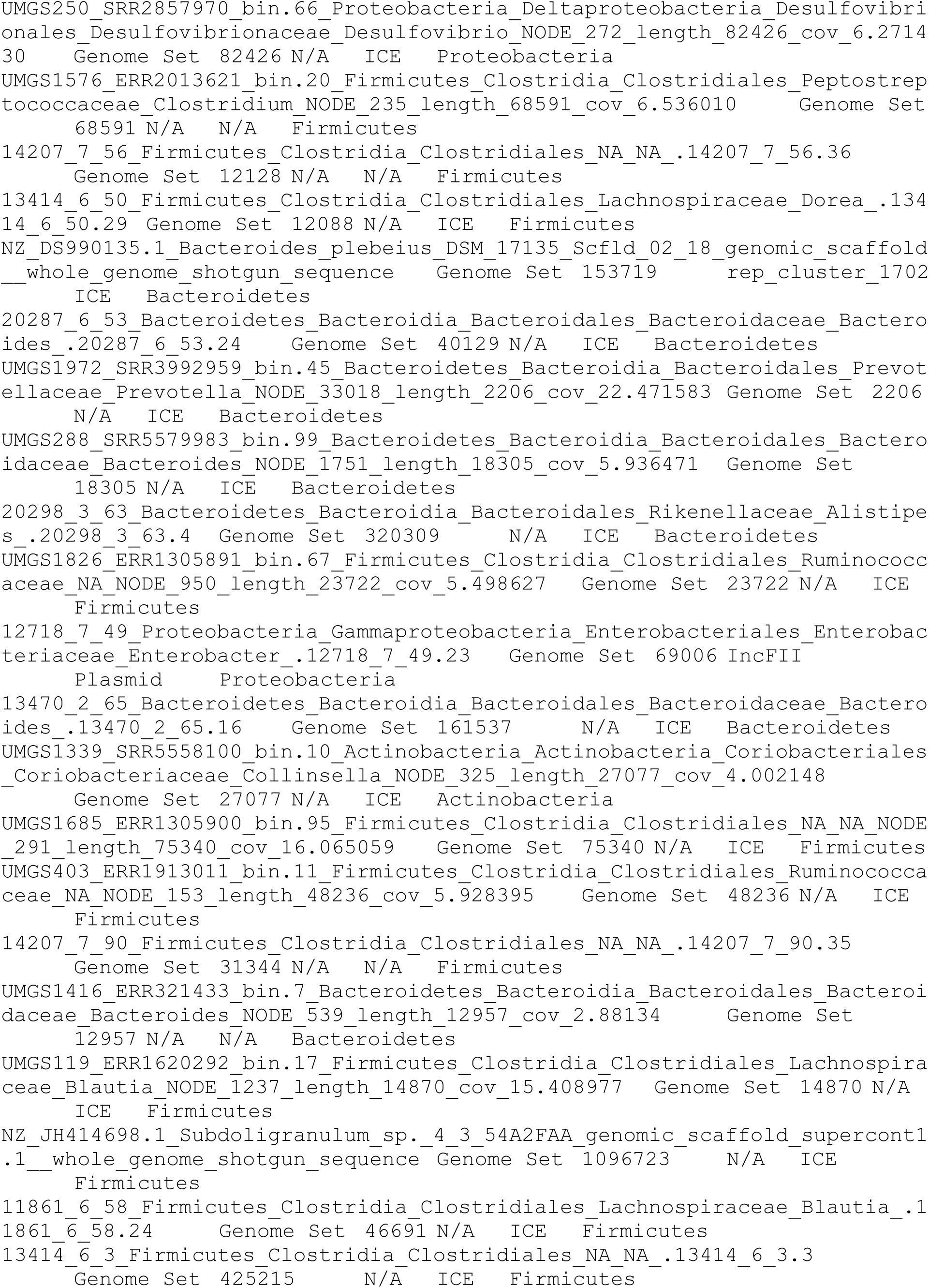
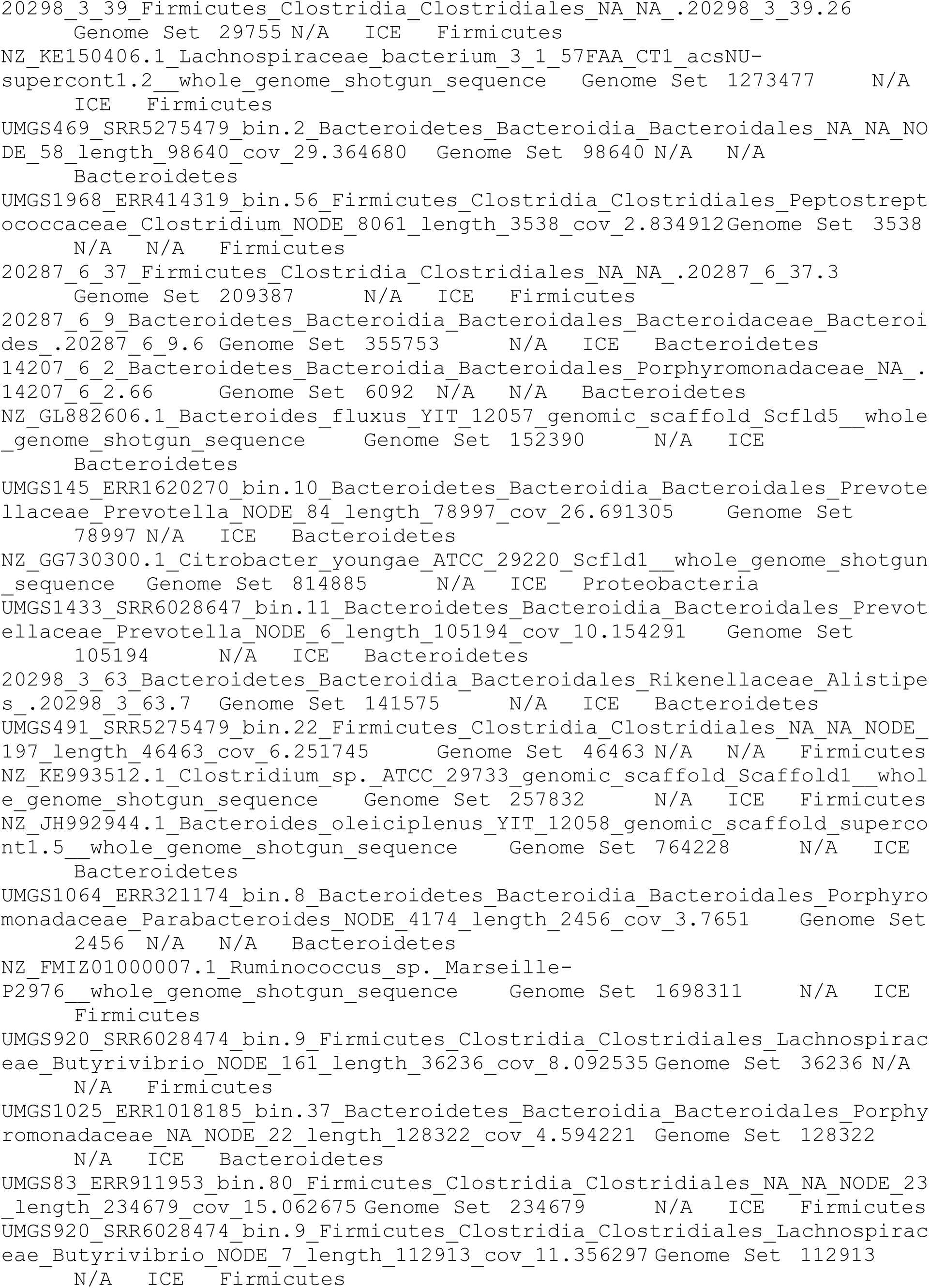
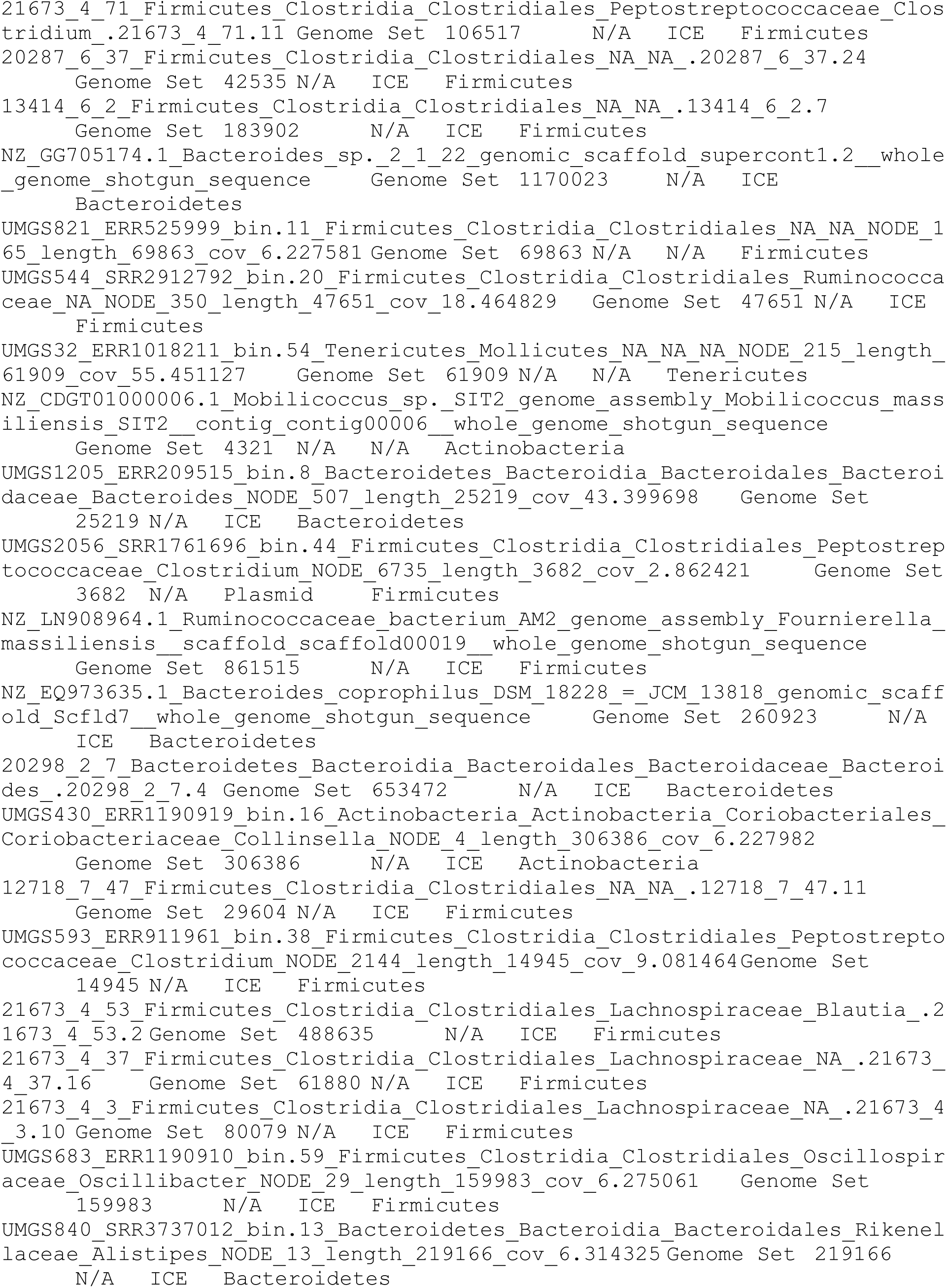
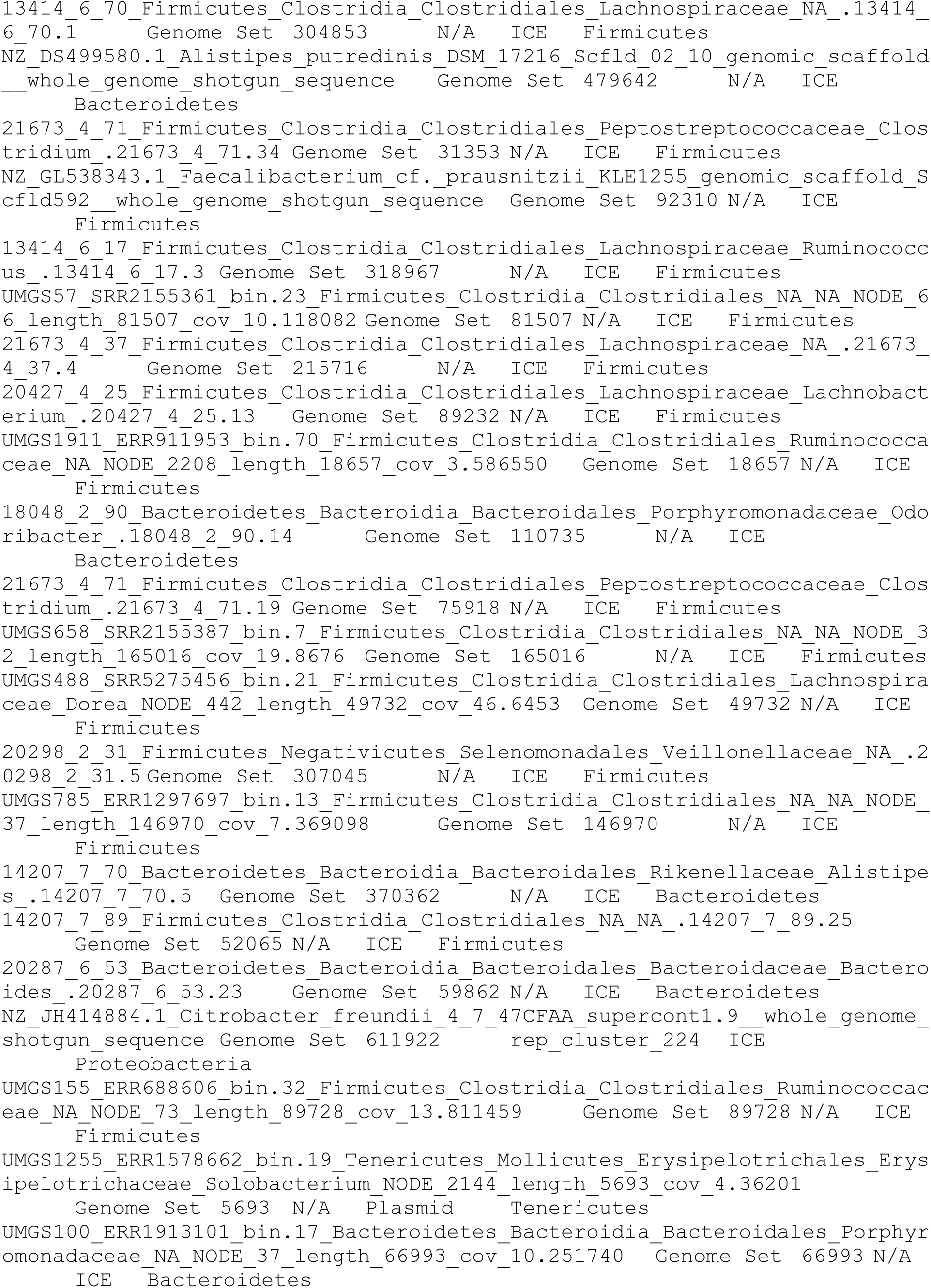
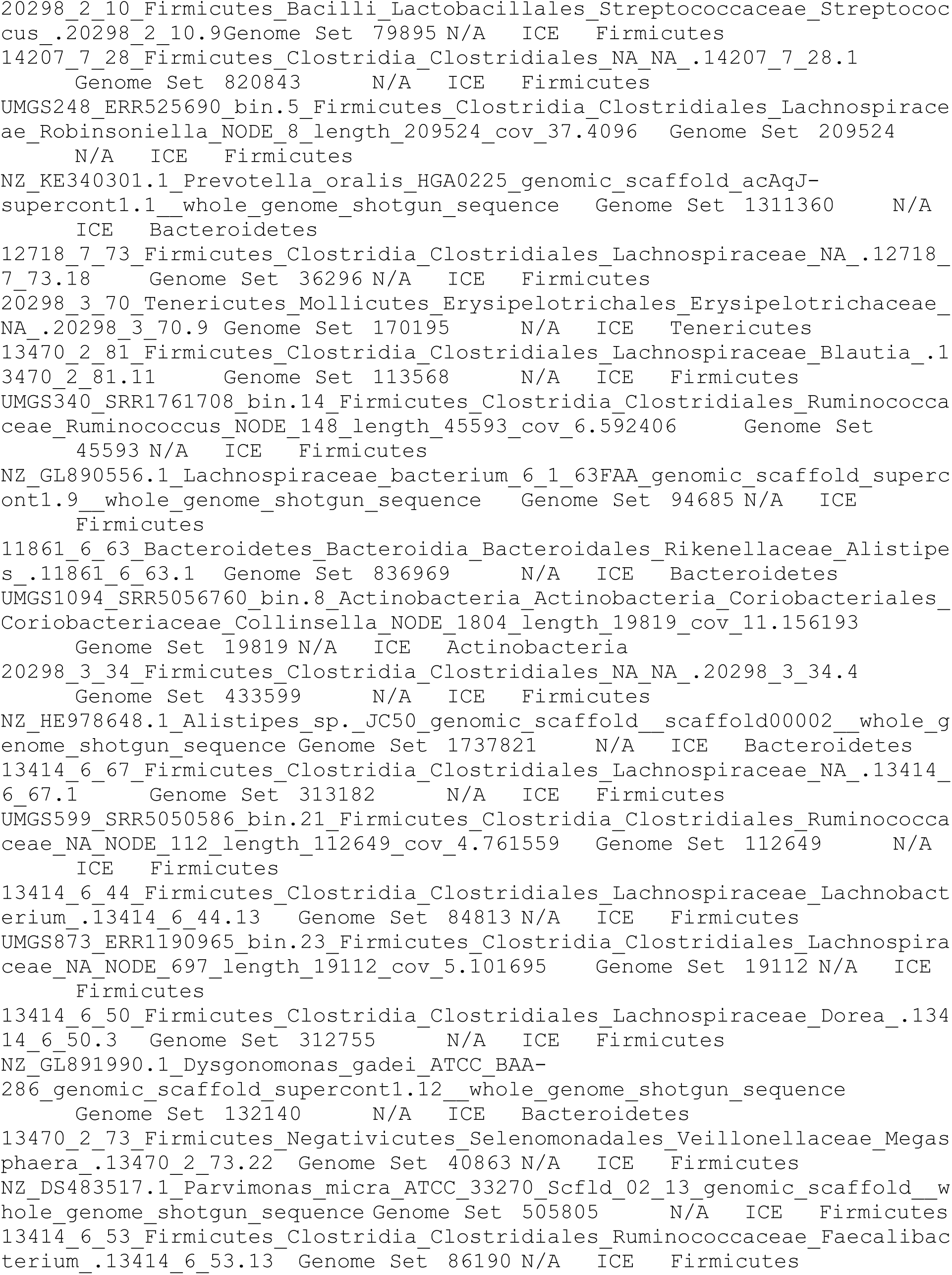
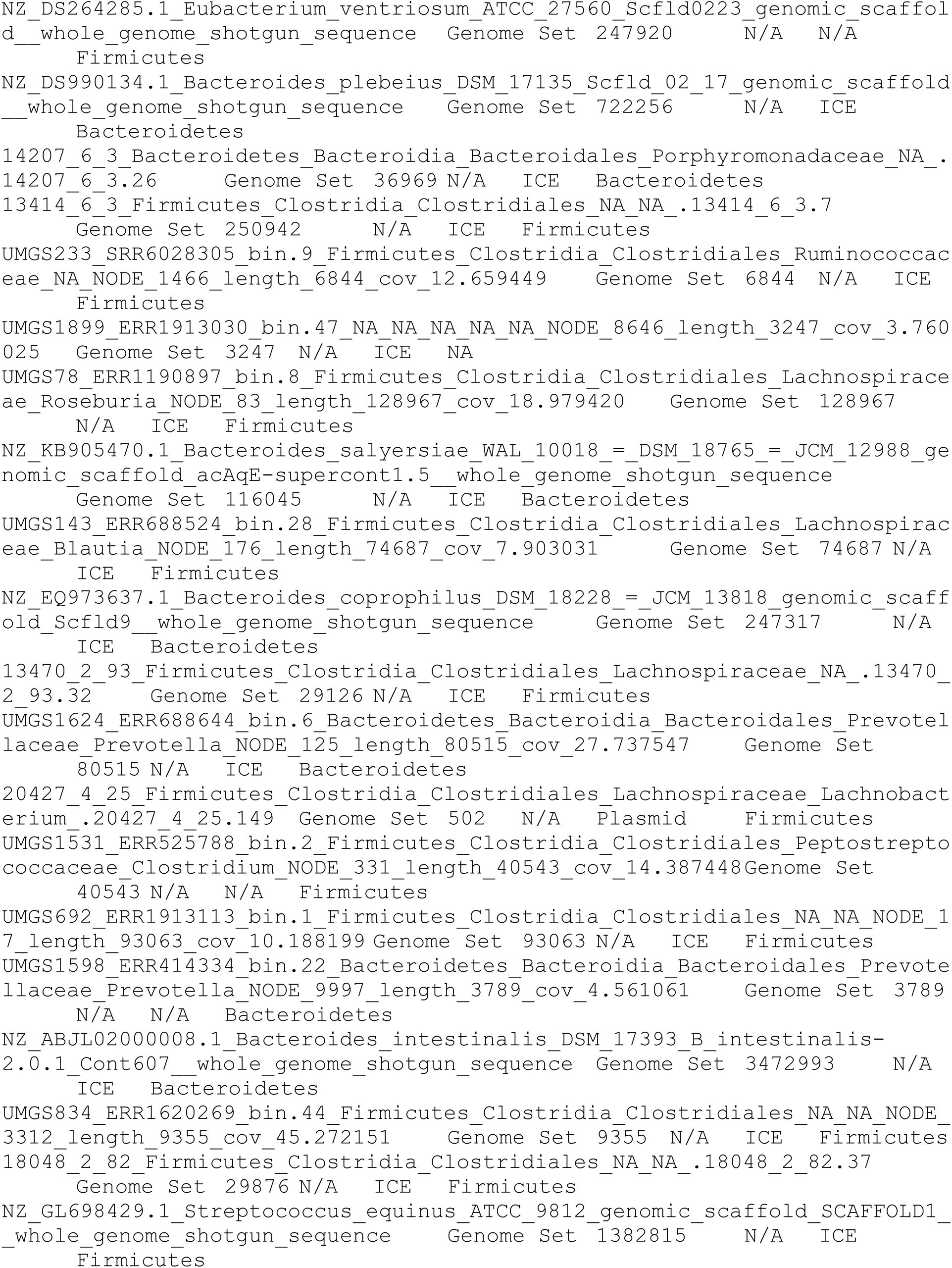
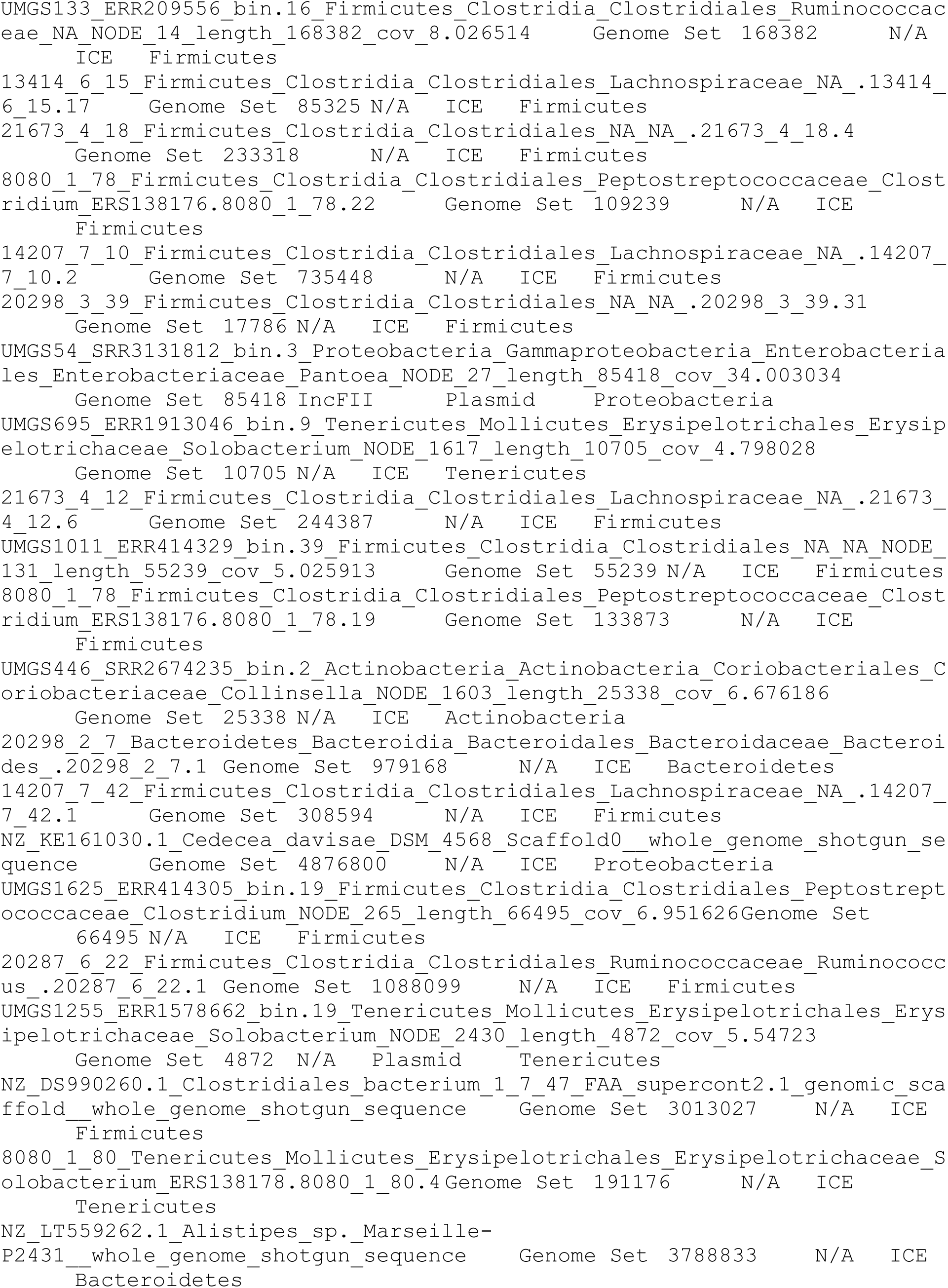
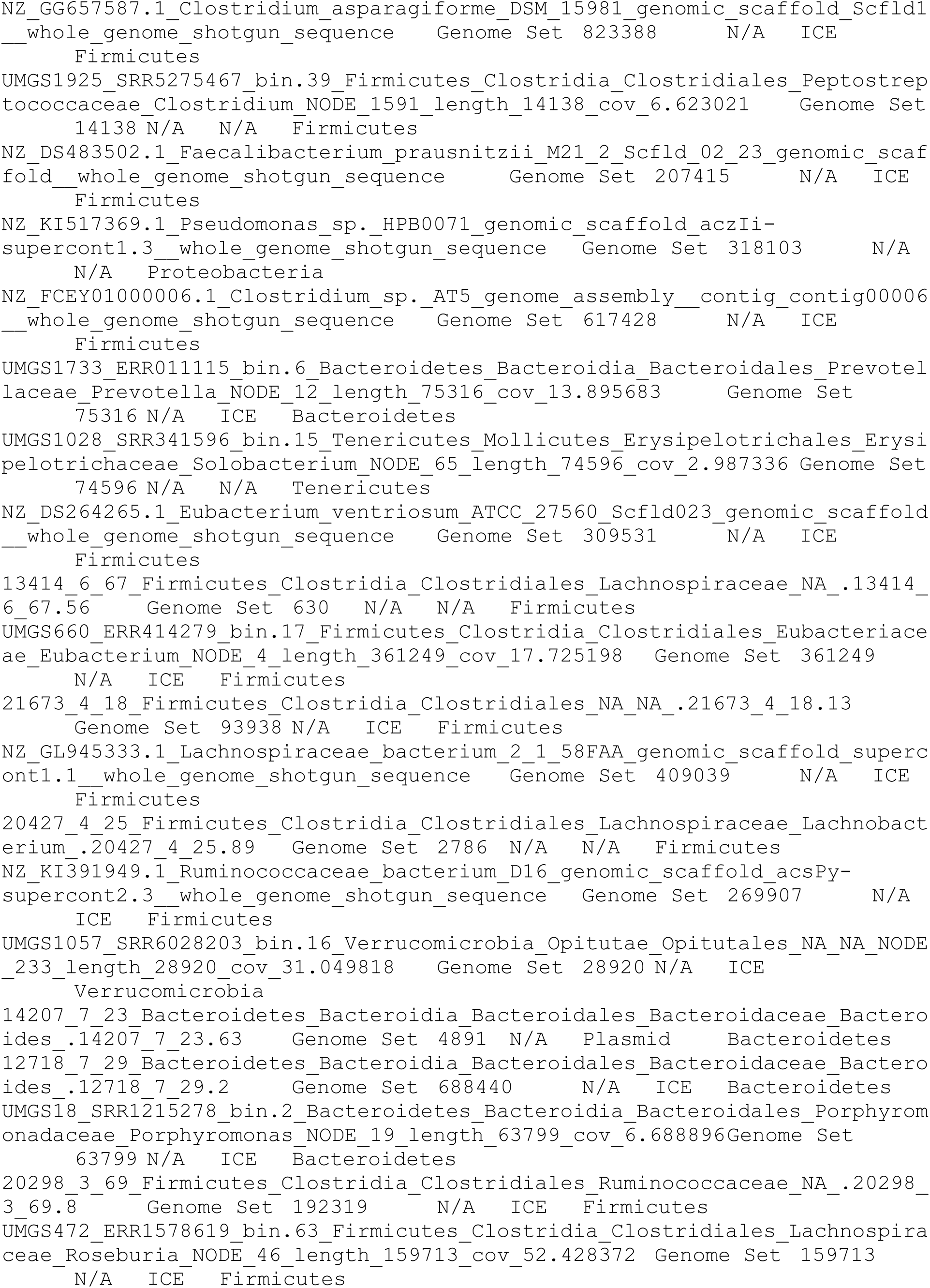
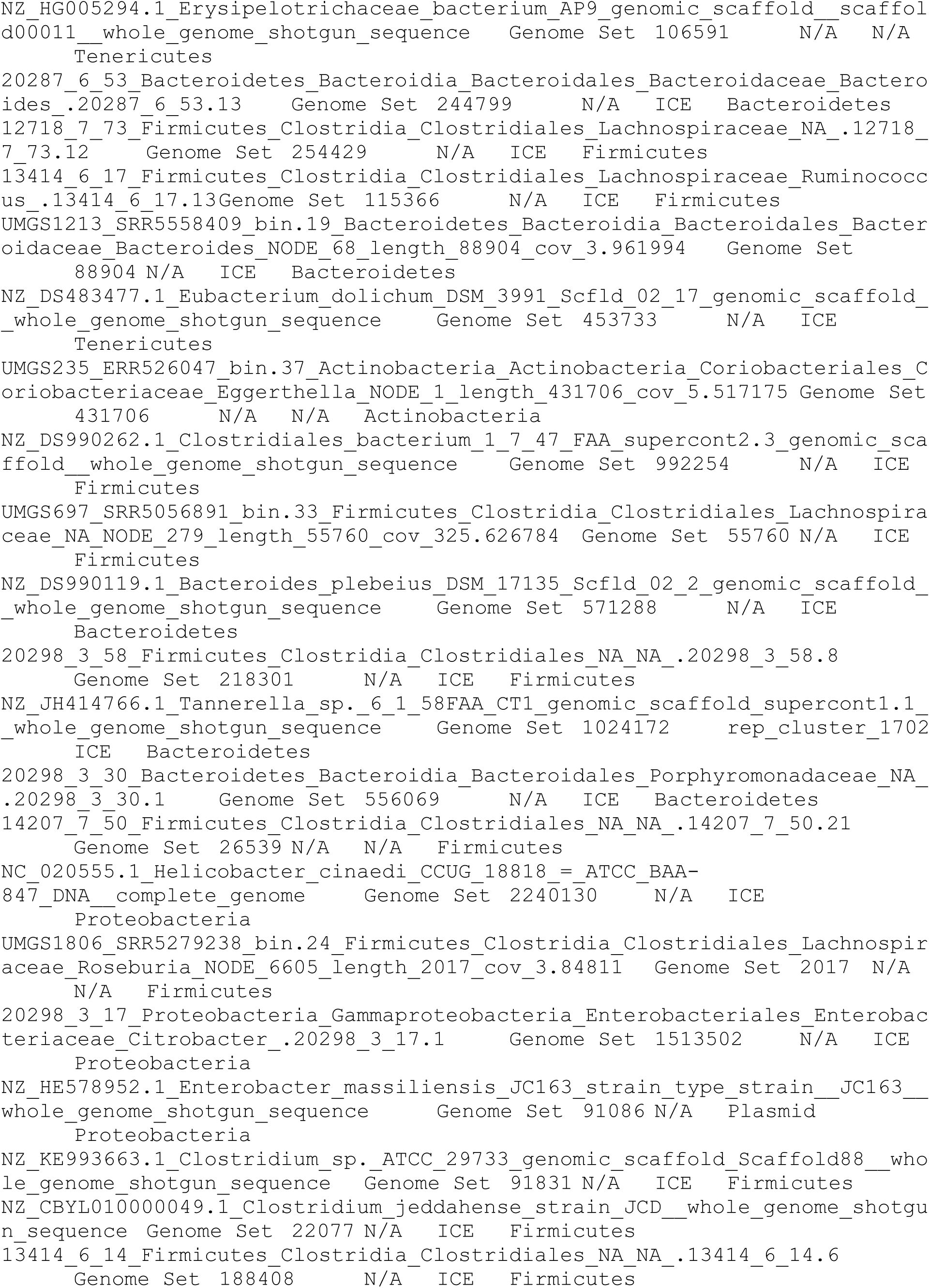
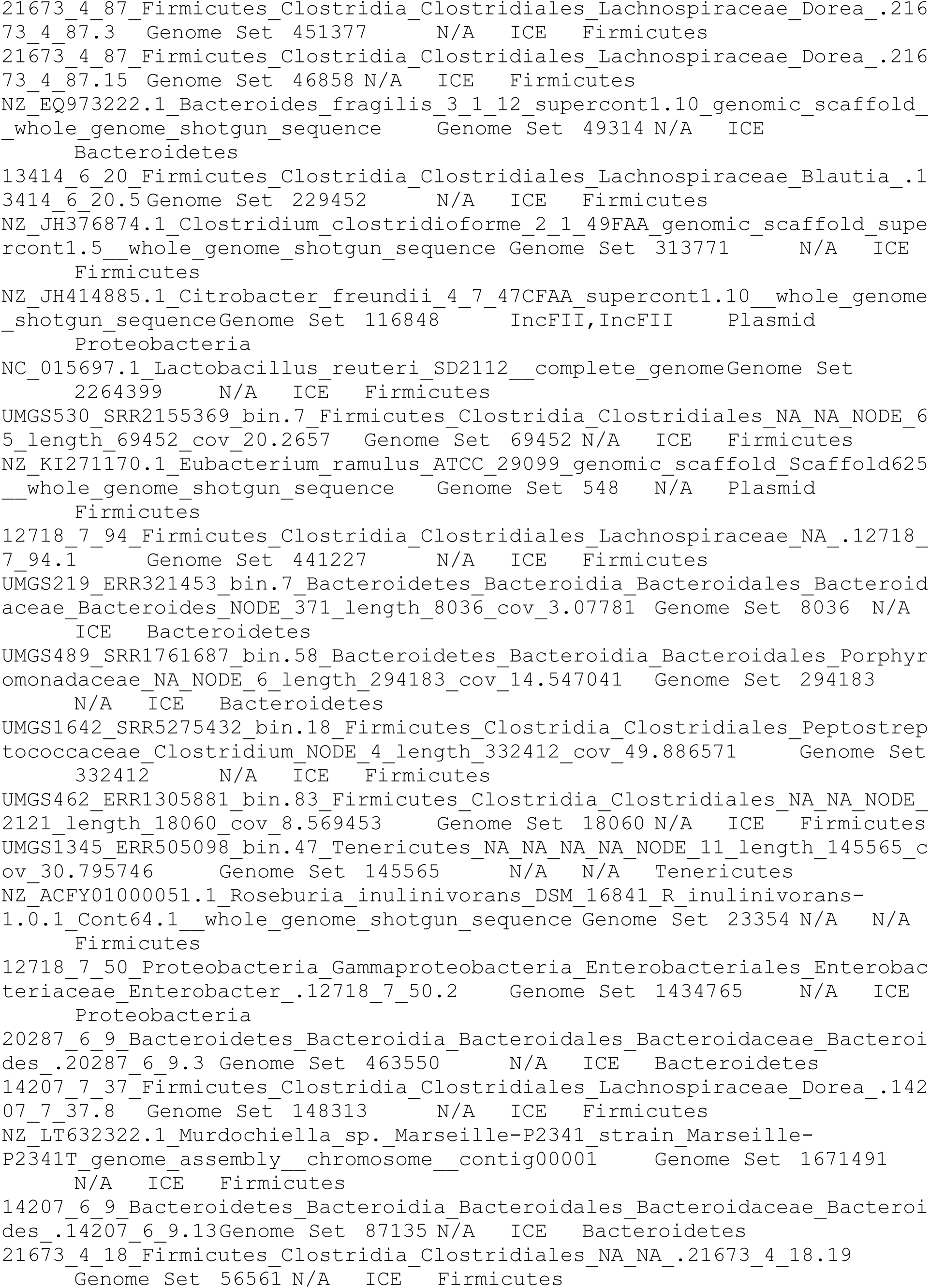
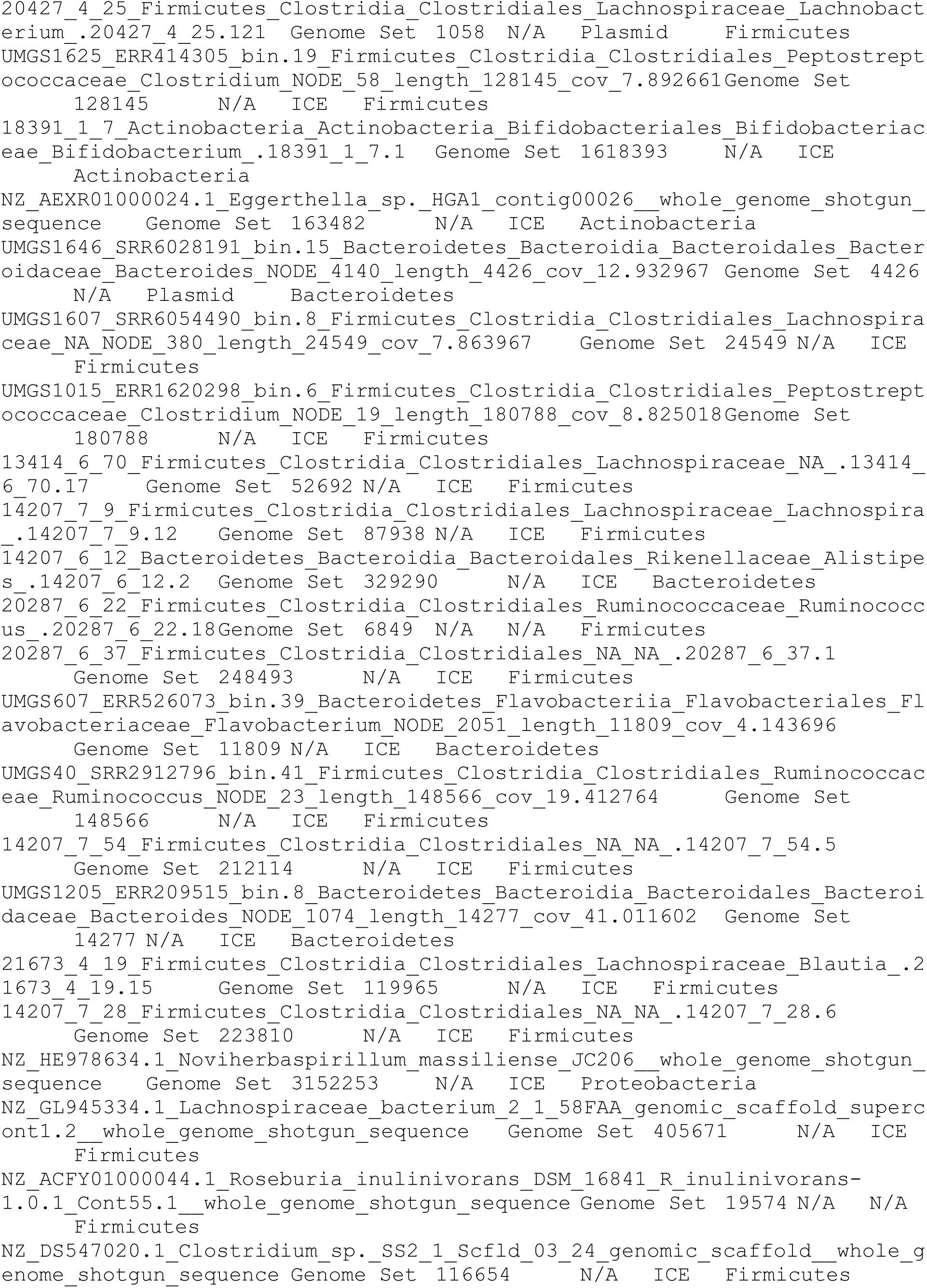
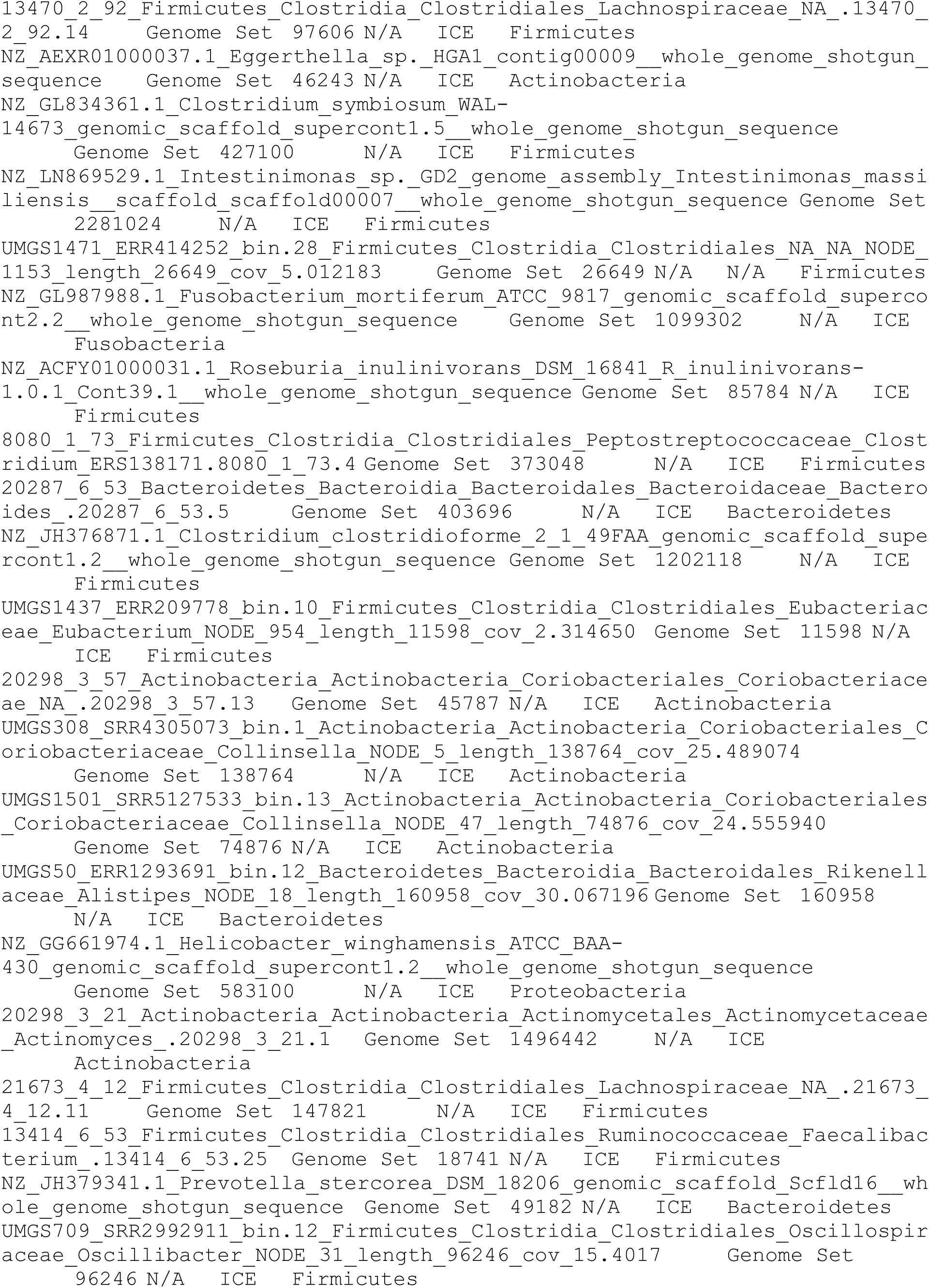
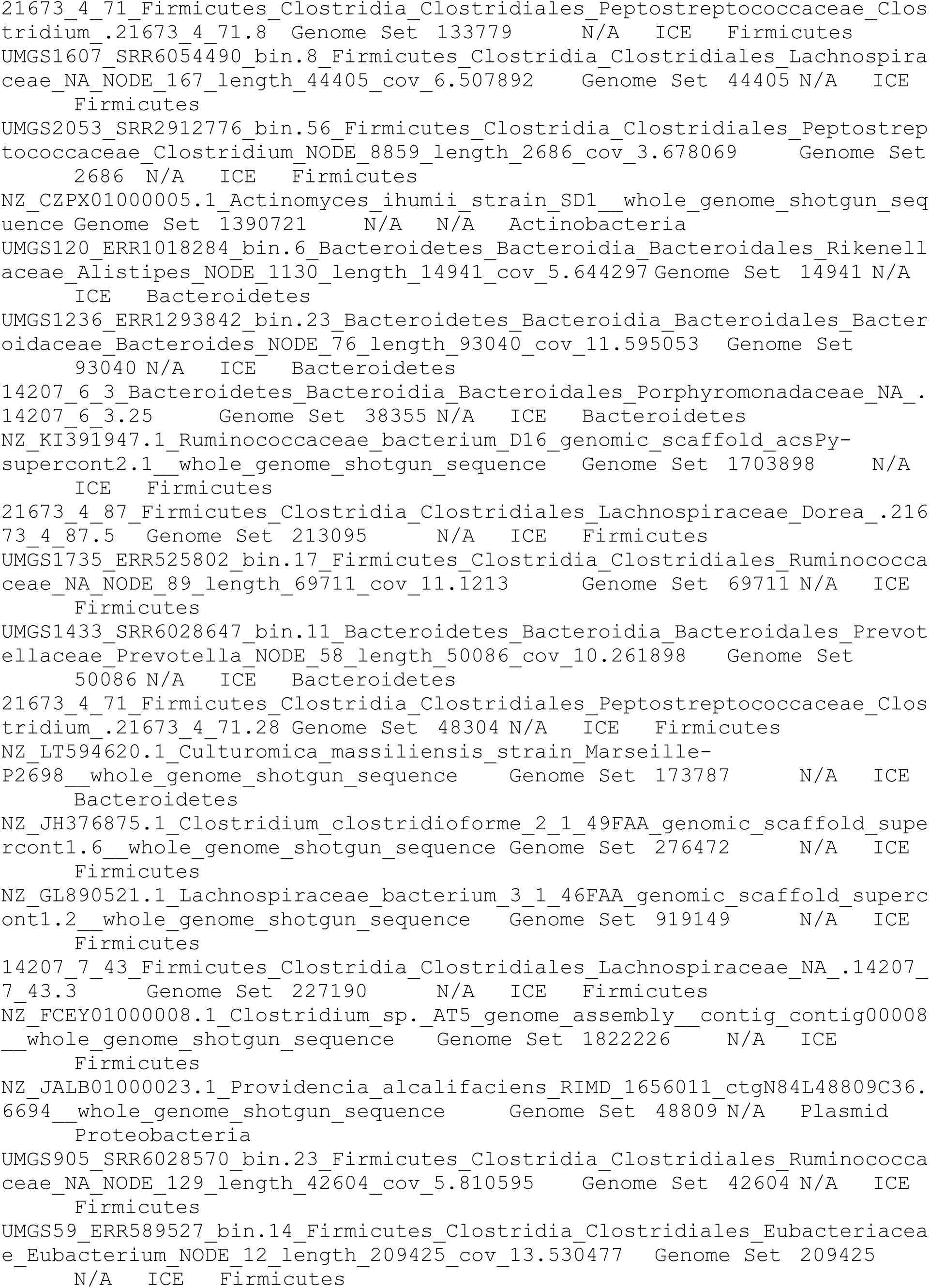
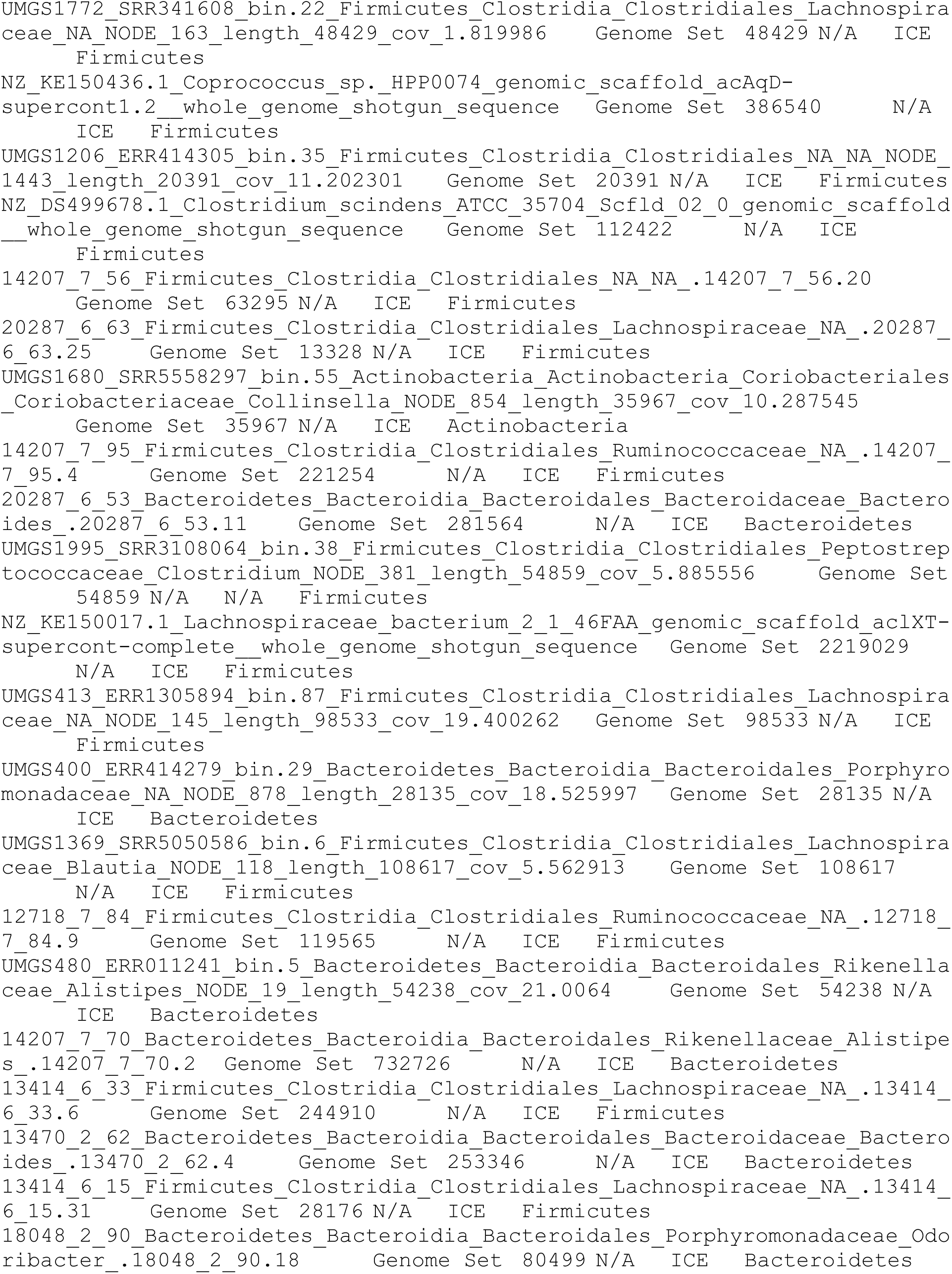
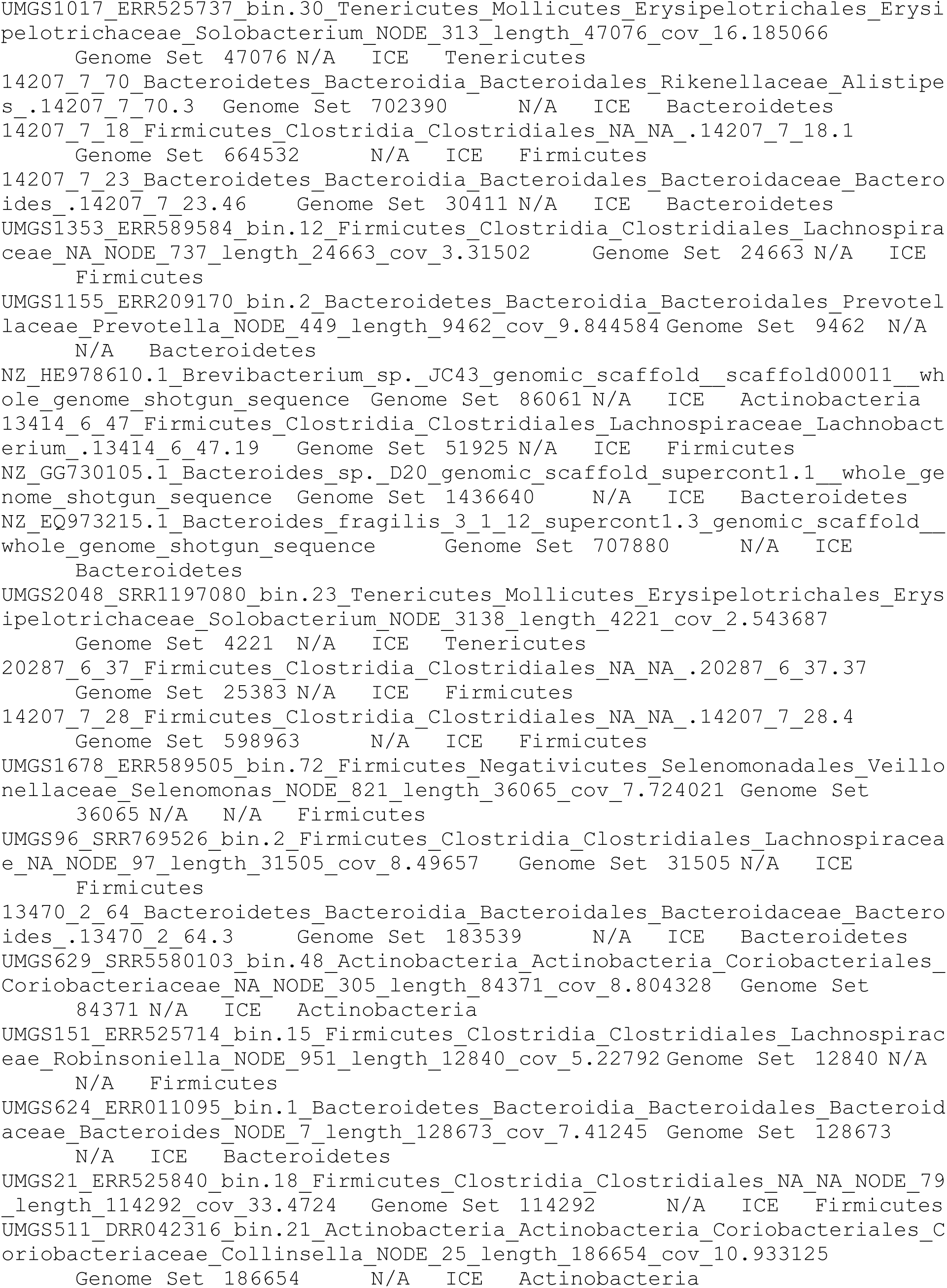
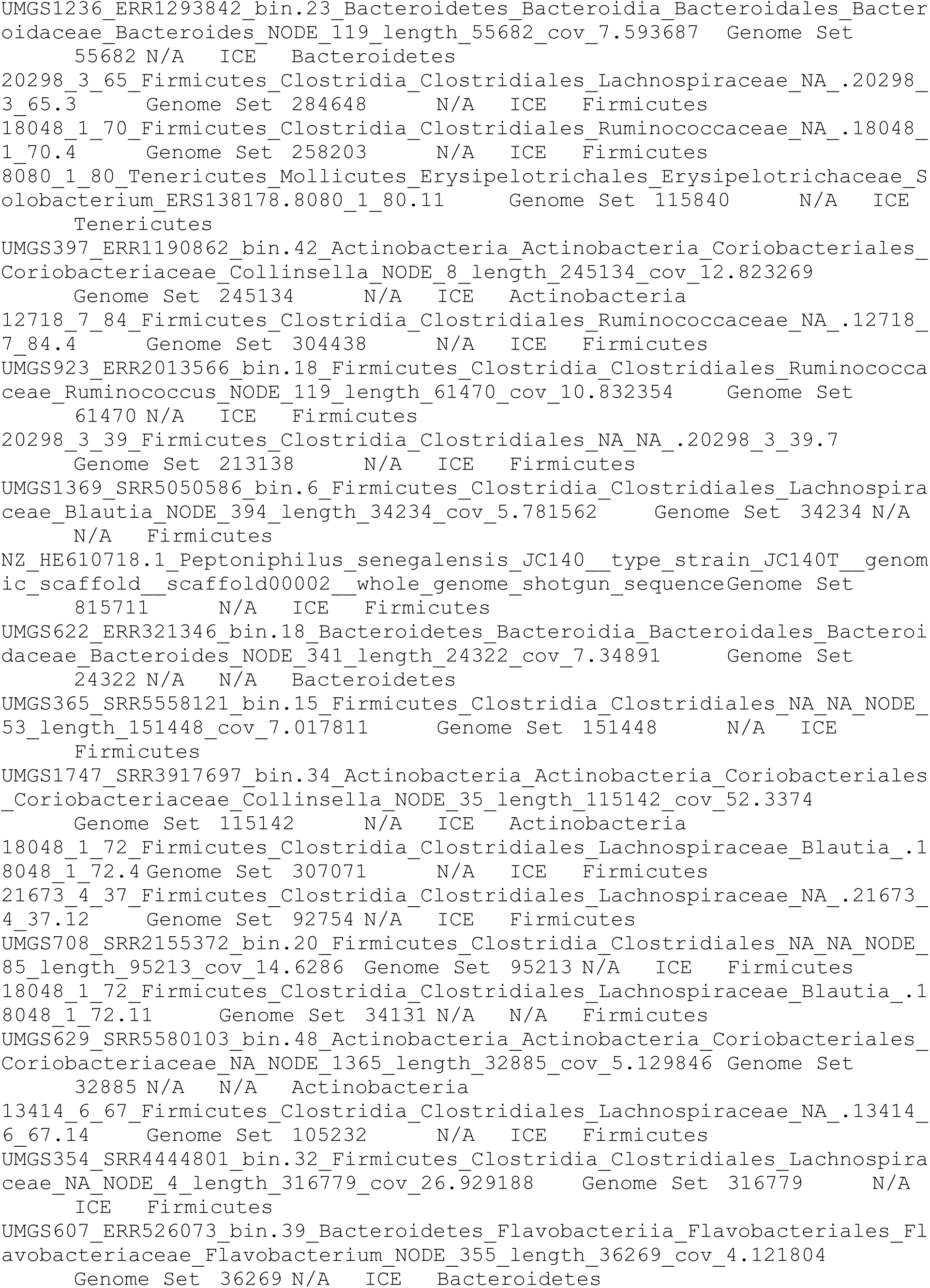
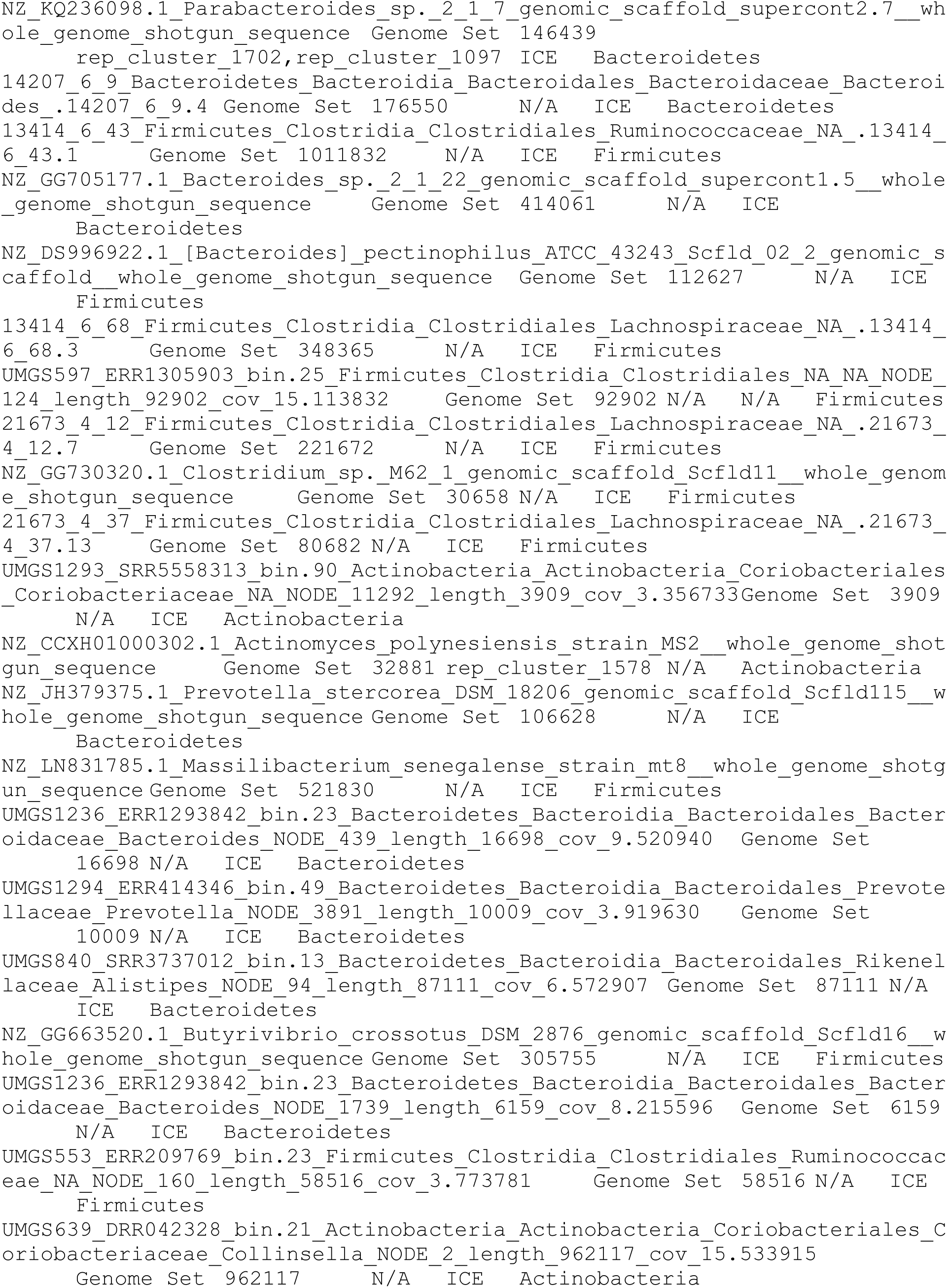
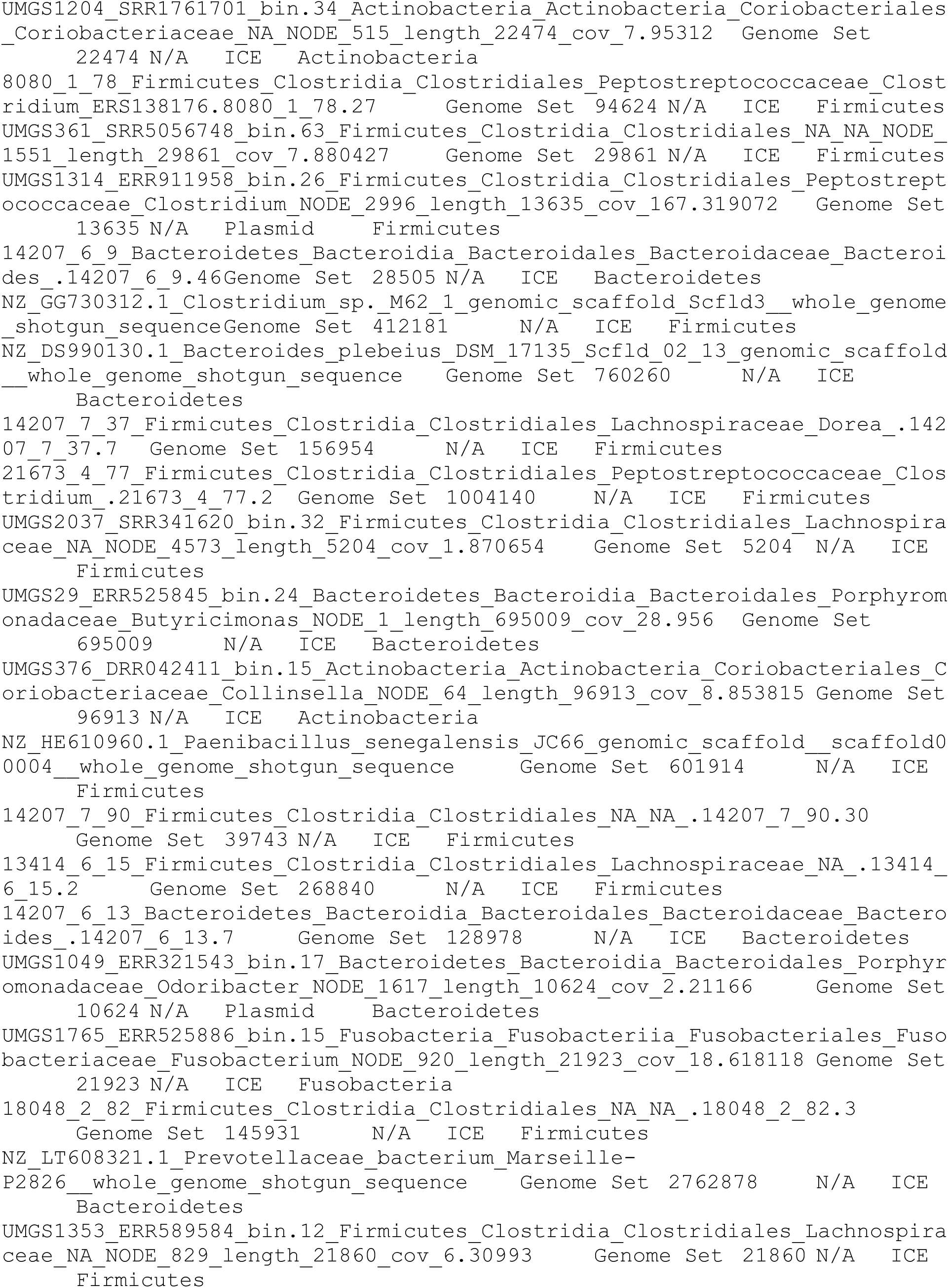
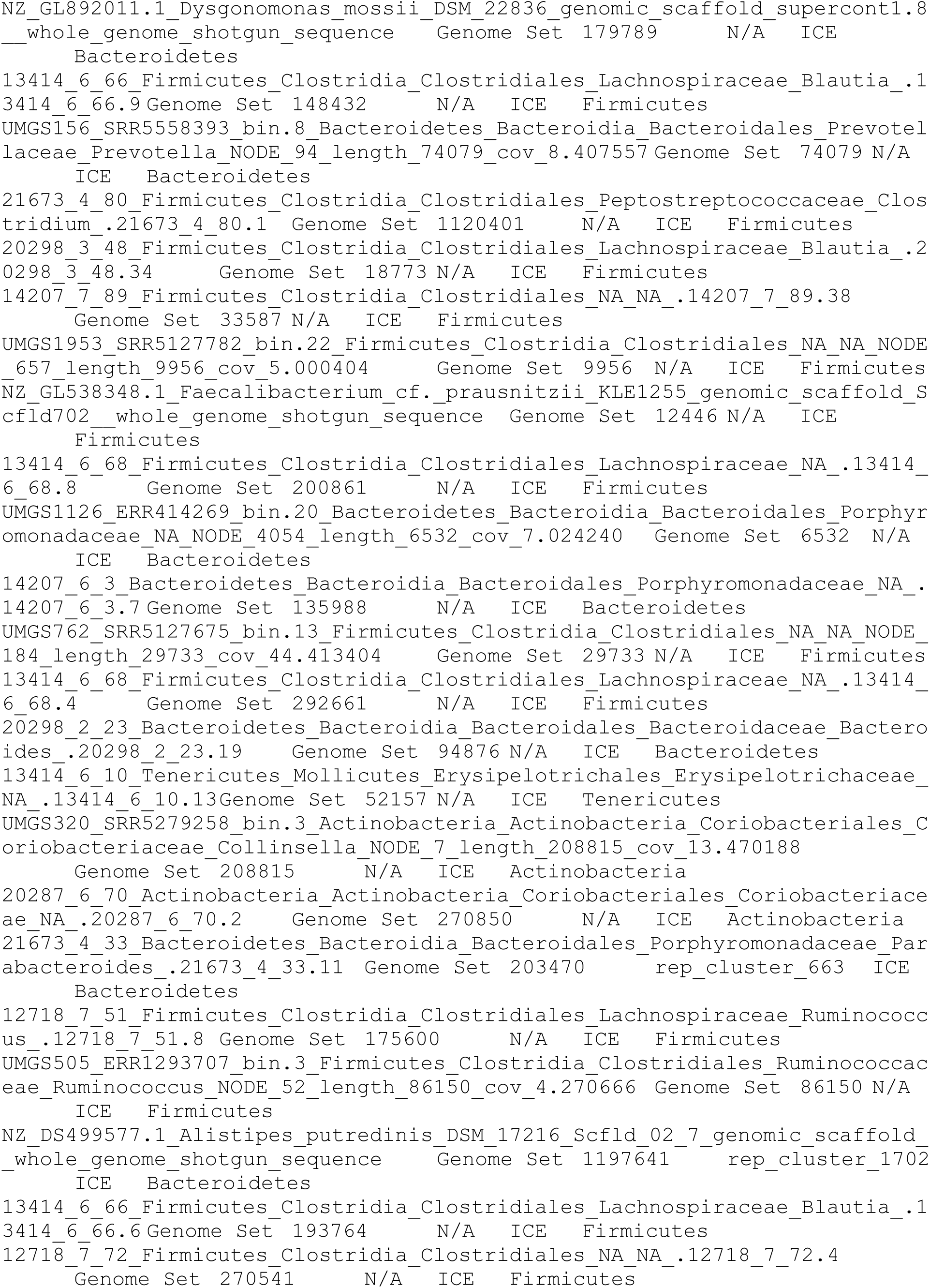
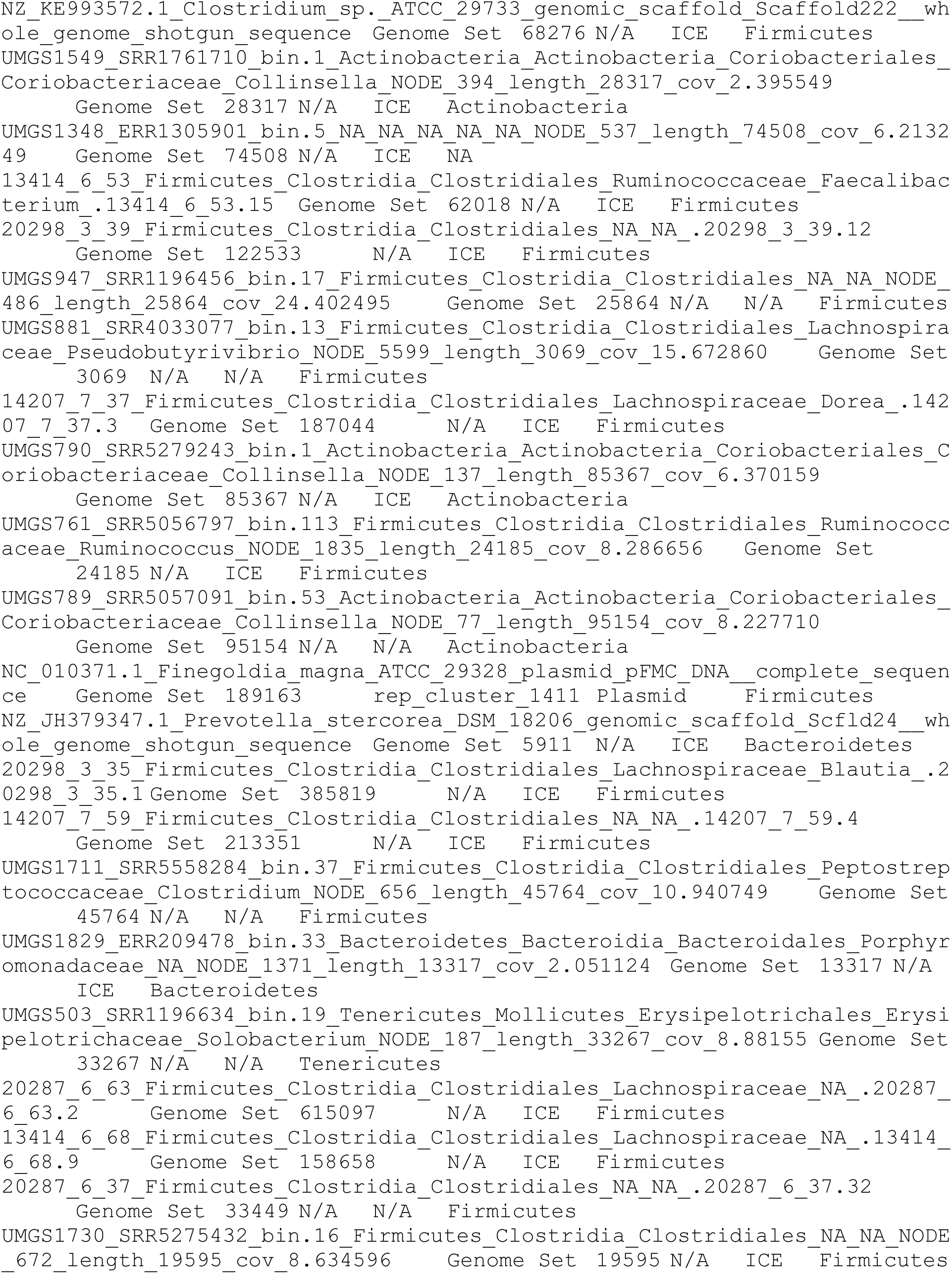
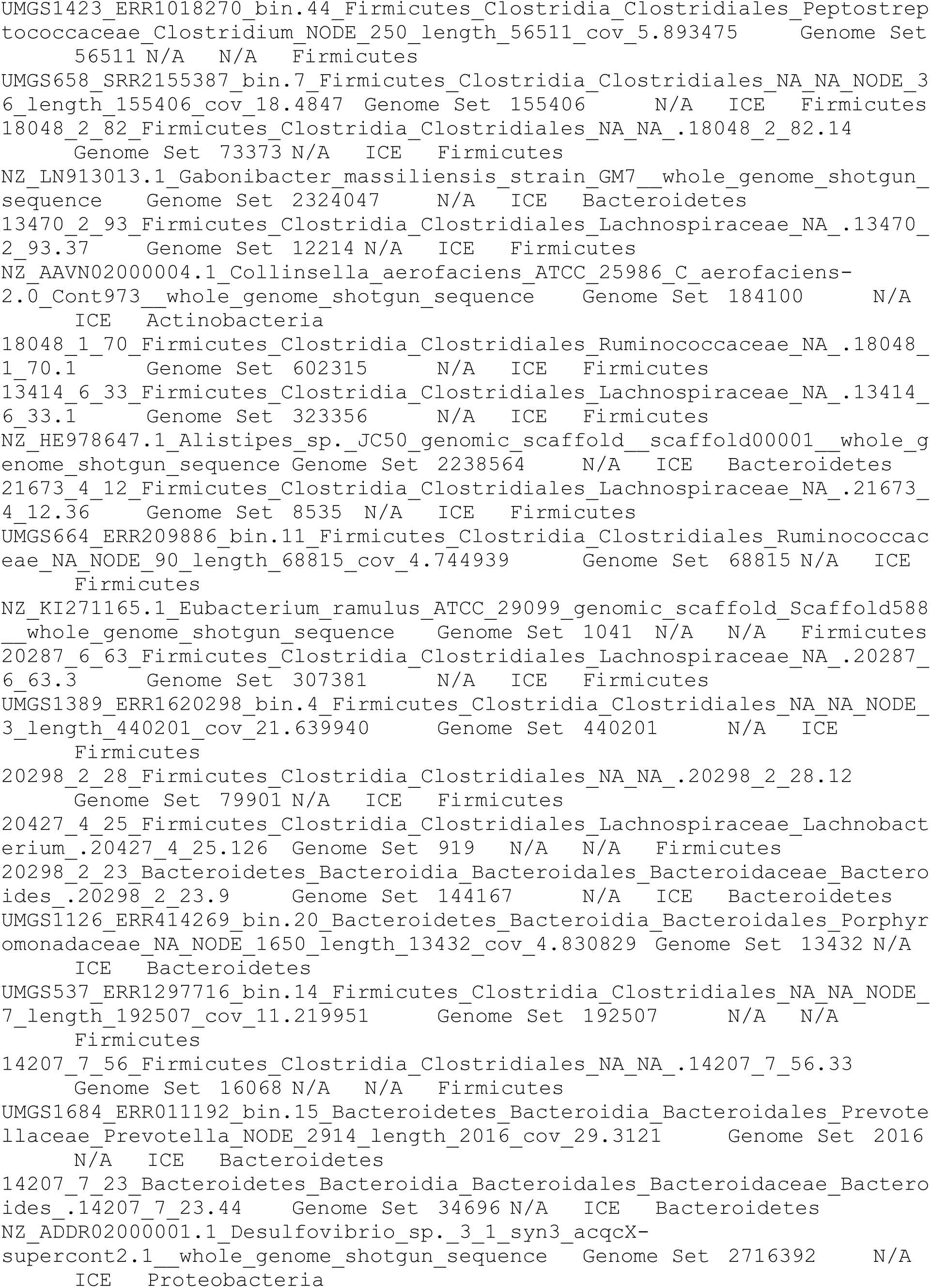
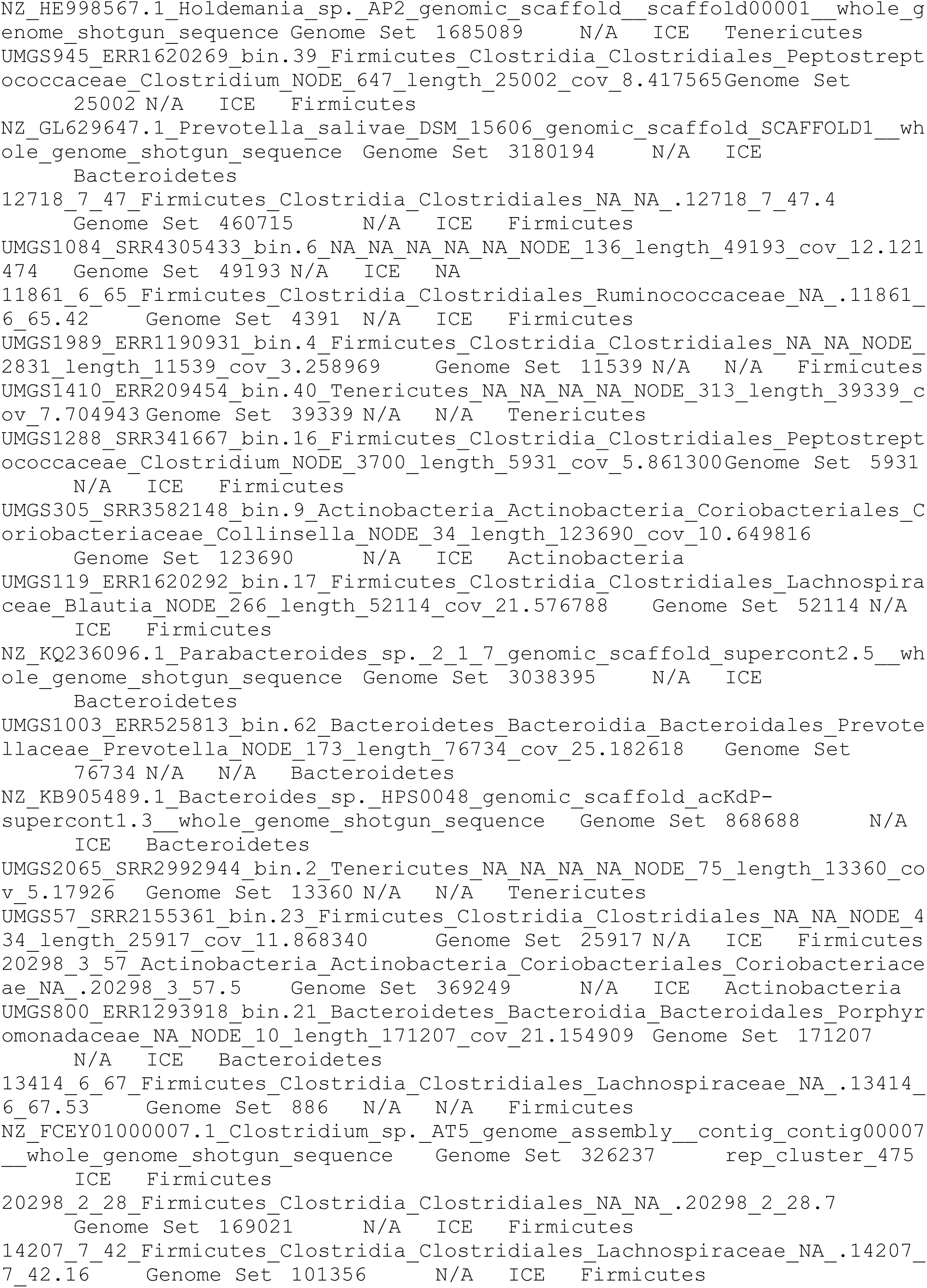
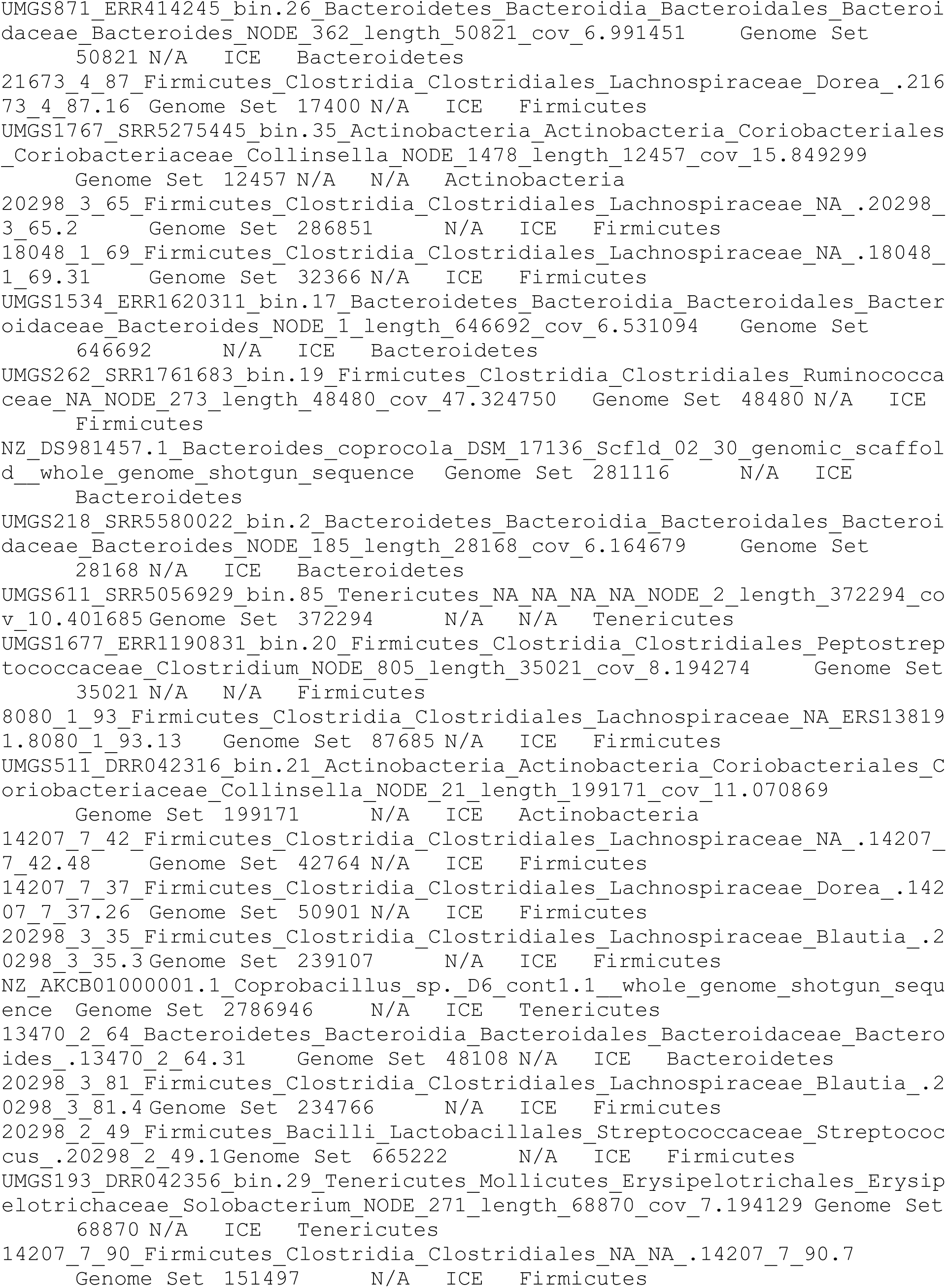
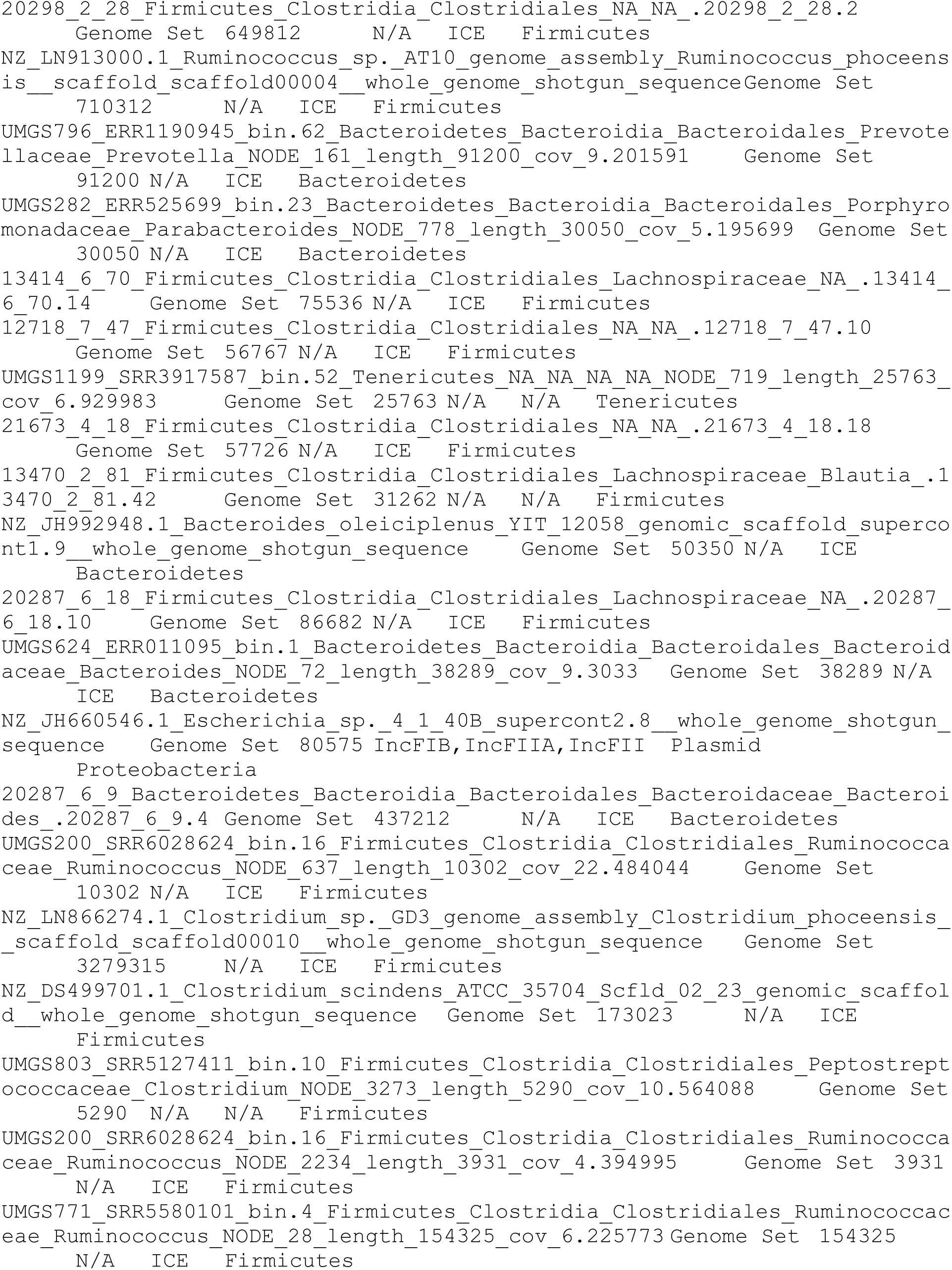
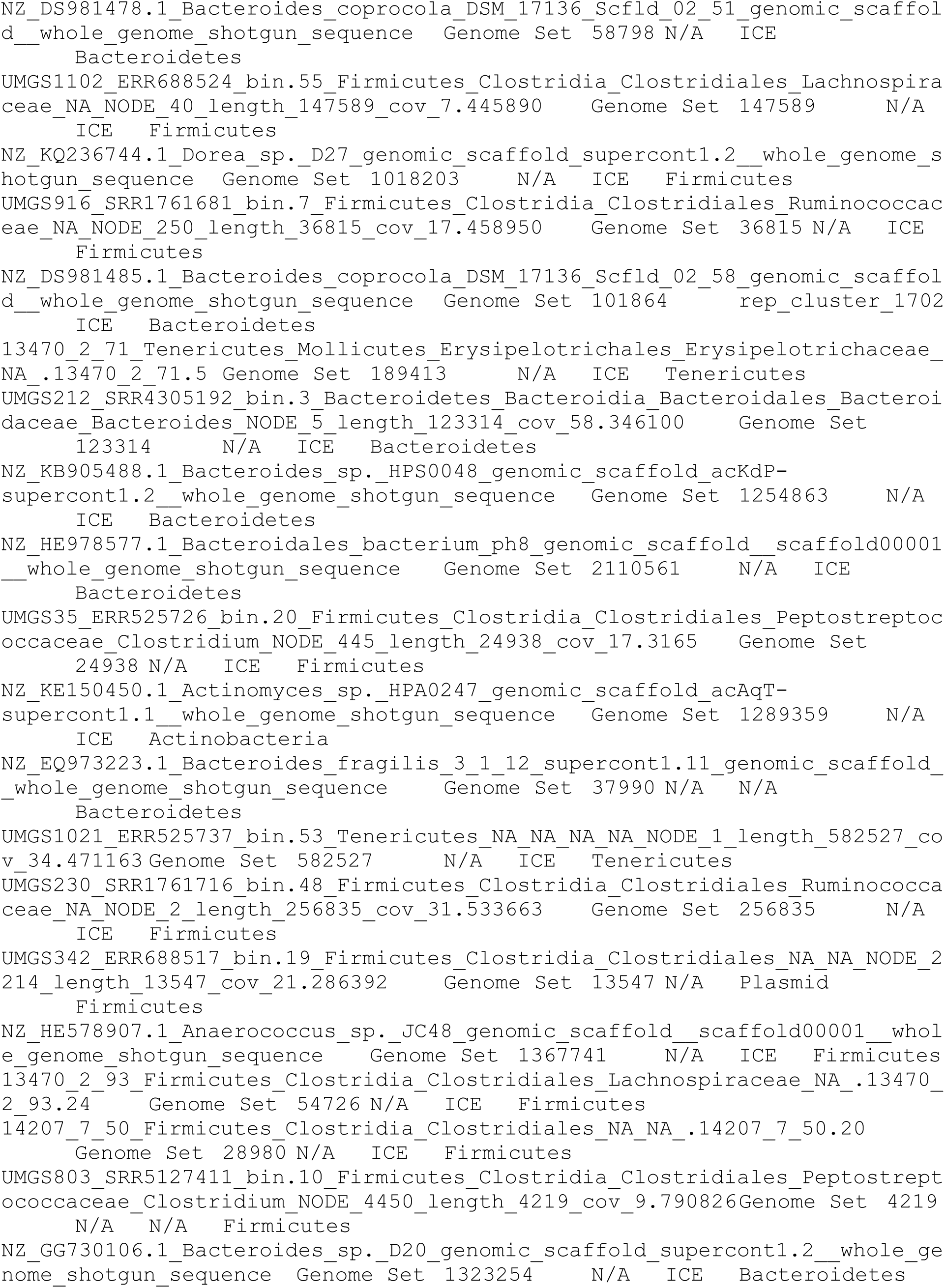
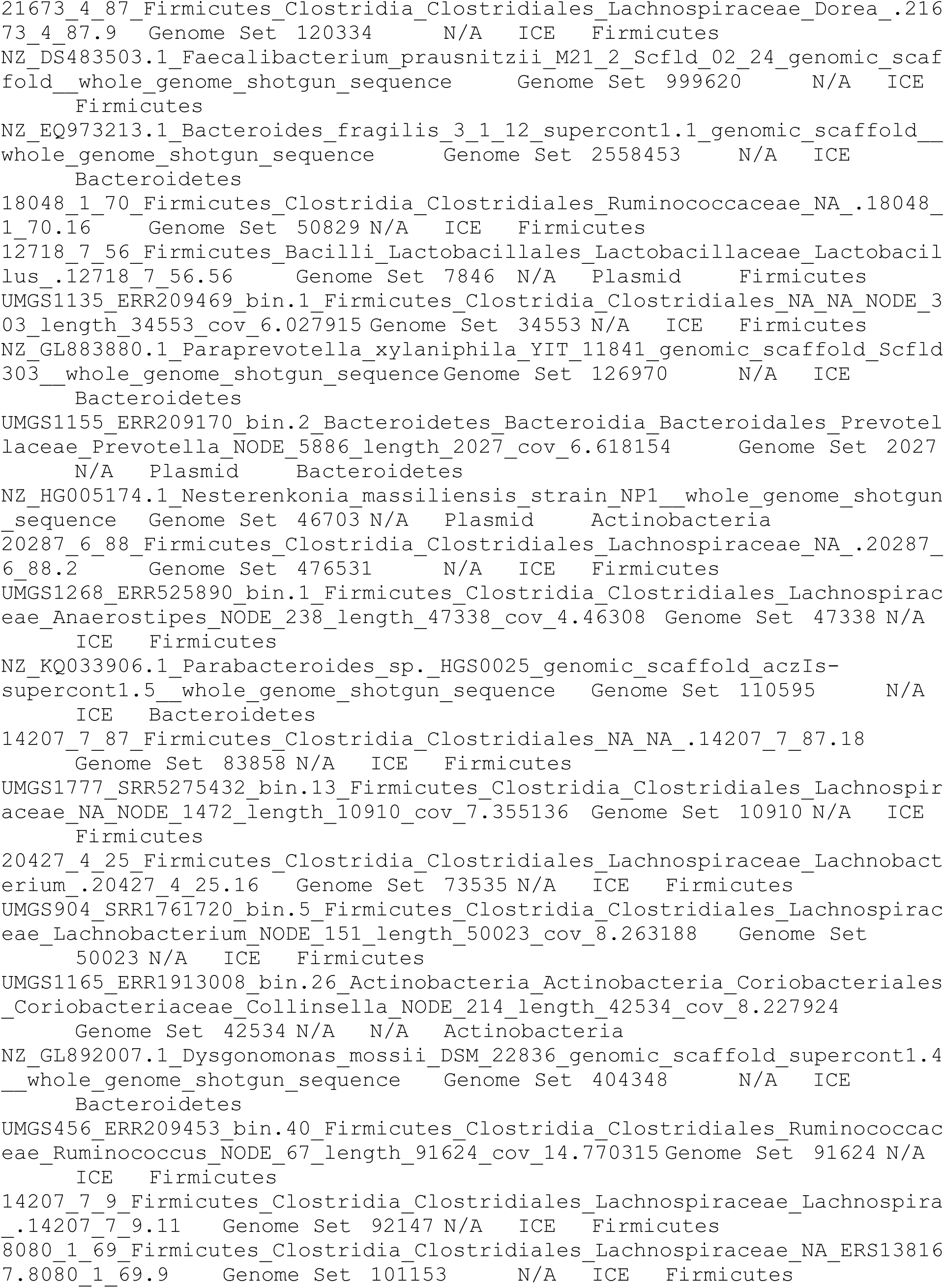
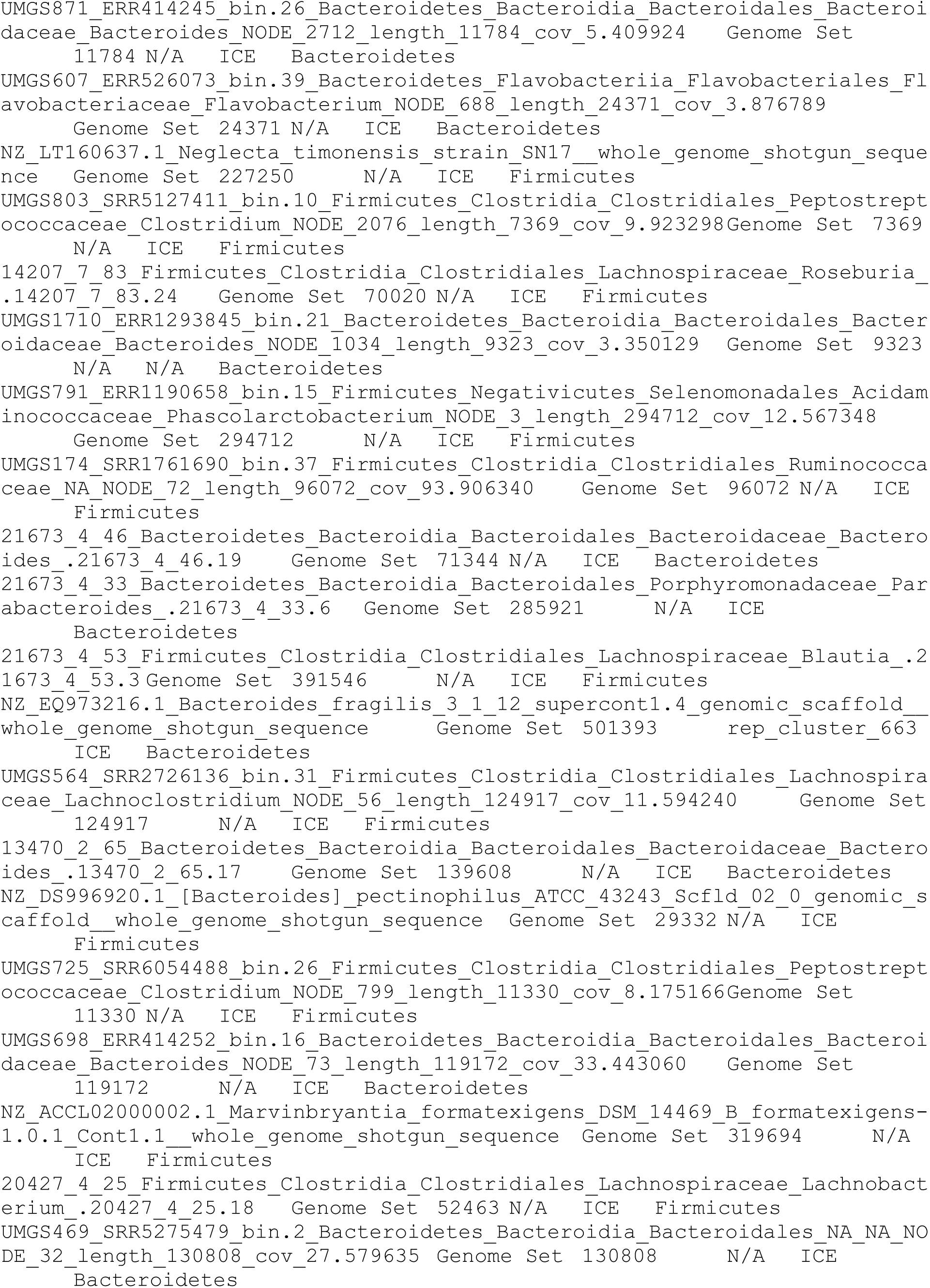
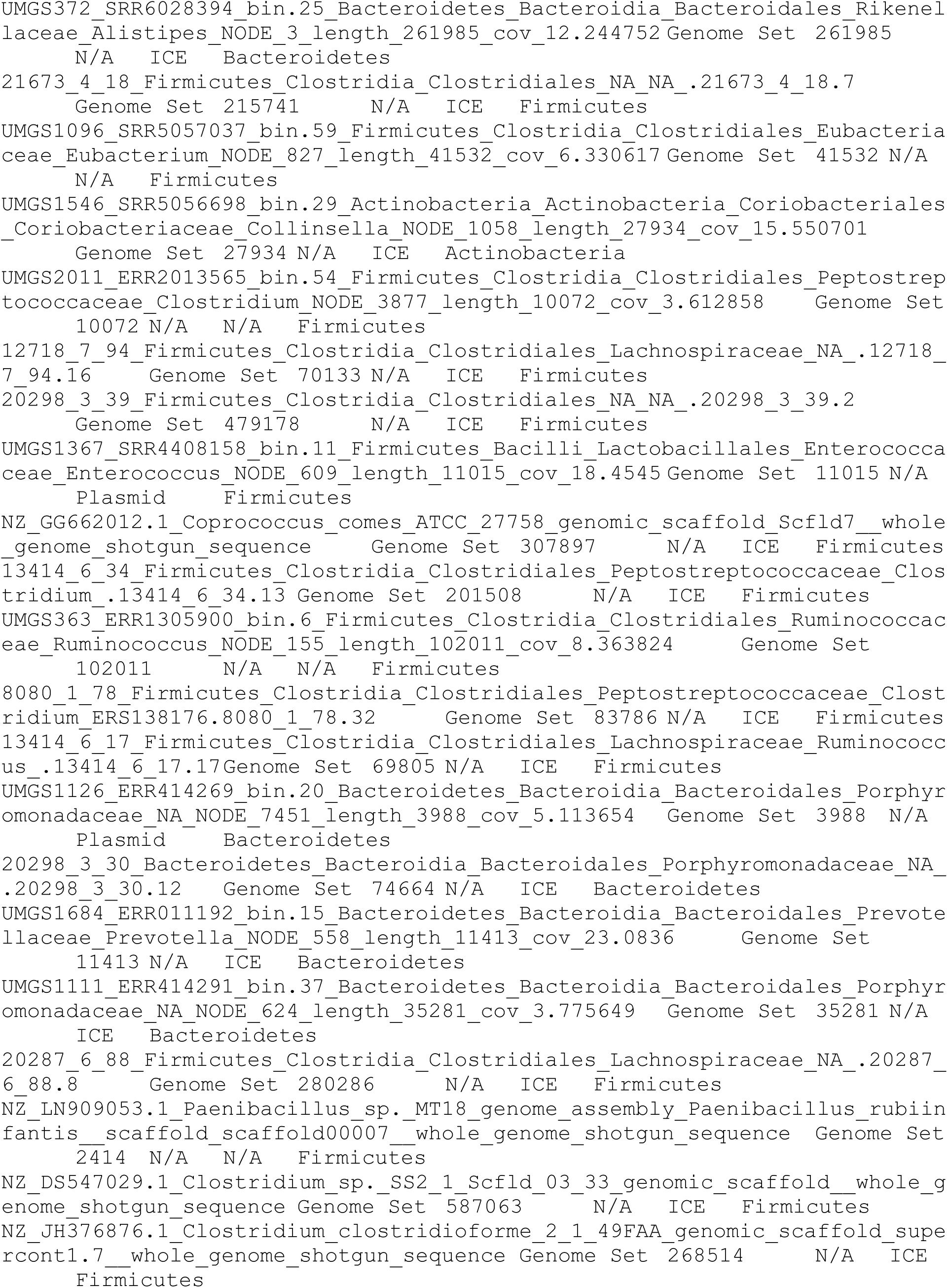
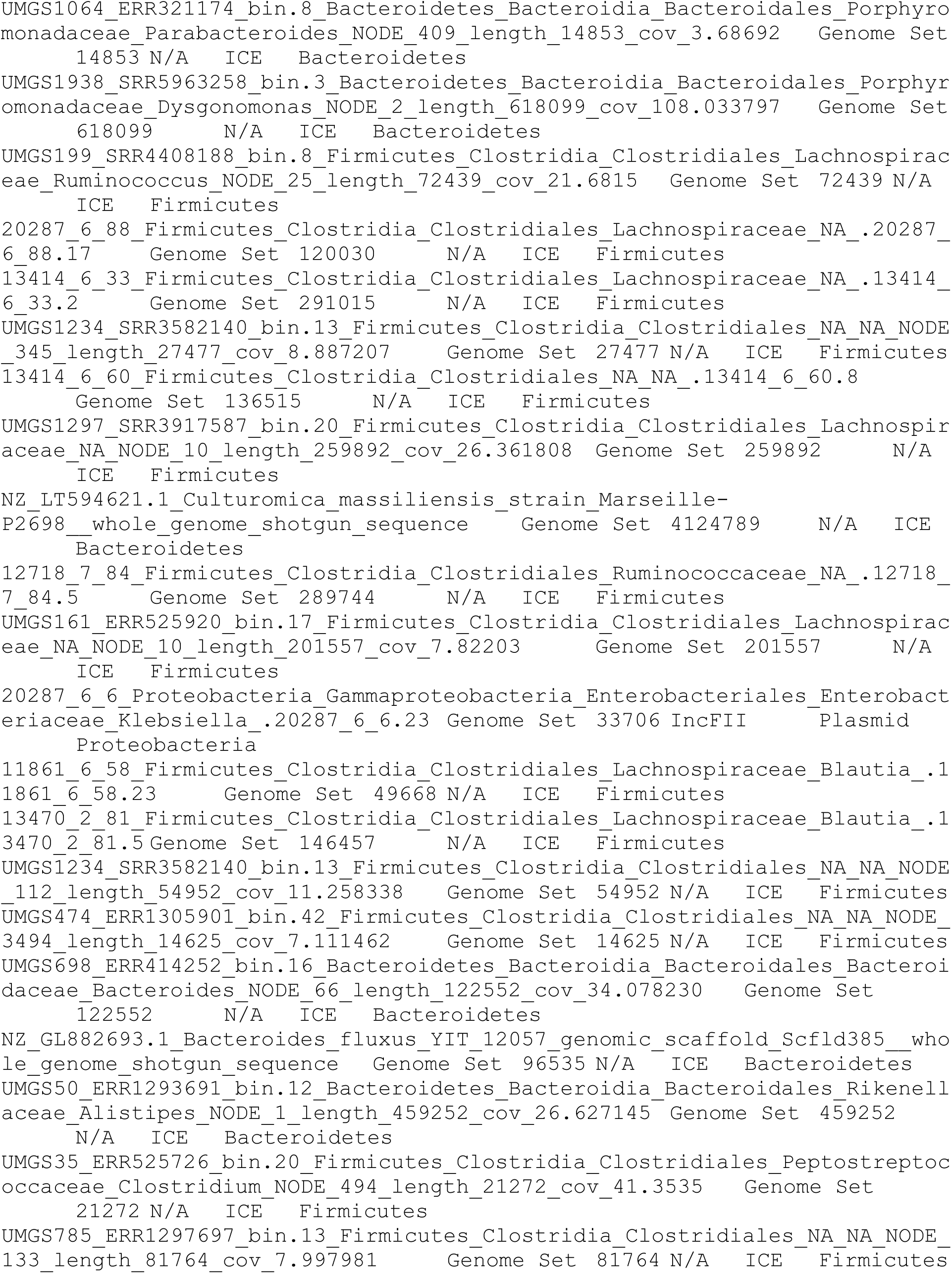
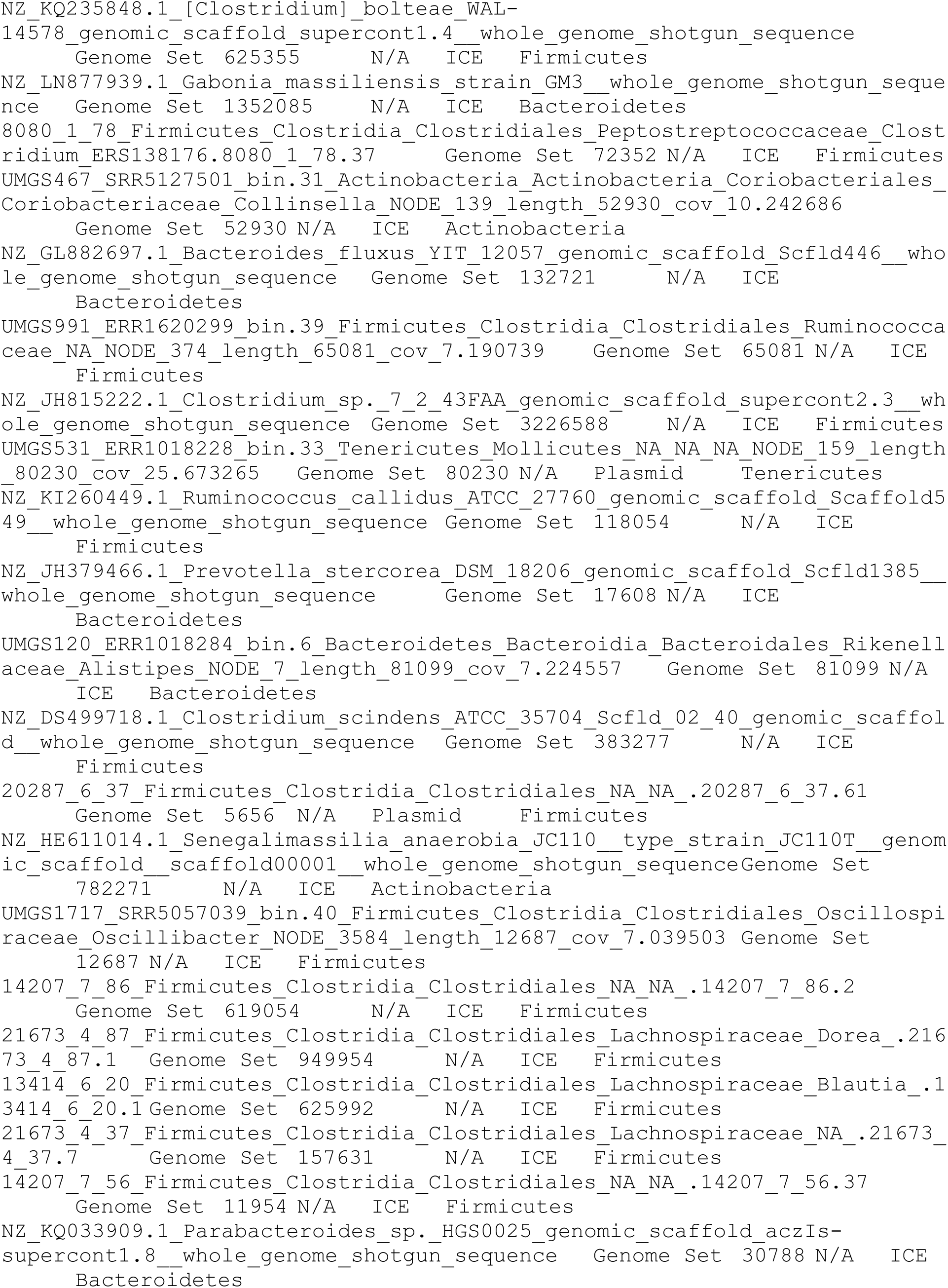
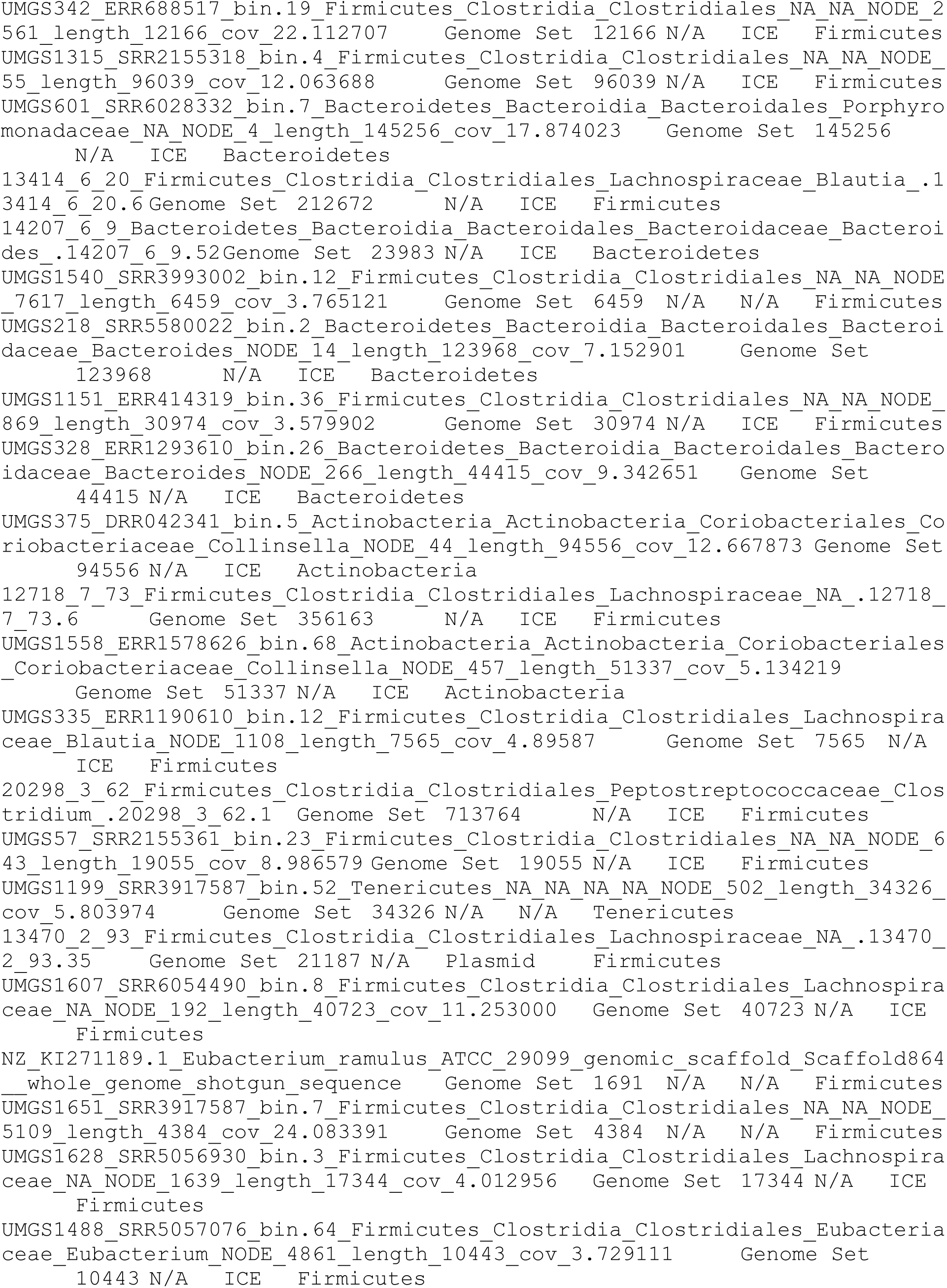
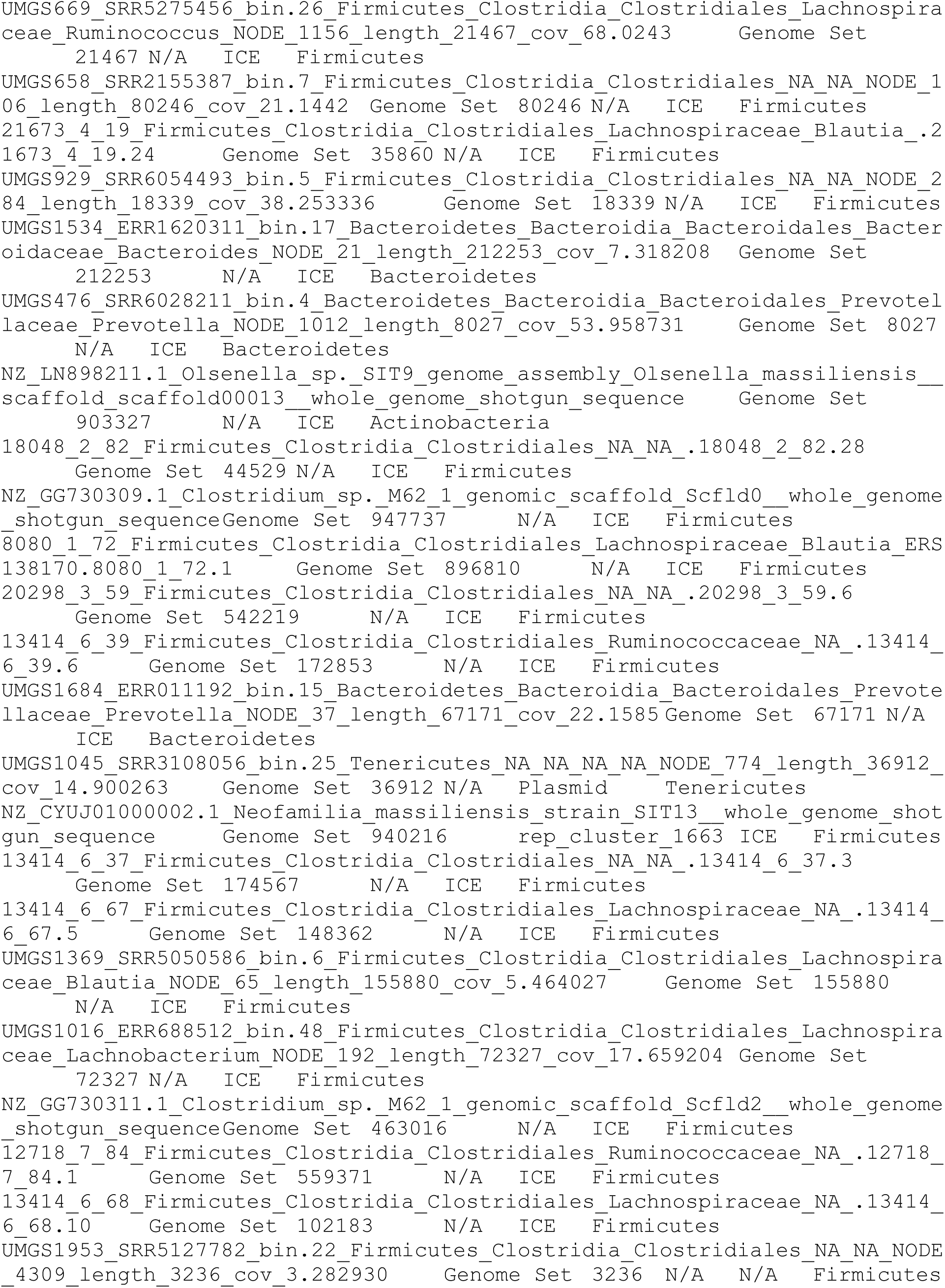
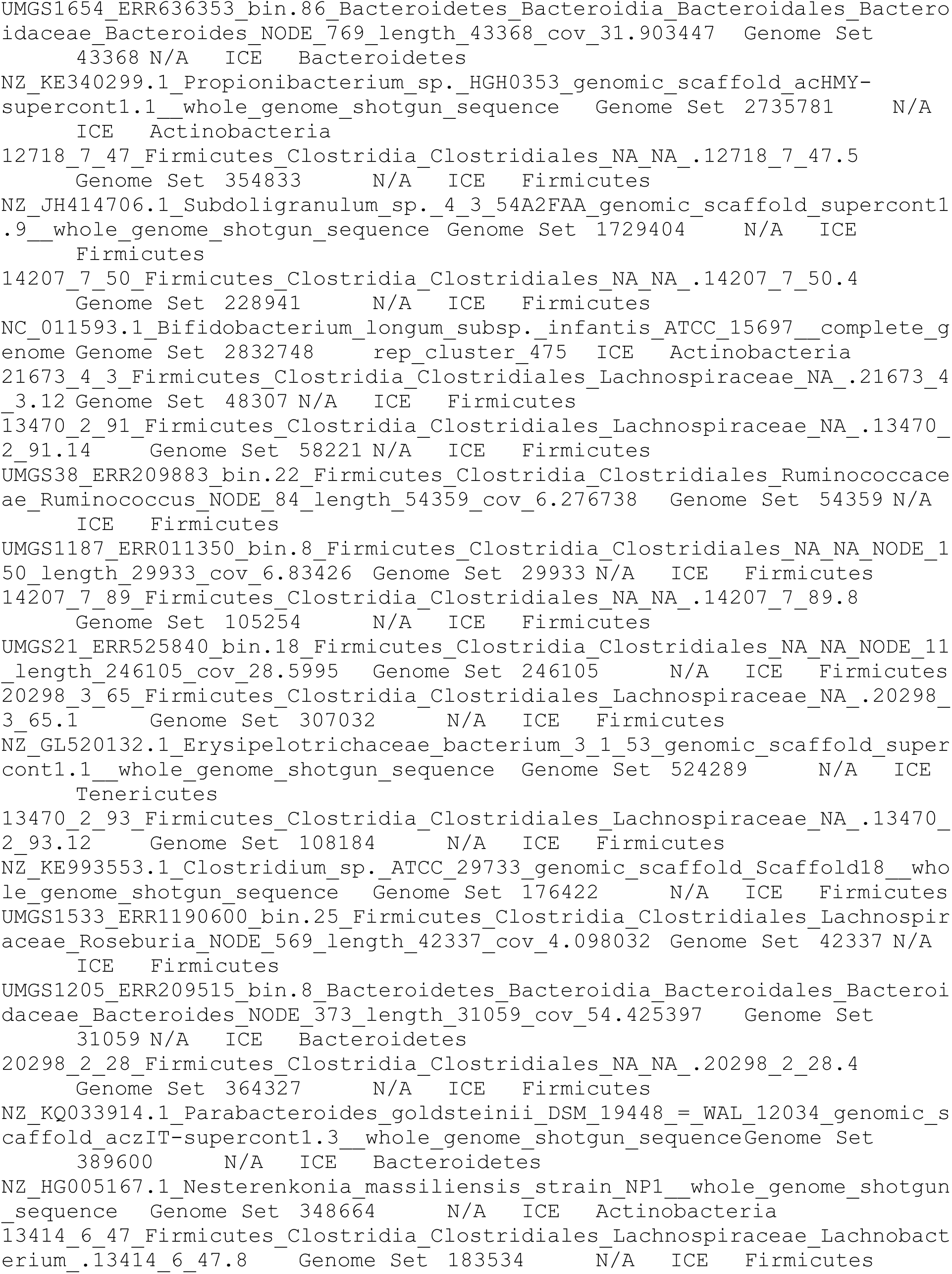
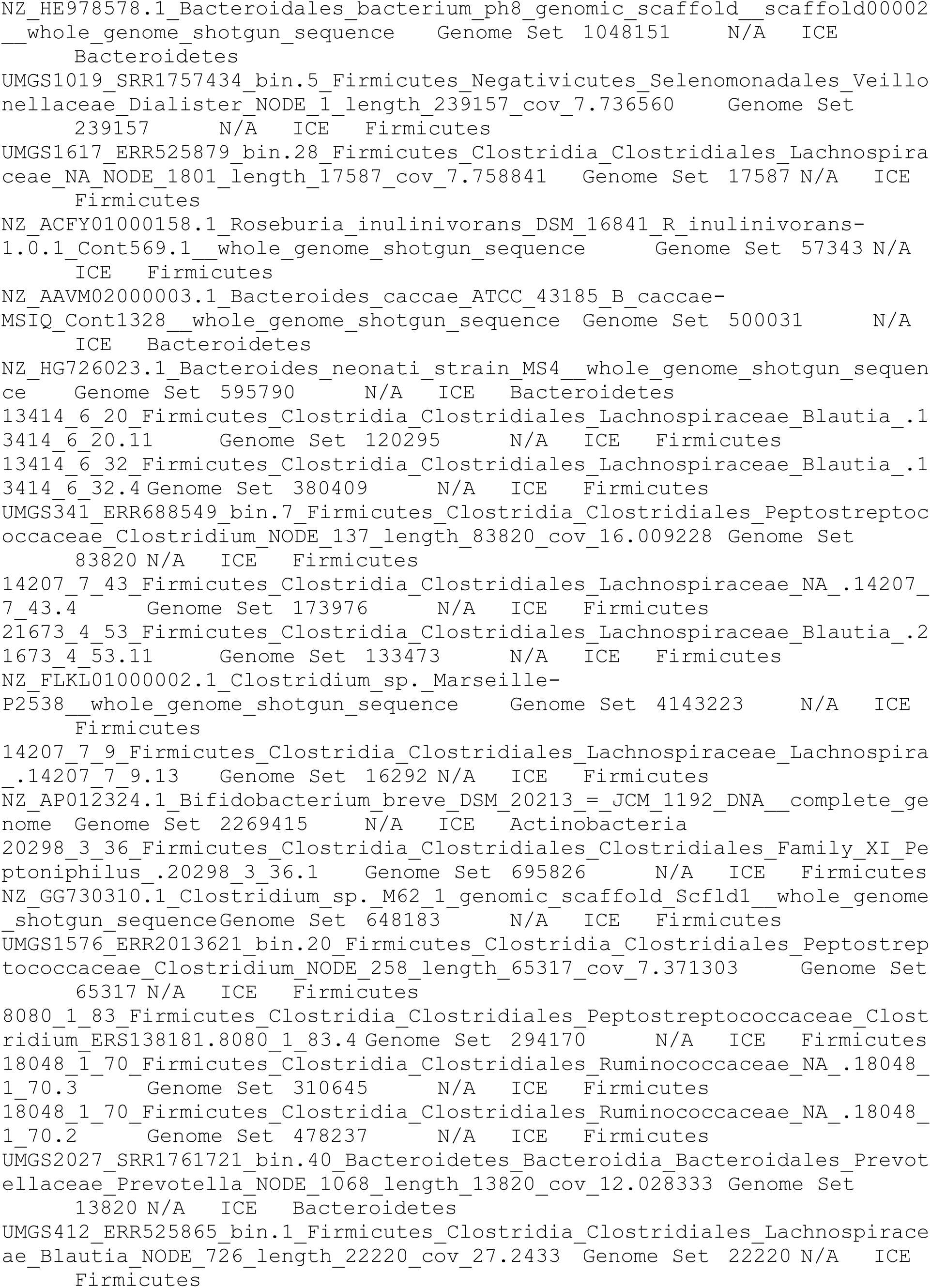
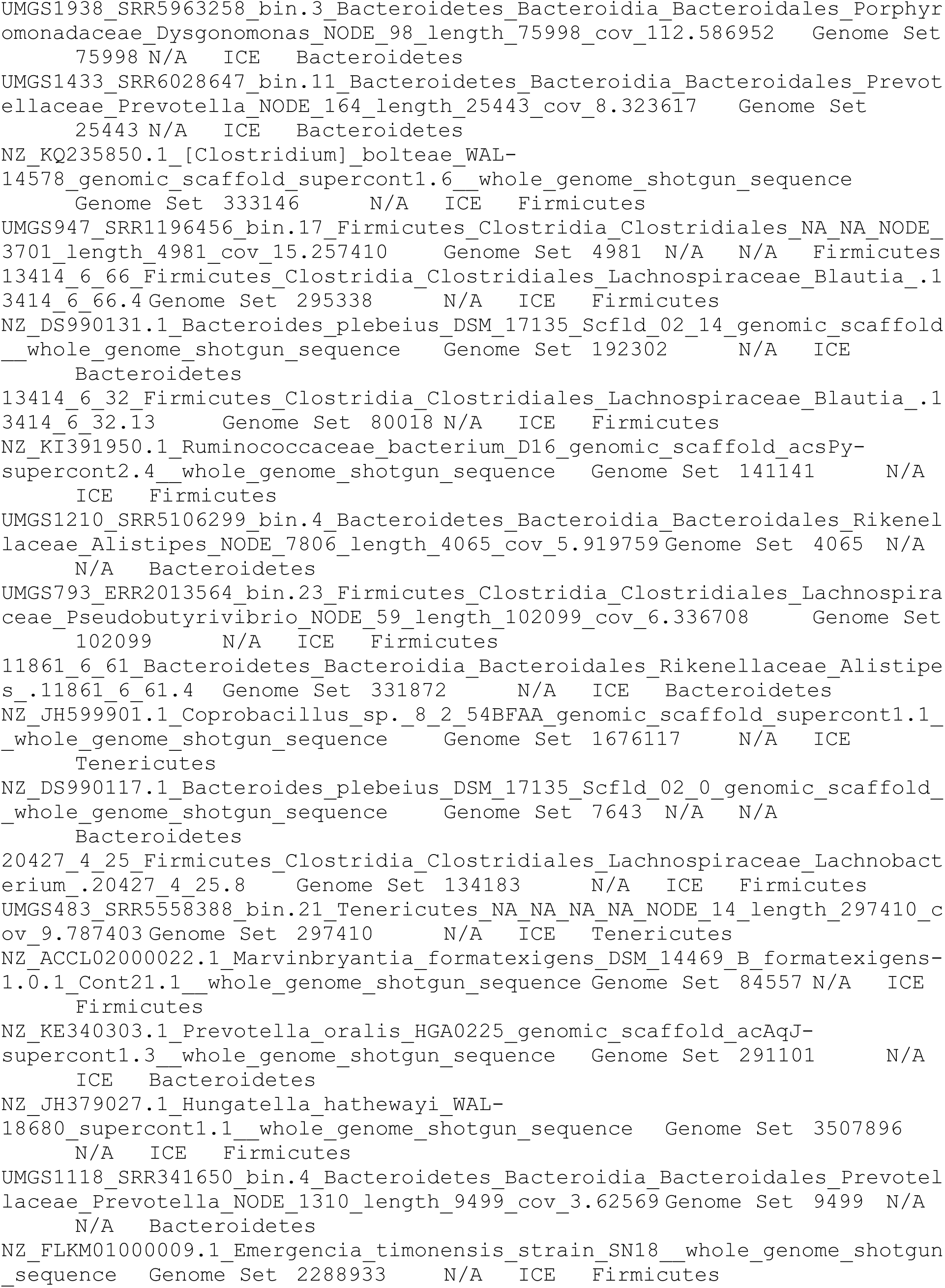
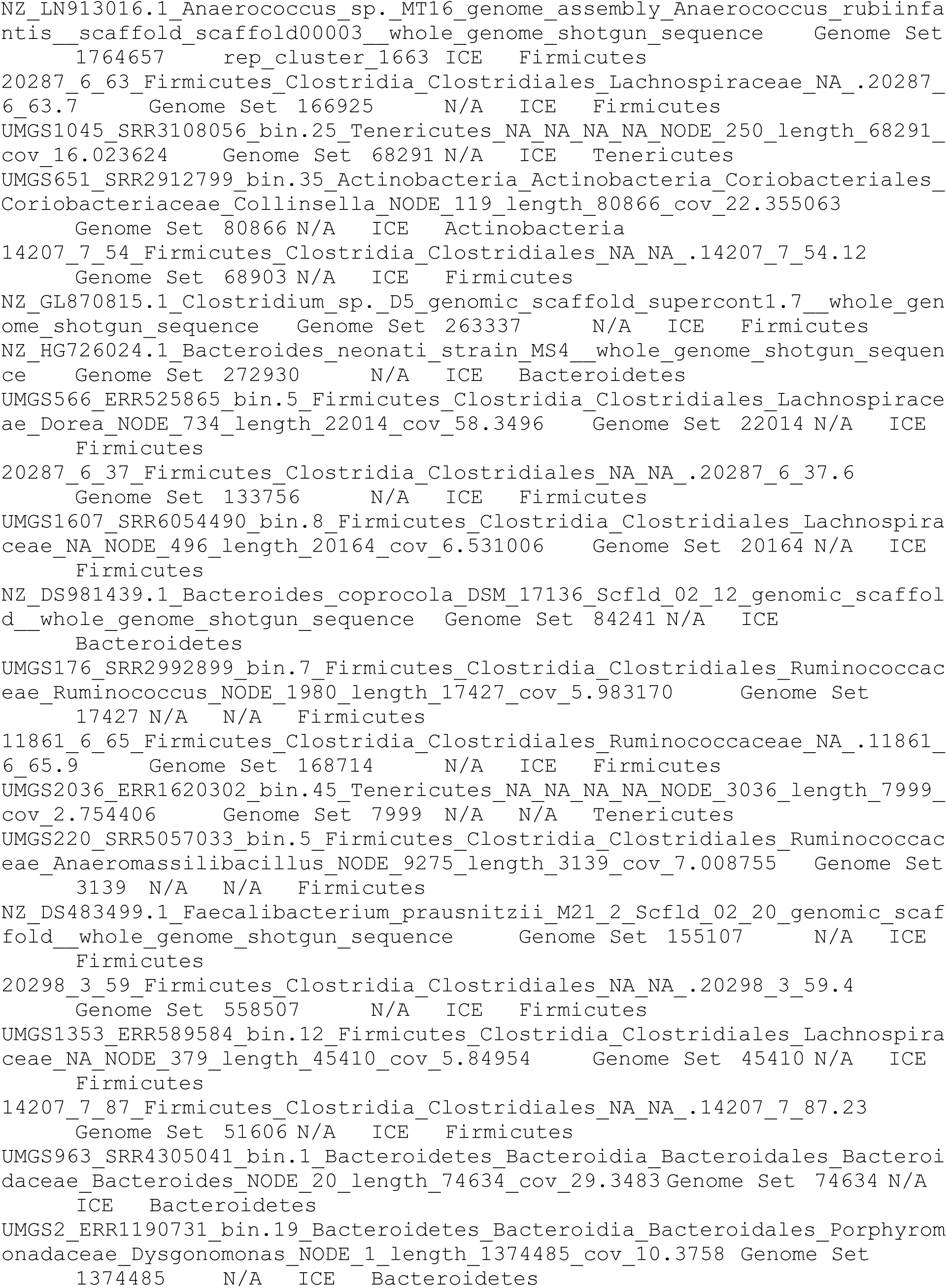
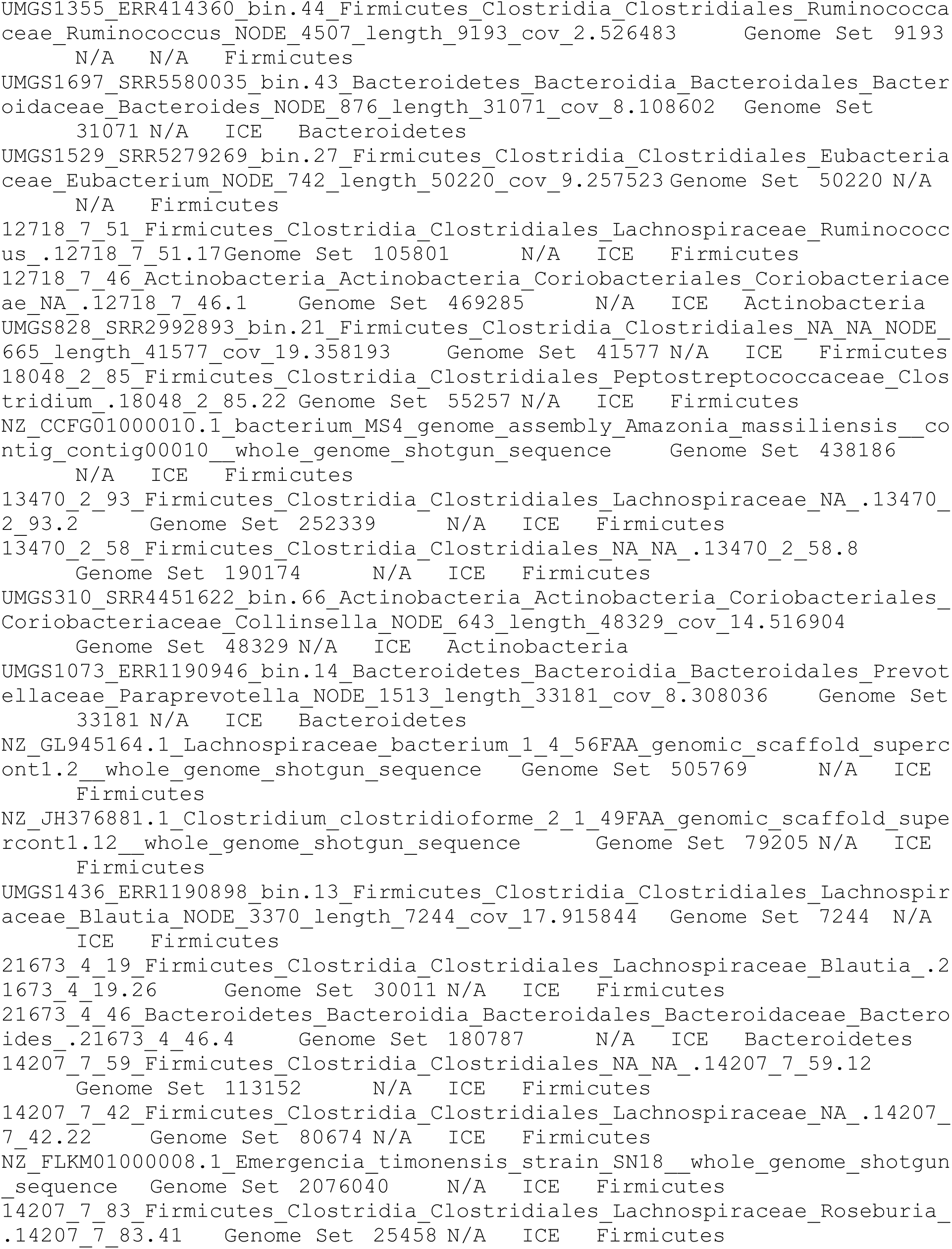

